# An atlas of the aging glycoproteome reveals how glycosylation modulates autoantibody binding in the brain

**DOI:** 10.64898/2026.10.28.762617

**Authors:** Michael Schoof, Johannes Reitenbach, Kevin Abraham, George Tilton-Low, Danwei Wu, Seyone Chithrananda, Yarden Freund, Divya Channappa, Lauren E. Pepi, Jeffrey E. Dunn, Marius Wernig, May H. Han, Tal Iram, Erin H. Seeley, Ian H. Guldner, Carolyn R. Bertozzi, Tony Wyss-Coray

## Abstract

Protein N-glycosylation generates substantial molecular diversity, yet how glycan composition varies throughout aging remains poorly understood. We mapped the N-glycoproteome of brain, liver, spleen, kidney, and plasma across nine ages spanning the mouse lifespan, identifying 79,517 unique N-glycopeptides at 9,217 glycosites on 4,397 glycoproteins, with matched proteomic trajectories (glycoproteinatlas.org). Organ-specific differences were larger than age-related changes, but aging still remodeled numerous glycosites. Brain aging increased short paucimannose glycans, driven partly by increased abundance of hexosaminidase B (HEXB). In white matter, aging increased paucimannosylation of myelin oligodendrocyte glycoprotein (MOG), a major autoimmune target. We identified MOG-reactive autoantibodies in aged individuals and patients with MOG antibody-associated disease (MOGAD) and showed that MOG glycosylation modulates autoantibody binding. Glycosylation changes associated with normal aging were also observed after experimental autoimmune encephalomyelitis (EAE), a model of autoimmune neuroinflammation. These findings reveal N-glycosylation as both a marker of age and a regulator of disease-relevant molecular interactions.

## Introduction

Aging is the greatest risk factor for neurodegenerative disorders^1-3^. At the molecular level, aging drives a cascade of detrimental changes in the brain that collectively impair functions ranging from higher-order processes like learning and memory to basic sensory and motor coordination^4,5^. These changes have been frequently assessed at the RNA level and, increasingly, at the protein level as well^6-8^. The protein space is extremely vast. Considering splice variants, ∼100,000 unique protein sequences can be generated from the ∼20,000 human genes, and these protein variants can have distinct functions, localization patterns, and disease implications^9-11^. The vast array of available post-translational modifications (e.g., glycosylation, phosphorylation, acetylation, and lipidation) further expands this complexity, with upward of 1 million distinct proteoforms existing within each cell when these modifications are considered^12,13^. Proteoform diversity is a critical but underexplored dimension of aging biology, and glycosylation is among the most pervasive and structurally diverse modifications that generate it^14^.

All living cells are coated with a complex assortment of glycosylated proteins (plus lipids and RNAs)^15,16^. Hundreds of processing enzymes across multiple organelles elaborate and trim these diverse glycans on varied scaffolds, and an even greater number of proteins play roles in necessary biosynthesis and recycling processes^17^. In mammals, over 2% of the genome is dedicated to these processes and hundreds of congenital disorders of glycosylation have been identified wherein this process is dysregulated^18,19^. Glycoproteins can exist in numerous glycoforms that vary in glycosite occupancy and glycan structure and composition. These glycoform patterns are known to be cell-type dependent and dynamic in disease settings, but to date, glycosylation dynamics have been studied mostly in the context of cancer and viral immune evasion^20-23^. Cancer cells often increase cell-surface sialoglycan density in response to pressures of immune surveillance. Viruses also leverage dense surface glycosylation as a ‘glycan shield’, promoting immune evasion. Recent work has started to address how the blood-brain barrier vascular glycocalyx responds to normal aging and neuroinflammatory disease, and there is emerging interest in the role of dysregulated glycosylation in neurodegenerative diseases such as Alzheimer’s disease (AD)^24-29^. Yet little has been done to systematically examine at a site-specific level and across multiple tissues how N-glycosylation of proteins changes during normal aging. Distinct glycans can reshape interaction surfaces and thereby influence molecular recognition, including binding of autoantibodies which are known to accumulate with aging^30^. Recent technical advancements that have made proteome-wide glycosylation mapping possible, as well as our growing appreciation of the importance of glycans in human biology, have motivated us to explore the role that N-glycosylation plays in aging^31-41^.

## Results

### A multi-organ atlas profiles N-glycosylation and protein abundance across the mouse lifespan

Seeking to explore how N-glycosylation varies during aging across organs, we assembled a cohort of mice (n = 6 males per age; 1, 3, 6, 9, 12, 16, 19, 22, and 28 months; all C57BL/6JN strain) and collected the brain, liver, spleen, kidney, and plasma from all animals **(Figure 1A)** for input into high-throughput N-glycoproteomics and proteomics workflows optimized for 96-well sample processing **(Figure 1B)**. These approaches generated robust datasets with consistent overall detection depth within each organ **(Figure S1)**. After post-processing and quality filtering across all 270 samples **(Figure S2)**, we identified 3,577,876 glycopeptide-spectrum matches (glycoPSMs), representing 79,517 unique N-glycopeptides and 9,217 glycosites located on 4,397 glycoproteins **(Figure 1C)**^31,36,42,43^. Glycosite detection varied substantially between organs, with 3,986 detected in brain and 1,050 in plasma **(Figures 1F and 1H)**. Replicate samples from a given organ yielded similar numbers of glycoPSMs, but the specific glycosites detected and their sampling depths varied across samples **(Figure 1H; Figure S1D)**. We therefore established a tiering system, ranging from Tier 0 sites supported by fewer than five total glycoPSMs to progressively more deeply sampled sites in Tiers 1–7 **(Figure 1H; Figure S3D)**. Parallel proteomic profiling quantified 10,679 total proteins across at least one organ **(Figures 1D and 1G; Figures S4A, S4B, and S4E)**, including 4,015 proteins in the 4 solid tissues and 346 more in all 5 datasets. There was generally strong coverage of the detected glycoproteins in both datasets – outside of the plasma about two thirds of the proteins with a detected glycosite in a given organ had matching proteomics data **(Figure 1E)**. Consistent with known features of N-glycosylated proteins, detected glycoproteins were enriched for extracellular matrix, lysosomal, and plasma membrane localization and for structural features and domains including transmembrane regions, signal peptides, and Ig-like and EGF-like domains **(Figures S5 and S6)**. Together, these datasets constitute the first site-specific multi-organ atlas of N-glycosylation across the mouse lifespan and are publicly available at glycoproteinatlas.org.

**Figure 1.**
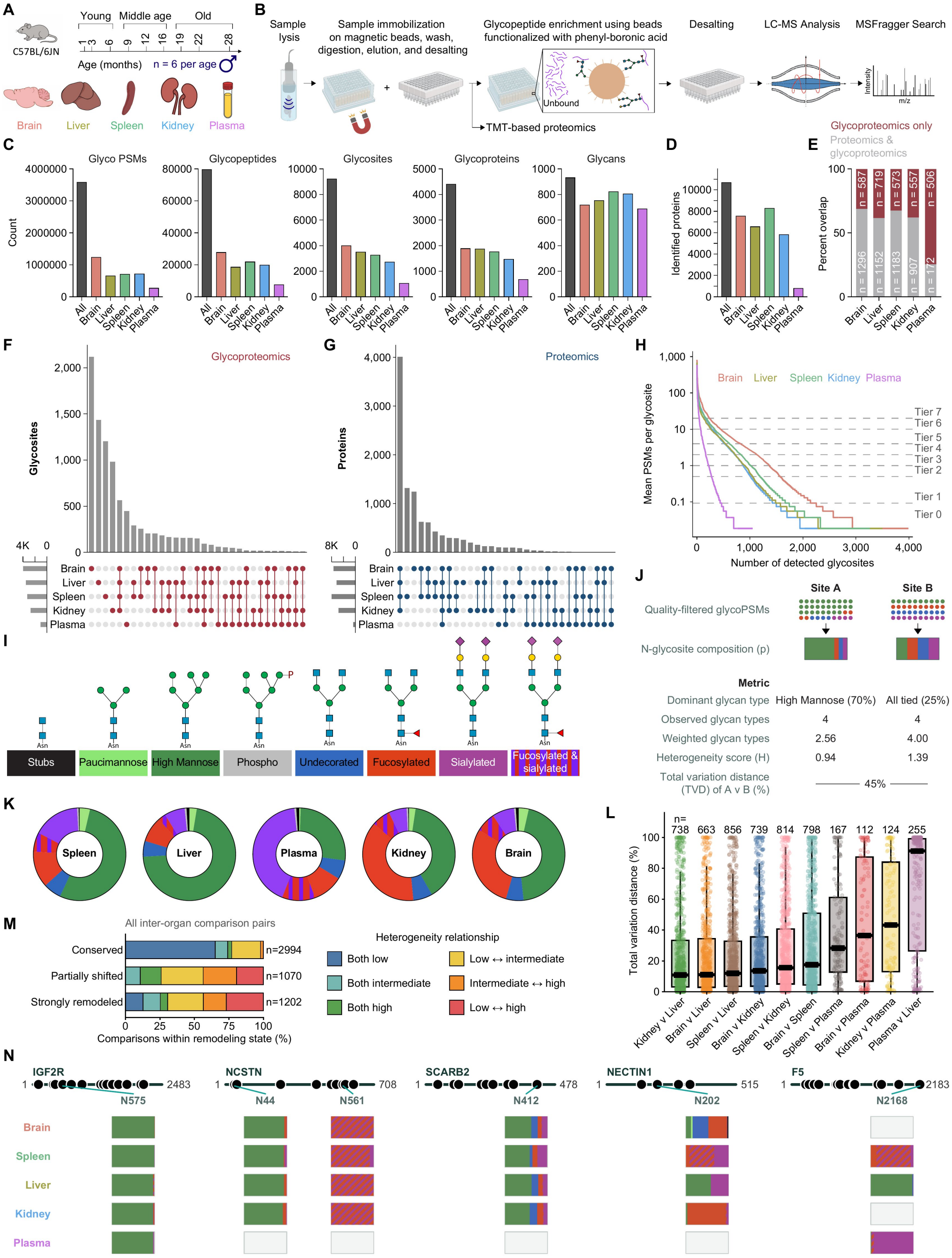
A multi-organ atlas of protein N-glycosylation across the mouse lifespan. **(A)** Cohort overview. Brain, spleen, liver, kidney, and plasma were collected from male C57BL/6JN mice at nine ages from 1 to 28 months (n = 6 per age). **(B)** Experimental workflow. Proteins were immobilized on magnetic beads and digested with trypsin for tandem mass tag (TMT)-based proteomics or enrichment of glycopeptides for glycoproteomics. **(C)** Numbers of quality-filtered glycoPSMs, glycopeptides, glycosites, glycoproteins, and unique N-glycans across each organ in the atlas. **(D)** Quality-filtered proteins identified by proteomics across each organ in the atlas. **(E)** Percent and number of glycoproteins identified only by glycoproteomics or by both proteomics and glycoproteomics across the five atlas organs. **(F and G)** UpSet plots showing detection across organs of glycosites via glycoproteomics **(F)** and proteins via proteomics **(G). (H)** Glycosites ranked by mean glycoPSMs per site in each organ. Dashed lines indicate coverage tiers 0–7, which are used to stratify coverage at each glycosite. **(I)** Composition-based classification of N-glycans into stubs, paucimannose, high mannose, phospho, undecorated, fucosylated, sialylated, and fucosylated & sialylated types. **(J)** Schematic showing how glycan type composition is calculated and the downstream metrics of observed glycan types, heterogeneity score **(H)**, weighted glycan types, and total variation distance (TVD). TVD is half the summed absolute differences between the proportions of each glycan type times 100 to report as a percentage. **(K)** Percent abundance of the eight glycan types in each organ based on pooled glycoPSM counts. **(L)** TVD between organs for glycosites reaching Tier 1+ in both organs. Each point is one shared glycosite; number of comparisons passing this threshold (n) is shown above the distributions. **(M)** Proportions of all 5,266 organ-pair glycosite comparisons classified as conserved (TVD < 20%), partially shifted (20% ≤ TVD < 50%), or strongly remodeled (TVD ≥ 50%), stratified by the relationship between weighted glycan types in the two organs. Low, intermediate, and high heterogeneity denote < 1.5, 1.5 to < 2.5, and ≥ 2.5 weighted glycan types, respectively. **(N)** IGF2R N575 illustrates a conserved site with low heterogeneity dominated by high mannose. NCSTN N44 and N561 illustrate additional conserved sites with low heterogeneity, dominated by high mannose and fucosylated & sialylated glycans, respectively. SCARB2 N412 illustrates a conserved, high-heterogeneity site across the four solid organs. NECTIN1 N202 illustrates strong organ-specific remodeling accompanied by changes in glycan type heterogeneity. F5 N2168 illustrates low-heterogeneity but strongly remodeled compositions dominated by different glycan types across spleen, liver, and plasma. In **(L)**, boxes show medians and interquartile ranges and whiskers extend to 1.5 × the interquartile range. In **(N)**, bars show pooled glycan type proportions and empty boxes indicate that the site was not measured in that organ.

### N-glycan composition and heterogeneity vary across organs and glycosites

We observed a profound heterogeneity of N-glycosylation in our atlas with 931 unique N-glycans which we classified into eight overall glycan types based on their size and monosaccharide composition **(Figures 1C and 1I; Figure S3A)**. The eight glycan types comprise stubs, paucimannose, high mannose, phospho, undecorated, fucosylated, sialylated, and jointly fucosylated & sialylated glycans. Glycan types also differed by subcellular localizations, with high mannose glycans enriched on ER-associated proteins, sialylated glycans on secreted proteins, and paucimannose glycans on lysosomal proteins **(Figure S6C)**. Over 90% of all detected glycoPSMs carried N-glycans between 1–3 kDa, with variable abundance patterns between organs **(Figures S7A–S7E and S7L)**. At the monosaccharide level, Neu5Ac predominated over Neu5Gc in the brain but was nearly absent from plasma, while N2H5 was the most abundant high mannose glycan in the brain and N2H9 predominated in the liver, spleen, and kidney. Meanwhile, in the liver just 4 glycans accounted for over 50% of all detected glycoPSMs while in the kidney it was 10. So despite tremendous available diversity, specific glycans and glycan types frequently dominated in individual tissues.

To compare site-specific variability in glycosylation across organs and ages, we quantified the relative abundance of the eight glycan types at each glycosite and used total variation distance (TVD) to measure differences in composition, with 100% representing no compositional overlap and 0% signifying complete similarity **(Figure 1J; Figure S3C)**. To quantify glycosite heterogeneity, we used the Shannon diversity index to calculate a heterogeneity score (H) that accounts for the number and relative abundance of glycan types, and from H, we calculated the weighted number of glycan types, which reflects how many glycan types meaningfully contribute to a site’s composition **(Figure S3B)**^44^. Glycosites varied widely in their degree of heterogeneity **(Figures S7F–S7H)**. In the brain at CNTNAP1 N861 we observed 83 total glycoPSMs, each instance a single fucosylated glycan (N3H4F1), whereas at the neighboring site N949 on the same protein we observed 130 glycoPSMs and 22 unique glycans across 5 glycan types (3.23 weighted glycan types). More broadly, only approximately 10% of Tier 1+ glycosites within individual organs were represented exclusively by a single exact glycan **(Figure S7J)**. By contrast, after exact glycans were grouped into glycan types, 43–62% of sites were dominated by a single glycan type depending on the organ (≥ 90% abundance), indicating that much exact-glycan microheterogeneity occurs within one predominant glycan type **(Figure S7K)**.

### Organ identity shapes glycosite composition

Global glycan composition differed markedly between organs as previously described, with high mannose glycans predominating in liver, fucosylated glycans enriched in kidney, and sialylated glycans enriched in plasma **(Figure 1K)**^45,46^. Glycosite-level comparisons further characterized these inter-organ differences. Among glycosites reaching Tier 1+ in both organs, pairwise comparisons revealed substantial organ-dependent variation in site-specific composition, with the greatest differences generally occurring between plasma and the solid organs **(Figure 1L)**. Across the 5,266 pairwise inter-organ comparisons, roughly 57% were conserved, 20% were partially shifted, and 23% were strongly remodeled, with variable levels of conserved and strongly remodeled sites between the organ pairs **(Figure 1M; Figure S8A)**. Across organ pairs, conserved sites were frequently low heterogeneity sites dominated by a single shared glycan type in both organs **(Figure 1M; Figures S8B and S8C)**. In most instances these low heterogeneity sites were dominated by high mannose glycans **(Figure 1N; Figure S8D)**, although conservation of more elaborate glycan types across organs was also observed (e.g., fucosylated & sialylated at NCSTN N561). Instances of high heterogeneity sites with conserved composition across multiple organs notably existed but were rare. SCARB2 N412 had high heterogeneity (weighted glycan types > 3 in all solid organs) but strong conservation between all organ pairs (highest TVD was 14.3% for spleen versus kidney). Strongly remodeled sites exhibited a wide range of heterogeneity relationships between the pair. Most common was a scenario where one site was low heterogeneity while the other site was high heterogeneity (e.g., NECTIN1 N202 spleen versus kidney). Remodeled sites where both organ compositions had low heterogeneity but were dominated by a distinct glycan type in each were especially common in the liver-plasma pairs, where in the liver the glycosite was frequently majority high mannose while in the plasma it was sialylated. Together, these findings indicate that glycosite composition reflects an interplay between glycosite identity and organ context: many sites retain a characteristic composition across organs, whereas others undergo substantial organ-specific remodeling.

Glycosite conservation and remodeling were associated with distinct molecular properties **(Figures S9A and S9B)**. High mannose content and ER localization were largely linked with conservation of glycosite composition, whereas sialylation was associated with greater organ-specific remodeling. Curiously, plasma membrane localization for any of the six solid organ comparisons was associated with increased probability of remodeling while it was associated with conservation for the comparisons with plasma. At the level of protein features, disulfide bond content were associated with higher levels of remodeling while being detected across more tissues was associated with greater conservation. Looking at the brain in particular **(Figure S9C)**, the inter-organ comparisons consistently showed that proteins with strongly remodeled glycosites were enriched for involvement in CNS-specific functions (axon, neurogenesis), suggesting that organ-specific glycoforms may serve a functional purpose in cell signaling or development.

### Glycosites on the same protein have similar compositions, particularly when nearby

We next asked whether glycosites on the same protein were more similar than randomly paired glycosites on different proteins. Within each organ, we calculated the total variation distance between every pair of glycosites supported by at least 27 total glycoPSMs (Tier 2+) at both sites, then classified these relationships as being within-protein comparisons or between-protein comparisons. Across all five datasets, the within-protein pairs had more similar compositions than the between-protein pairs, though the magnitude of this difference varied heavily between organs, with the largest differential observed in the plasma and the smallest in the liver **(Figure S10A)**. For each within-protein pair we next calculated the physical distance between each glycosite pair, using solved structures when available and AlphaFold models when not, and retaining distance measurements only from well-resolved paths **(Figure S10)**. Nearby glycosites tended to have more similar glycan compositions in the brain, kidney, and spleen, although this relationship was significant only in the brain and kidney **(Figure S10D and S10E)**. The ECM protein BGN highlighted this relationship between proximity and composition. Detected in the 4 solid tissues, N271 and N312 were consistently of a similar composition, regardless of tissue, even though the composition at these sites could vary between organs **(Figures S10B and S10C)**. Both sites were nearly 100% fucosylated in the brain while in the spleen a heterogeneous mix of high mannose, sialylated, and fucosylated & sialylated glycans was observed at both sites. Thus, glycosites on the same protein tend to share compositional features, particularly when they are close together, while organ identity influences what that shared composition is.

### Aging remodels glycosite composition across organs

To assess age-related changes in glycosite composition, we compared glycosite composition differences between young, middle-aged, and old mice **(Figure 2A; Figures S11A and S11B)**. Aging remodeled glycosite composition in all five organs, although the magnitude was generally smaller than the compositional differences between organs **(Figure 2B)**. In old versus young comparisons, 158 brain, 187 spleen, 90 liver, 126 kidney, and 68 plasma glycosites met the significance threshold **(Figure 2A)**. Age-related remodeling of glycosites appeared strongest early in life – the old versus middle-aged comparison produced the fewest hits and the old versus young comparison produced the most hits in all 5 tissues **(Figure S11C)**. Proteins carrying age-remodeled glycosites were associated with distinct functions across organs: perineuronal net and myelin-related processes in brain, immune and trafficking pathways in spleen, broader cellular trafficking pathways in liver and kidney, and regulatory and homeostatic functions in plasma **(Figure S12A)**. Thus, these associations were largely organ-specific, although endocytic and endolysosomal annotations recurred in spleen, liver, and kidney. The direction of glycan remodeling also varied by organ **(Figure 2C; Figure S11D)**. Among old versus young hits, loss of high mannose as the dominant changing glycan type was more common than gain in all four solid tissues. Brain showed a particularly strong bias toward paucimannose: it was the dominant gain at 26 altered sites but the dominant loss at only 1. An increase in brain paucimannose abundance was also visible at the organ-wide level **(Figure S12B)**. Dominant sialylated gains were more common than losses in spleen and plasma; however, global sialic acid content dropped in the plasma, driven by decreased sialylation on select highly abundant proteins **(Figure S12B)**. Protein abundance changes accompanied some of these shifts. Proteins carrying a remodeling hit showed modestly larger protein absolute abundance changes in the four solid tissues **(Figure S13A)**. Across those tissues, changes in high mannose content were negatively correlated with protein abundance changes **(Figure S14)**. Associations for other glycan types varied by organ: increasing protein abundance was associated with increasing paucimannose content in brain and increasing undecorated glycans in spleen and kidney. Overall, aging drove organ-specific remodeling of glycosite composition, affecting distinct biological pathways and glycan types and, in some cases, accompanying changes in protein abundance.

**Figure 2.**
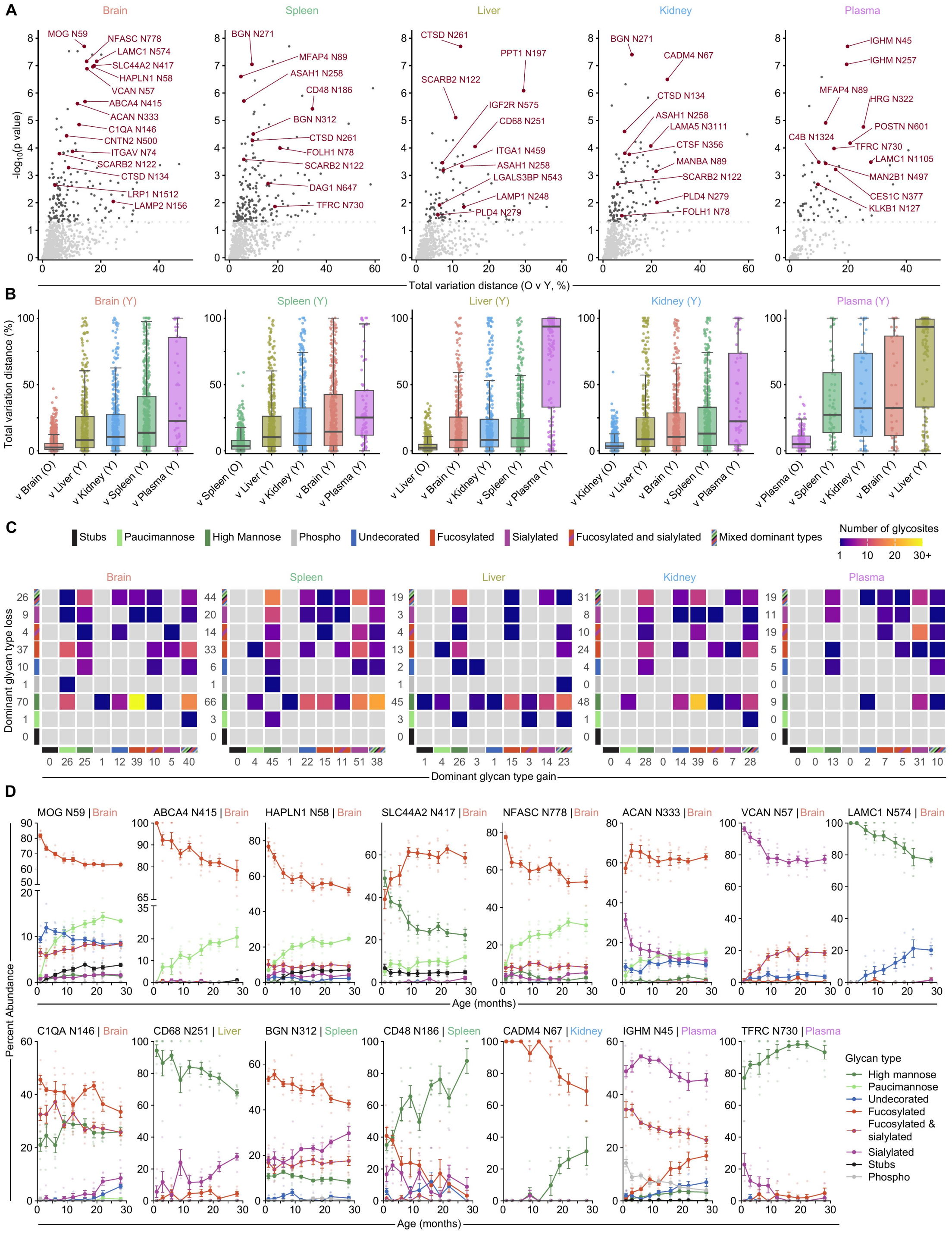
Aging remodels glycosite composition across organs. **(A)** Old versus young (0 v Y) glycosite remodeling in brain, spleen, liver, kidney, and plasma. Each point is a Tier 2+ glycosite detected in at least nine animals with at least 18 total glycoPSMs in each age bin. The x axis shows TVD; the dashed line marks p = 0.05. The comparisons identified 158, 187, 90, 126, and 68 remodeled candidates, respectively. **(B)** TVD distributions comparing old with young animals within the indicated reference organ and comparing that organ with each other organ in young animals. Each point is a shared glycosite. The same inclusion criteria for the O v Y comparison were used for the inter-organ comparisons. **(C)** Numbers of O v Y hits classified by the dominant glycan type lost and gained. Edge values give row and column totals; mixed denotes sites without a single dominant changing glycan type. A glycan type was designated the dominant gain or loss if it accounted for at least 60% of the summed gains or absolute losses, respectively. **(D)** Age trajectories of glycan type percent abundance at representative glycosites. Faint points represent individual animals; symbols and error bars show mean ± SEM. Young **(Y)**, middle-aged (MA), and old **(O)** bins comprise 1–6, 9–16, and 19–28 months, respectively. Boxes in **(B)** show medians and interquartile ranges and whiskers extend to 1.5 × the interquartile range.

Individual glycosites illustrated both these broader patterns and their exceptions **(Figure 2D; Figure S11E)**. Brain THY1 N94 and spleen HSP90B1 N502 provided examples with comparatively stable composition. Brain NFASC N778, MOG N59, ABCA4 N415, and HAPLN1 N58 showed declining fucosylated glycan abundance alongside increasing paucimannose. Other sites followed different trajectories: high mannose declined at liver PPT1 N197, whereas sialylation increased at spleen BGN N312 and plasma HRG N322. Brain PROM1 N291 shifted strikingly from fucosylated toward paucimannose glycans between young and middle-aged animals. Across these diverse trajectories, increased paucimannosylation emerged as a prominent feature of brain aging.

Most proteins with an age-associated glycosite had only one changing site, but a subset showed remodeling at several positions **(Figure S15A)**. In brain, 38 proteins had at least two distinct sites significant in one or more age-bin comparisons; these proteins were enriched for myelin sheath and specialized extracellular matrix annotations **(Figure S15B)**. Four sites on the extracellular matrix protein ACAN gained paucimannose with age, although the glycan type that declined differed among them **(Figures S15C and S15D)**. ACAN N126 lost high mannose, N333 lost sialylated glycans, and N611 and N667 lost fucosylated glycans. NFASC likewise contained sites with different starting compositions and aging trajectories: N778 and N881 showed marked fucosylated-to-paucimannose shifts while N1026 lost high mannose content and gained undecorated and fucosylated glycans **(Figures S15E and S15F)**. Together, these examples show that aging can remodel multiple glycosites on the same protein along distinct compositional trajectories.

Sharing of old versus young hits between organs was limited, with only about 13% of hit glycosites shared across two or more organs **(Figures S13B and S13F)**. Among sites testable in both members of an organ pair, shared hits were more frequent than expected for some pairs, particularly between the spleen, liver, and kidney **(Figures S13C and S13E)**. Proteins carrying hits shared by at least two organs were enriched for lysosomal, endosomal, and extracellular matrix annotations **(Figure S13D)**. Even when a site changed in several organs, the dominant glycan types gained or lost often differed between them **(Figure S13G)**. SCARB2 N122 illustrates both commonality and variation: its high mannose fraction declined with age in all four solid tissues, while the glycan types that increased differed among organs **(Figure S13H)**. Together, these results showed that aging changes glycosite composition, but the affected proteins, direction of remodeling, and trajectories of individual sites depended strongly on organ context.

### The white matter is a hotspot for age-associated paucimannosylation

While many proteins underwent age-related shifts in glycosylation, particularly in the brain, the regional localization of these changes was unclear. To examine spatial differences in N-glycosylation, both within age and across ages, we assembled an orthogonal cohort of 32 total animals aged 3–25 months and performed matrix-assisted laser desorption/ionization mass spectrometry imaging (MALDI-MSI) to visualize the localization of N-glycans in the brain **(Figure 3A; Figure S16)**. Using H&E staining, we annotated 8 distinct brain regions and quantified the abundance of N-glycans at 6 different timepoints, generating a comprehensive spatial atlas of N-glycosylation over time and demonstrating region-specific glycosylation patterns.

**Figure 3.**
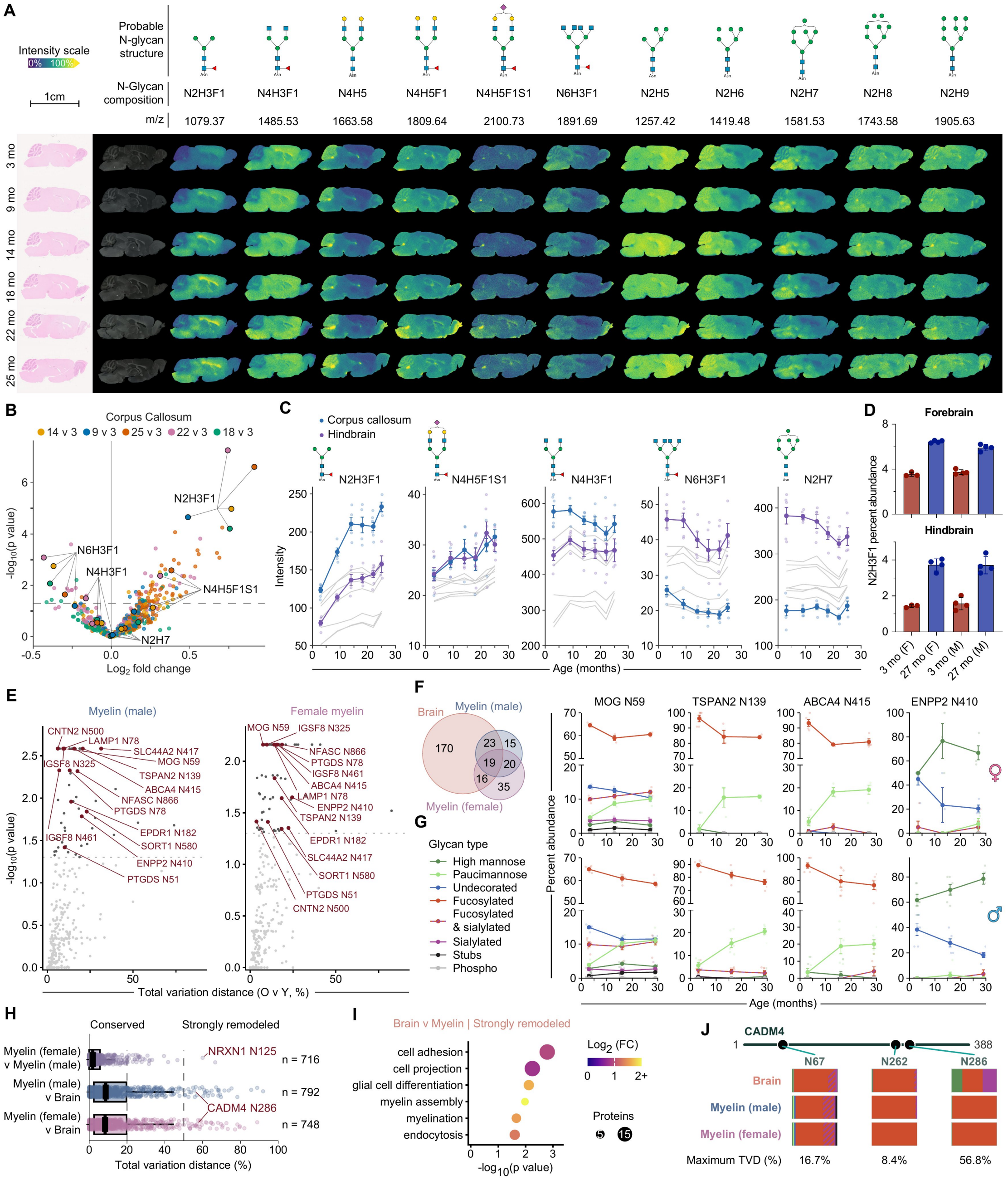
White matter is a hotspot for age-related paucimannosylation. **(A)** MALDI-MSI of N-glycans showing selected N-glycans in sagittal brain sections from mice aged 3-25 months. Rows show representative animals from each age and columns show the indicated compositions, probable structures, m/z values, and regional distribution pattern of the measured species. Images were root mean square (RMS) normalized with hotspot removal performed on the full set of 32 brain images independently for each glycan. **(B)** Corpus callosum abundance changes for each older age versus 3 months. Colors identify the age comparison; the dashed line marks p **=** 0.05 from two-sided Welch t tests on log_2_-transformed intensity. **(C)** Age trajectories of selected glycans in corpus callosum and hindbrain; gray lines show the other annotated brain regions. Points represent individual animals and lines and error bars show mean± SEM. **(D)** Orthogonal glycomics quantification of N2H3F1 in forebrain and hindbrain from 3- and 27-month-old female and male mice (3-month females, n **=** 3; all other groups, n **=** 4). Error bars show mean± SEM. **(E)** Old versus young (0 v Y) plots of glycosite remodeling for male and female myelin. The x axes show TVD, and the dashed lines mark p **=** 0.05. Selected glycosites of interest are annotated. **(F)** Overlap of age-remodeled glycosites in whole brain, male myelin, and female myelin. The set of glycosites includes any site that was a hit in an O v Y, MA v Y, or O v MA comparison. **(G)** Glycan type percent abundance trajectories at selected sites shared between brain and one or both myelin datasets. Upper and lower rows show female and male myelin, respectively; points represent animals and lines and error bars show mean ± SEM. **(H)** Site-level TVD between female and male myelin and between each myelin dataset and whole brain. Dashed lines mark conserved (TVD < 20%), partially shifted (20% ≤ TVD < 50%), or strongly remodeled (TVD ≥ 50%); n values indicate shared sites; boxes show medians and interquartile ranges and whiskers extend to 1.5 × the interquartile range. **(I)** Selected significant GO terms enriched among proteins carrying sites strongly remodeled between brain and both myelin datasets. **(J)** CADM4 site map and glycan type compositions at N67, N262, and N286 in brain and male and female myelin. Values below the bars give the maximum TVD across the three datasets. The MALDI–MSI cohort contained 32 mice across six ages. Male myelin glycoproteomics used 18 mice at 3, 16, and 29 months; female myelin glycoproteomics used 12 mice at 3, 13, and 27 months.

Individual N-glycans exhibited distinct localization patterns. The high mannose glycan N2H5 was enriched in the forebrain, N2H6 and N2H7 were enriched in the hindbrain, and N2H8 and N2H9 were evenly distributed throughout all brain regions **(Figure 3A)**. The undecorated glycan N4H5 was enriched in the cerebellum while the fucosylated form (N4H5F1) was evenly distributed throughout the whole brain. Across all glycans, the paucimannose glycan N2H3F1 exhibited the most dramatic age-related change, particularly in the corpus callosum, where the sialylated N4H5F1S1 glycan also increased with age **(Figures 3B and 3C)**. Concomitantly, larger species with more HexNAc monosaccharides (N4H3F1, N6H3F1) decreased in abundance **(Figure 3C)**. And while abundance of most high mannose glycans was stable across all ages and timepoints, N2H7 levels decreased in the hindbrain but remained stable in the corpus callosum **(Figure 3C)**. To further illustrate the exact structure of N2H3F1 and validate that this increase is observed in males and females, we analyzed the N-glycan composition of the brain using an orthogonal glycomics technique where our samples are directly compared to a library of structurally validated N-glycan standards **(Figure 3D)**. We compared 3- and 27-month-old male and female mice, separating forebrain and hindbrain. Across both regions and sexes we again saw increases of the paucimannose glycan N2H3F1 and validated the proposed structure.

We next performed glycopeptide enrichment on isolated myelin from 12 female mice at 3, 13, and 27 months of age and 18 male mice at 3, 16, and 29 months, and analyzed these data alongside previously reported proteomic measurements from the same cohorts **(Figures S17 and S18)**^47^. Male and female mice showed broadly consistent global patterns of glycosylation **(Figure S18C)**. In both males and females, with age we saw increased paucimannosylation at the global level, a decrease in fucosylated glycans, and at the monosaccharide level a decrease in non-core HexNAc content, consistent with the changes we observed in our MALDI–MSI experiments **(Figures S18B and S18C)**. At the glycosite level, we again calculated significant age-related changes in glycosite composition for both the male and female myelin datasets **(Figure 3E)**. 19 glycosites showed age-associated remodeling in the brain and both myelin datasets **(Figure 3F)**. Among these, ABCA4 N415, CNTN2 N500, LAMP1 N78, and MOG N59 showed remodeling with elevated paucimannose content consistent with the bulk brain data **(Figure 3G; Figure S19A)**. TSPAN2 N139, PTGDS N78, and SORT1 N160 also exhibited shared increases in paucimannose content in both the male and female datasets while the PTGDS N51, STBD1 N135, and NAGA N177 glycosites showed increased paucimannosylation in the brain dataset and one of the myelin datasets **(Figures 3E and 3G; Figure S19A)**. Myelin-specific glycosite remodeling outside of the context of altered paucimannosylation was also observed. ENPP2 N410, IGSF8 N325, and IGSF8 N461 exhibited a loss of undecorated glycans with age and a decrease in non-core HexNAc content **(Figures 3E and 3G; Figures S19A and S19B)**. Overall, four independent datasets consistently identified increased paucimannosylation as a feature of brain aging.

### Myelin- and sex-specific glycoforms

At the individual glycosite level, male and female myelin were highly similar **(Figure 3H)**. Rare sex-associated differences were nevertheless observed. At position N125 on the cell adhesion protein NRXN1, male myelin contained exclusively high mannose glycans, whereas female myelin was predominantly fucosylated **(Figure S19D)**. This difference is particularly intriguing because *NRXN1* expression has been reported to be higher in female oligodendrocytes, while *Nrxn1α* loss produces sex-dependent behavioral phenotypes in mice, including reduced locomotion and affiliative social behavior in females and increased aggression in males^48-50^. As expected, both myelin datasets were more similar to whole brain than to other organs **(Figure S19C)**. Yet this overall similarity concealed sites that consistently distinguished myelin from whole brain: 25 glycosites on 23 proteins were conserved between male and female myelin but strongly remodeled relative to whole brain in both datasets. These proteins were enriched for GO annotations including cell adhesion, glial differentiation, and myelination **(Figure 3I)**. CADM4, a protein involved in cell adhesion, illustrated how this distinct composition depended on the glycosite. N67 and N262 remained relatively similar across datasets, whereas N286 had nearly identical compositions in male and female myelin but differed markedly from whole brain **(Figure 3J)**. Other cell adhesion proteins (NRCAM, TENM4, CNTN2, and CNTNAP1) similarly showed a mixture of stable sites in both myelin and whole brain and myelin-specific glycoforms **(Figures S19E–S19I)**. Together, these findings show that glycosite composition is broadly conserved across whole brain and myelin, while revealing select myelin- and sex-specific glycoforms with potential functional significance.

### HEXB modulates paucimannosylation in oligodendrocyte-lineage cells

The age-related increase in brain and myelin paucimannosylation prompted us to examine proteins that might influence its production. While the exact mechanism of synthesis for some glycans is rigorously worked out in terms of the enzymes needed, the order of action, and the site of this enzymatic activity, this is less clear for paucimannose glycans which are presumed to require, at minimum, the removal of terminal GlcNAc sugars **(Figure 4B)**. HEXA and HEXB, which form β-hexosaminidase complexes, have been proposed as relevant for mammalian paucimannose generation through removal of terminal GlcNAc^51^. HEXA and HEXB abundance increased from young to old in whole brain and in both male and female myelin **(Figure 4A; Figures S20A and S20B)**. Other N-glycan processing enzymes also changed in abundance with age across myelin and the atlas organs. We reasoned that these enzymes might also influence paucimannosylation by changing the availability of glycans that β-hexosaminidase complexes can trim, or by directing those glycans toward other products. We therefore selected eight candidates spanning a range of processing steps (HEXA, HEXB, MGAT1, MGAT2, ST6GAL1, FUT8, MAN2A1, and MAN2A2) to test their effects on global glycan composition and glycosylation at individual sites. We generated stable knockdown and overexpression lines in human oligodendroglioma (HOG) cells, using constitutive dCas9-KRAB expression for knockdown. Viable lines constitutively overexpressing *HEXB, MAN2A 1*, or *FUTB* using the same lentiviral system could not be established. We then collected three replicates of each knockdown or overexpression line, as well as control cells, for input into our established glycoproteomics pipeline.

**Figure 4.**
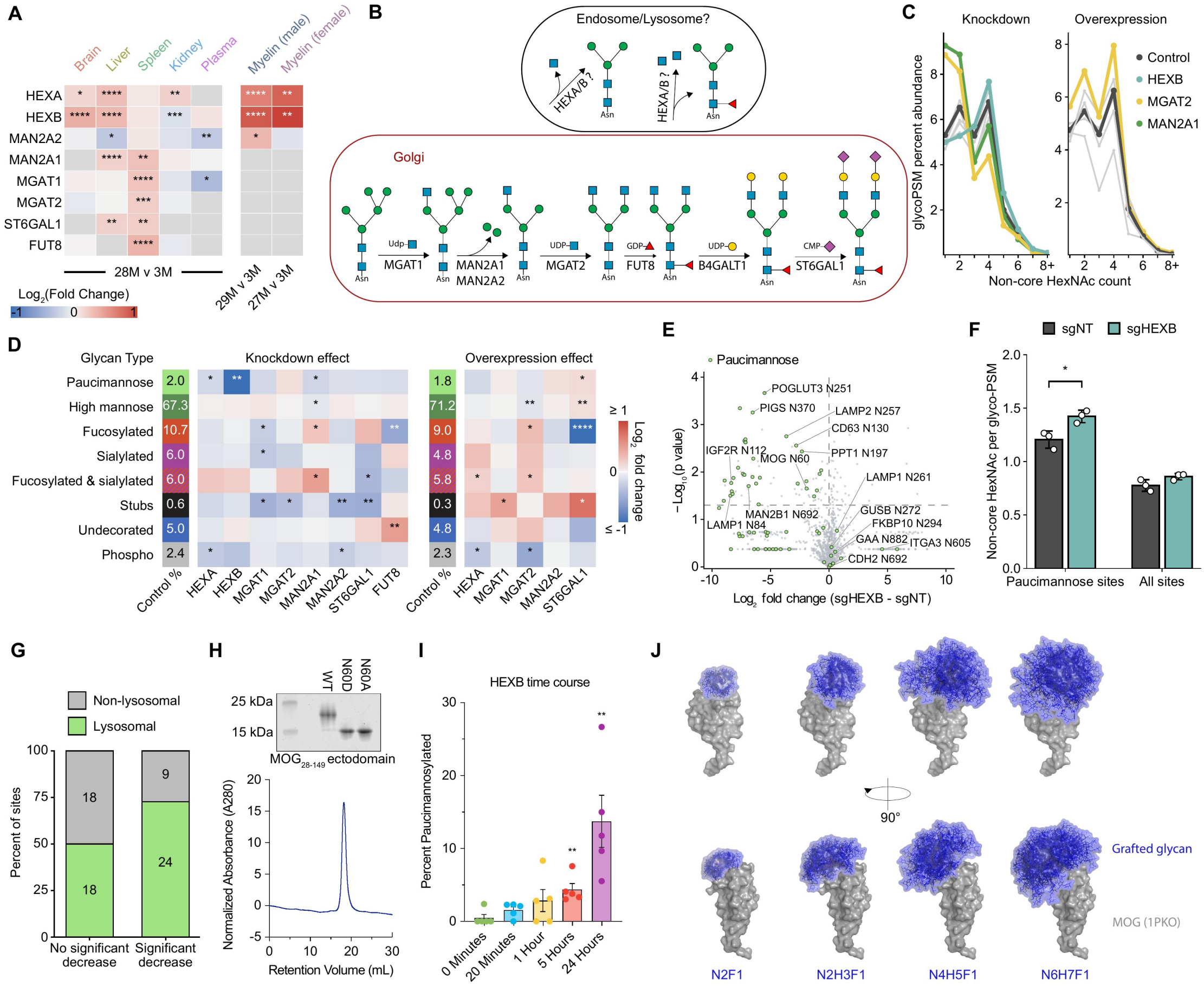
HEXB drives paucimannosylation in oligodendrocyte-lineage cells. **(A)** Age-associated abundance changes in selected N-glycan processing enzymes in atlas organs and male and female myelin. Colors show log_2_ fold change between the indicated oldest group and the corresponding 3-month group; gray denotes an unavailable measurement. Significance symbols denote p values from two-sided Welch t tests comparing the indicated oldest and 3-month groups (*p < 0.05, **p < 0.01, ***p < 0.001, and ****p < 0.0001). **(B)** Schematic of candidate processing steps that generate complex and paucimannose N-glycans. HEXA/HEXB-mediated removal of terminal GlcNAc could occur in an endolysosomal compartment. **(C)** Distribution of glycoPSMs by non-core HexNAc count after knockdown or overexpression of the indicated enzymes in HOG cells. **(D)** Global glycan type composition in control HOG cells and log_2_ fold changes after enzyme knockdown or overexpression. Values in the control columns are the percent abundance values of a given glycan type in control cells; heatmap significance symbols denote p values from two-sided Welch t tests (*p < 0.05, **p < 0.01, ***p < 0.001, and ****p < 0.0001). (E) Site-level fold change in paucimannose percent abundance after HEXB knockdown versus sgNT control. Green points are glycosites carrying paucimannose glycans with selected sites labeled. P values arise from two-sided Welch t tests. **(F)** Mean non-core HexNAc count per glycoPSM across 69 paucimannosylated sites and across all sites in sgNT and sgHEXB cells. Bars and error bars show mean± SD and the asterisk denotes p < 0.05 by a two-sided Welch t test. **(G)** Lysosomal annotation among paucimannosylated sites with (upper left quadrant of panel E) or without a significant decrease in paucimannose abundance after HEXB knockdown. **(H)** SOS-PAGE of purified WT, N60A, and N60D human MOG ectodomains and size-exclusion chromatogram of the purified WT ectodomain. (I) Time-dependent generation of paucimannose glycans at N60 on purified MOG after incubation with recombinant HEXB. Bars show mean± SEM, and points represent biological replicates (n = 5 per time point). **p < 0.01 versus the 0-min condition by a two-sided Mann-Whitney test. **(J)** Structural models of the MOG ectodomain bearing progressively larger N-glycans at N60. Blue conformational ensembles illustrate the increasing surface region sampled by each grafted glycan. The MOG 1PKO crystal structure was used for modeling. HOG cell perturbation analyses in **(C-F)** used three biological replicates per condition.

Perturbing the various N-glycan processing steps produced distinct patterns. *MGA T2* and *MAN2A 1* knockdown shifted glycan abundance toward species containing one or two non-core HexNAcs and away from species containing three or more, whereas *MGA T2* overexpression increased the abundance of both groups **(Figure 4C)**. At the individual glycan level, *MGA T1* knockdown increased Man5 (N2H5) while decreasing its GlcNAc-extended, N3H5 and N3H5F1 products. *FUTB* knockdown reduced fucosylated glycans and increased undecorated glycans and *MGA T2* knockdown depleted branched glycans such as N6H7F1 and N6H7F1S1, both of which increased with *MGAT2* overexpression **(Figure 4D; Figure S21)**. *MAN2A1* knockdown increased fucosylated and fucosylated & sialylated glycans while decreasing high mannose and paucimannose glycans, whereas *MAN2A2* knockdown or overexpression had minimal effects. Thus, perturbing a single processing step can remodel multiple downstream glycan classes, and closely related enzymes do not necessarily have equivalent effects in these cells. The magnitude and direction of these changes also varied among glycosites, emphasizing that global shifts incompletely predict the response at an individual site **(Figure S22)**.

Global paucimannose abundance changed only modestly following most perturbations. *HEXA* and *MAN2A1* knockdown produced a small decrease and *ST6GAL1* overexpression produced a small increase. However, *HEXB* knockdown dramatically reduced paucimannose from approximately 2.0% to 0.46% of glycoPSMs **(Figure 4D)**. This effect extended across individual glycosites: 33 of the 69 paucimannosylated glycosites detected in all sgNT and sgHEXB replicates showed a significant decrease in paucimannose content following *HEXB* knockdown **(Figure 4E)**. The response varied substantially among glycosites **(Figure S22)**. Fucosylated glycans increased at GBA1 N309 while fucosylated & sialylated glycans increased at PLD3 N387. However, as paucimannose declined at GNS N183, GLA N192, and PIGS N370, the proportion of high mannose glycans rose. So despite its function as a hexosaminidase, paucimannose glycans were not exclusively replaced by structures with terminal GlcNAc. Still, non-core HexNAc content increased by 18% at these 69 paucimannosylated sites after *HEXB* knockdown (compared to 11% across all sites) **(Figure 4F)**.

One property influencing how a glycosite responded to *HEXB* knockdown was protein localization **(Figure 4G)**. Paucimannosylated proteins were heavily enriched for lysosomal localization, and of the 33 sites with significant loss of paucimannose, 73% had a lysosomal subcellular annotation while 50% of the 36 sites without a significant loss were annotated as non-lysosomal. Notable exceptions to this pattern were observed. Even sites on the same protein could respond differently. LAMP1 N84 lost paucimannose, whereas LAMP1 N261 retained it. Substantial paucimannose also persisted at GAA N882 despite its lysosomal localization while glycosites on the membrane proteins CDH2 N692 and CDCP1 N310 showed little apparent change and MOG N60 and APMAP N160 lost paucimannose. Together, these data nominated HEXB as a key modulator of paucimannosylation in oligodendrocyte-lineage cells but protein and glycosite-specific factors also influenced paucimannosylation.

To establish that HEXB can directly generate paucimannose, we purified recombinant human MOG ectodomain, confirmed its stability and glycosylation **(Figure 4H)**, and incubated it with recombinant HEXB **(Figure 4I)**. Glycoproteomic analysis demonstrated a time-dependent conversion of glycans at MOG N60 (the human glycosite is shifted by one amino acid) to paucimannose (predominantly N2H3F1), increasing from less than 1% before treatment to approximately 12% after 24 hours **(Figure 4I)**. HEXB alone, outside of a lysosomal context, was therefore sufficient to induce paucimannosylation of the MOG ectodomain.

### An elevated immune-reactive environment with age

Because the MOG ectodomain is relatively compact (∼15 kDa), changes in its N-glycosylation could substantially alter the steric environment of the protein and epitope shielding. Structural modeling predicted that progressively larger glycans at N60 sampled an increasingly broad region of the MOG surface, potentially changing the accessibility of surface epitopes **(Figure 4J)**. This suggested that the functional consequence of MOG glycosylation may depend on the occupancy, size, and composition of the attached glycan. Concurrent with age-related increases in paucimannosylation, we observed increased immunoglobulin (IGKC, IGHG1, IGHG2B, IGHM) detected in the brain and elevated levels of complement proteins (C1q, C4B, CFH) **(Figures S20C and S20D)**. Based on the Neu5Ac content of C1QA, most appeared to be produced by locally resident microglia, but sporadic Neu5Gc-containing PSMs were detected in samples from 16-, 22-, and 28-month-old mice, suggesting that some of the complement entering the brain may also be from an external source or from infiltrating macrophages **(Figure S20E)**. Altered glycosylation therefore has the potential to be another factor modulating the increasingly immune-reactive environment of the aging brain.

### MOG-reactive IgG is detected in individuals without MOGAD

MOG is a known autoantibody target. Clinically, myelin oligodendrocyte glycoprotein antibody-associated disease (MOGAD) is an inflammatory demyelinating disorder of the central nervous system that most commonly presents with optic neuritis, myelitis, or acute disseminated encephalomyelitis^52^. Detection of MOG-reactive IgG is a defining diagnostic feature of the disease. While studies have observed MOG-reactive autoantibodies in control populations, these results are frequently dismissed as ‘false positives’^53,54^. However, there should be a clear distinction between ‘false positive’ as it pertains to a clinical test used to diagnose a full-blown autoimmune disorder and a biochemical assessment of binding. Because MOGAD affects only approximately 1–4 individuals per 100,000, even infrequent MOG reactivity among individuals without a compatible clinical phenotype could substantially outnumber clinically recognized disease^55,56^. We were therefore interested in more thoroughly examining 1) the extent to which autoantibodies targeting MOG could be identified in non-MOGAD patients, 2) whether the antibodies from these different populations exhibited distinct binding properties, 3) the role of the MOG N-glycan in modulating binding, and 4) whether autoantibodies in non-MOGAD patients might contribute to CNS pathology.

Seeking to test a wide range of samples, we obtained plasma from four different sources – Project BIG (n = 40), Stanford Blood Center (n = 15), Stanford Alzheimer’s Disease Research Center (ADRC) (n = 340), and the Stanford Athlete Study (n = 67) **(Figure S23A)**. We grouped these samples into young healthy controls (Young (HC); n = 77), old healthy controls (Old (HC); n = 208), old with a neurological disorder (Old (ND); n = 137), and MOGAD (n = 40). This overall group of 462 plasma samples represented 350 unique patients as 104 patients were sampled multiple times **(Figure S23B)**. Cell-based assays in HEK cells overexpressing MOG are used to test for autoantibody presence, but key aspects of oligodendrocyte identity are not recapitulated in HEK cells (e.g., oligodendrocyte protein expression, membrane content)^57^. We therefore engineered a series of HOG cells to overexpress EGFP-tagged MOG mutants **(Figure 5A; Figure S23F)**. Using flow cytometry, we then screened patient plasma for binding to control cells and MOG-EGFP expressing cells and calculated the IgG Binding Index (IBI), the ratio of antibody signal on MOG-EGFP expressing cells versus control cells. Consistent with the Mayo Clinic’s thresholding, we categorized samples with an IBI > 2.5 as a positive candidate^53^. As expected, MOGAD patients demonstrated high levels of MOG autoantibody positivity, but we also detected autoantibodies in Young (HC), Old (HC), and Old (ND) populations, with a higher proportion in the older populations, though the small cohort size and sex bias of the young samples should be kept in mind **(Figures 5B and 5C; Figures S23C–S23E)**. For 64 Old (HC), 32 Old (ND), and 8 MOGAD patients we also measured binding from a second plasma draw, typically a year apart from the first draw. Reactivity against MOG was remarkably stable between draws across all three groups, with the only major change in signal coming from one MOGAD patient in remission after receiving treatment **(Figure 5D)**. The correlation between IBI in Draw 1 and Draw 2 was exceptionally strong (R^2^ = 0.845) **(Figure 5E)**.

**Figure 5.**
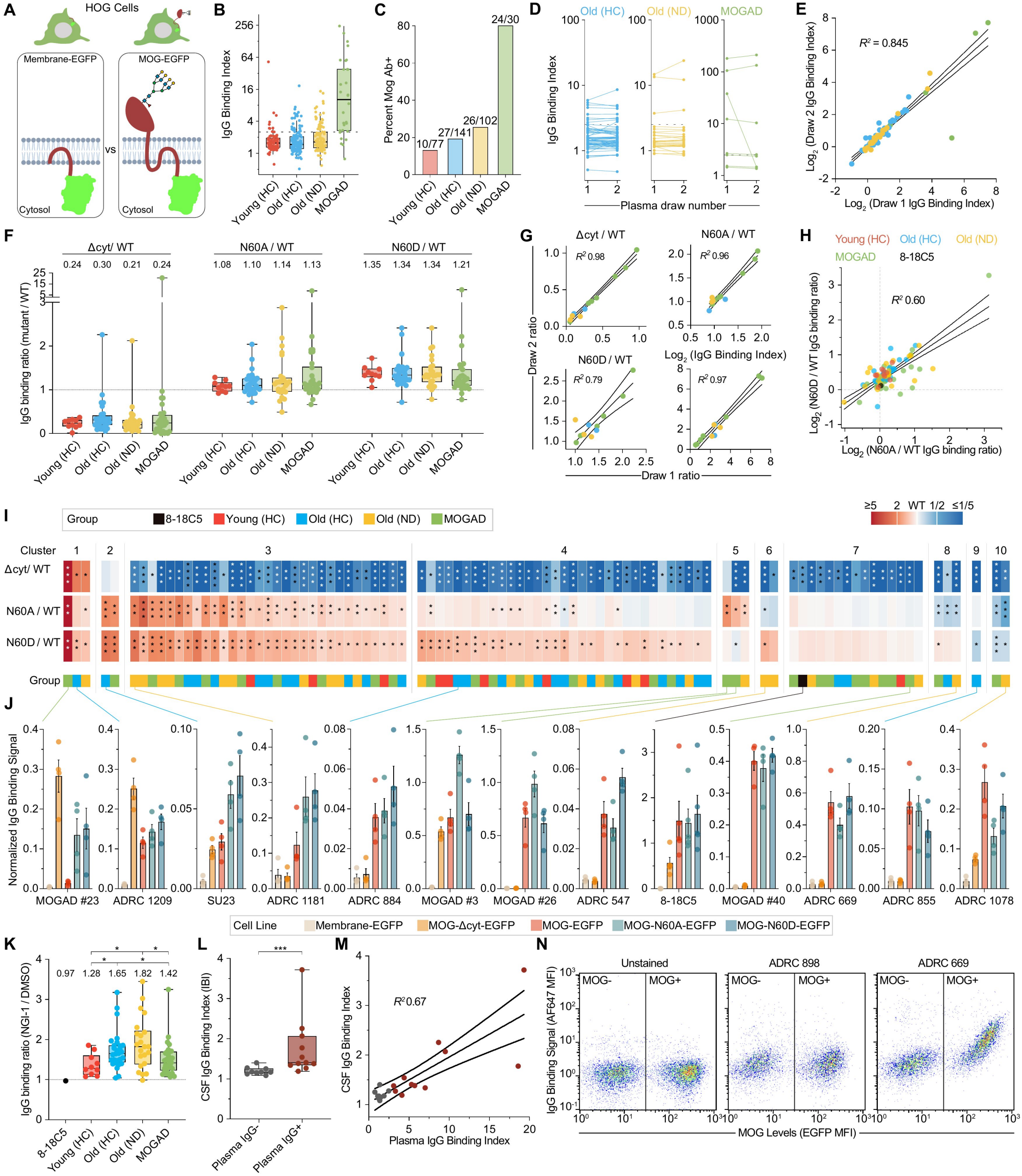
MOG N-glycosylation modulates autoantibody binding across age and disease groups. **(A)** Schematic overview of HOG cells expressing the Membrane-EGFP negative control or MOG-EGFP. **(B)** Box plots of the lgG Binding Index (IBI) for one plasma sample per participant in each group: Young (HC) (n = 77), Old (HC) (n = 141), Old (ND) (n = 102), and MOGAD (n = 30). The dotted line marks the positive candidate threshold of IBI > 2.5. Each value is the mean of two technical replicates. **(C)** Frequency of participants positive for MOG autoantibodies using the IBI > 2.5 threshold. **(D)** Paired IBI values from two plasma draws in Old (HC) (n = 64), Old (ND) (n = 32), and MOGAD (n = 8) participants. (E) Relationship between IBI values from the two plasma draws. Each point represents one participant. The line shows the ordinary least-squares fit with 95% confidence interval. (F) lgG binding ratios for MOG-.Lkyt-EGFP, MOG-N60A-EGFP, and MOG-N60D-EGFP relative to MOG-EGFP in 94 plasma samples: Young (HC) (n = 10), Old (HC) (n = 28), Old (ND) (n = 26), and MOGAD (n = 30). The dotted line marks equal binding to the alternative construct and MOG-EGFP; numbers above the boxes are group medians. **(G)** Agreement between plasma draws for the MOG-.Lkyt-EGFP/MOG-EGFP, MOG-N60A-EGFP/MOG-EGFP, andMOG-N60D-EGFP/MOG-EGFP binding ratios and IBI. For Old (HC) and Old (ND), draw 1 values are means of three biological replicates and draw 2 values are single biological measurements. MOGAD values are means of four biological replicates at each draw. Lines show ordinary least-squares fits with 95% confidence intervals. **(H)** Relationship between MOG-N60A-EGFP/MOG-EGFP and MOG-N60D-EGFP/MOG-EGFP binding ratios. The line shows the ordinary least-squares fit with 95% confidence interval; the 8-18C5 monoclonal antibody control is shown in black. **(I)** Heatmap of mean MOG-Δcyt-EGFP/MOG-EGFP, MOG-N60A-EGFP/MOG-EGFP, and MOG-N60D-EGFP/MOG-EGFP binding ratios for 94 plasma samples (10 Young (HC), 28 Old (HC), 26 Old (ND), and 30 MOGAD) and the 8-18C5 control. Samples were assigned to the 27 possible response patterns using ratios < 0.85 for decreased binding, 0.85–1.15 for unchanged binding, and > 1.15 for increased binding. Ten response patterns were observed. Heatmap color denotes the binding ratio, the lower annotation denotes participant group, and asterisks mark p values from two-sided one-sample t tests of biological-replicate log_2_ ratios against 0. **(J)** Representative samples from the binding response patterns in **(I)**. Bars show mean normalized IgG binding to Membrane-EGFP, MOG-Δcyt-EGFP, MOG-EGFP, MOG-N60A-EGFP, and MOG-N60D-EGFP cells; points show biological replicates and error bars show SEM. **(K)** Effect of N-glycosylation inhibition on MOG IgG binding. Displayed are the IgG binding ratios of the NGI-1 treated MOG-EGFP cells relative to DMSO treatment of MOG-EGFP cells. Binding signal to the correspondingly treated Membrane-EGFP cells is used for background subtraction; the dotted line marks no change and numbers above the boxes are group medians. Brackets denote p < 0.05 by two-sided Mann–Whitney tests. **(L)** CSF IBI in non-MOGAD participants classified as negative (n = 9) or positive (n = 11) for MOG autoantibodies in plasma. Each point is the mean of three biological replicates. **(M)** Relationship between plasma and CSF IBI in the 20 participants from **(L)**. Plasma values are means across 3–4 replicates and CSF values are means of 3 replicates. The line shows the ordinary least-squares fit with 95% confidence interval. **(N)** Representative flow cytometry plots showing CSF IgG binding (AF647 channel) versus MOG-EGFP signal in unstained cells and in plasma-negative (ADRC 898) and plasma-positive (ADRC 669) samples. Vertical gates distinguish EGFP-negative parental HOG cells from MOG-EGFP-positive cells. Boxes show medians and interquartile ranges and whiskers extend to 1.5 × the interquartile range. Where shown, significance symbols denote *p < 0.05, **p < 0.01, ***p < 0.001, and ****p < 0.0001.

### MOG N-glycosylation modulates autoantibody binding

We next sought to determine the distinct binding properties of each patient sample, with a particular interest in how altered glycosylation impacts autoantibody engagement and whether distinct patient populations would show distinct binding properties. Looking at all 4 populations as a whole, deletion of the cytosolic domain and second transmembrane region tended to reduce binding, consistent with the proposed importance of these regions for MOG dimerization^58^ **(Figure 5F)**. Meanwhile, glycosite mutation via either N60A or N60D generally increased binding **(Figure 5F)**. N60D elicited a stronger phenotype, but the effect of both mutations was well correlated, with samples affected by one generally affected by the other (R^2^ = 0.60) **(Figure 5H)**. We also tested blood draws from at least one year apart and saw similarity in binding properties as well – when Draw 1 IgG bound better to N60D, Draw 2 IgG exhibited the same behavior **(Figure 5G)**. When Draw 1 IgG bound worse after cytoplasmic domain deletion, so did Draw 2 IgG, suggesting that not just presence of any antibody but presence of antibodies with the same binding properties was stable.

To further delineate the binding properties of each patient sample, we clustered them based on their response to each mutant, generating 10 clusters of binding patterns **(Figures 5I and 5J)**. The most abundant clusters (clusters 3 and 4) exhibited increased binding to one or both glycosite mutants and decreased binding to MOG-Δcyt-EGFP. Samples that bound worse to all mutants (cluster 10) were rare. Another rare but previously unobserved phenotype was increased binding to MOG-Δcyt-EGFP, which also coincided with increased binding to the N60A and N60D mutants in three samples. Notably, clusters exhibited a diverse spread of patient samples within them – no one cluster was dominated by a single patient group, suggesting that there are no specific pathogenic and non-pathogenic epitopes on MOG.

To further demonstrate the importance of the MOG N-glycan (and indeed other N-glycans) on the cell surface, we treated cells with the oligosaccharyltransferase inhibitor NGI-1 and repeated the cell-based assays **(Figure 5K)**. We found that after NGI-1 treatment, MOG autoantibody binding increased in all populations, though Old (HC) and Old (ND) samples showed the strongest effect. These data demonstrate that both the glycan on MOG and the broader collection of cell surface glycans can modulate autoantibody binding.

Finally, to examine the potential clinical impact of these autoantibodies, we tested CSF from 20 non-MOGAD patients – 9 that were negative for MOG-reactive IgG in the plasma and 11 that had tested positive. The patients that were positive for MOG-reactive IgG in the plasma showed a significant increase in the presence of MOG-reactive antibodies in the CSF, and the degree of this positivity correlated with plasma IBI, with one patient (ADRC 669) demonstrating particularly strong evidence for MOG-reactive IgG circulating in the CSF **(Figures 5L–5N)**. These data raise the possibility that with age low levels of autoantibodies against proteins like MOG (or the plasma B cells that produce them) can leak into the brain, not at a level sufficient to cause a full-blown autoimmune disorder, but perhaps at sufficient amounts to contribute to local damage. Decreased glycan length (and reduced epitope shielding) may then further exacerbate the situation.

### MOG_35–55_-induced EAE recapitulates a subset of age-associated glycosite changes

To test the *in vivo* effects of MOG-directed autoimmune neuroinflammation and demyelination, we induced EAE in mice using the MOG3_5 – 55_ peptide **(Figure 6A)** and collected the spinal cords for glycoproteomic and proteomic assessment in our established workflows **(Figures S24 and S25)**. We observed clear changes in animal locomotion and canonical signatures of EAE (elevated TF, VCAM1, VIM, C1QA, C48) at the protein level **(Figures 68 and 6C)**. 97 glycosites showed differential glycosylation after EAE induction **(Figure 6D)**. Of those, 23 glycosites on 21 unique proteins also showed differential glycosylation in the aging brain **(Figure 6E)**. This group was enriched for lysosomal and immune response-related proteins, beyond the already existing enrichment for these pathways in the full set of proteins bearing the 97 altered glycosites **(Figure 6G)**. Comparing the degree of change at these differentially glycosylated sites, EAE induced larger changes both across all differentially glycosylated sites and among the 23 shared sites **(Figure 6F)**. At these 23 shared hit sites the compositional changes were also largely conserved **(Figure 6H)**, though the magnitude of change was frequently more pronounced in EAE versus aging **(Figure 61)**. Of particular note was C1QA. At the latest timepoints we observed increased sialylated glycans and undecorated glycans while fucosylated and fucosylated & sialylated glycans showed a notable decrease. These same changes were observed after EAE induction, with even stronger effect sizes and a clearer shift towards Neu5Gc-containing glycans **(Figures S25G-S25H)**, which could be explained by the large number of infiltrating macrophages after EAE^59^. ITGAV N74, LRP1 N1512, LAMP2 N156, SCARB2 N122, and CTSD N134, among other sites, further demonstrated the general pattern of similar directional change as aging but at a larger magnitude. Glycosylation and immune responses may therefore form a bidirectional relationship: glycan composition can shape antigen recognition, while the resulting immune response can in turn initiate programs that remodel glycosylation. These data further reveal shared signatures of immune-mediated damage in EAE and aging and raise the possibility that overlapping mechanisms contribute to both.

**Figure 6.**
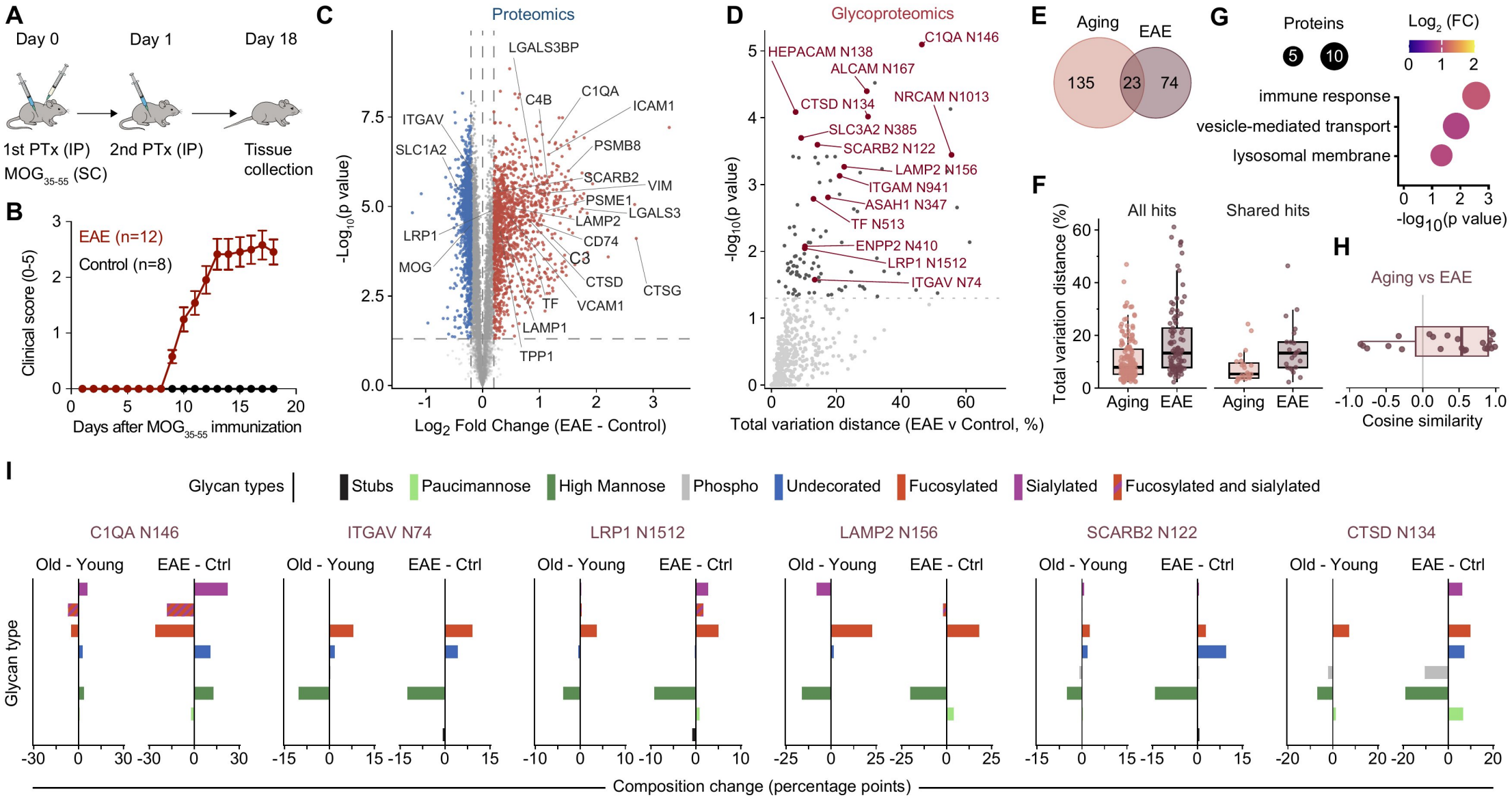
EAE recapitulates a subset of age-associated glycosite changes. **(A)** EAE induction and collection schedule. Mice received MOG35-55 and the first pertussis toxin injection on day O and a second pertussis toxin injection on day 1; tissue was collected on day 18. **(B)** Clinical scores after MOG35-55 immunization in EAE mice (n = 12) versus control mice (n = 8). Values are mean± SEM. **(C)** Spinal cord proteomic changes in EAE mice versus control mice. Selected EAE-associated proteins are labeled. Significance testing uses a two-sided Welch t test. **(D)** Spinal cord glycosite composition changes in EAE mice versus control mice. The x axis shows TVD; the dashed line marks p = 0.05 and selected sites are labeled. **(E)** Overlap of old versus young (0 v Y) brain hits and EAE versus control spinal cord hits. Twenty-three glycosites on 21 proteins were shared. **(F)** TVD distributions for all hits and the 23 shared hits in aging and EAE. Points represent glycosites and boxes show medians and interquartile ranges and whiskers extend to 1.5 x the interquartile range. **(G)** GO enrichment among proteins carrying the 23 shared glycosite hits, using proteins carrying EAE-versus-control glycosite hits as the background. **(H)** Cosine similarity between the signed glycan type composition changes in aging and EAE at the shared sites. Positive values indicate concordant remodeling. (I) Glycan type composition changes at representative shared sites in old versus young brain and EAE versus control spinal cord. Bars show changes in each glycan type, expressed in percentage points.

## Discussion

New technologies and methodologies are allowing ever finer investigation into how tissues, cells, organelles, and biomolecules change with age. In this context, we provide a new framework for understanding an understudied level of complexity -N-glycosylation - between organs, with age, and during neuroinflammation and demyelination. In these areas we showcase clear differences in N-glycosylation as well as persistent patterns of stability. At the glycosite-level, an old brain is more like a young brain than an old liver. The N-glycan composition at detected glycosites is largely stable with age, but there are notable deviations from this pattern. Further, there are unique organ signatures based on the global monosaccharide content (elevated Neu5Ac/Neu5Gc in the mouse brain) and glycan type content (high levels of fucosylated type glycans in the kidney) as well as distinct glycosite-specific patterns (unique NECTIN1 N202 glycosylation patterns in each organ). It is unlikely that this complexity exists for no reason other than decoration. N-glycans clearly perform functions beyond assisting in protein folding, and indeed for a subset of proteins their function appears to be something entirely unrelated to protein folding^60,61^. The sole MOG glycosite can be mutated with no impact on protein stability or expression levels and is therefore dispensable for proper folding, yet this site has high glycan occupancy and is conserved across species from humans to quokkas^62,63^. As we demonstrate, it can function as an epitope shield / modulator of autoantibody binding, but it is likely to serve other yet unknown functions.

Investigating the functional significance of a distinct N-glycan at a specific glycosite is a uniquely challenging problem. Cellular N-glycan processing enzyme levels can be modulated (as we show here) but the effects are not restricted to a single site. Alternatively, individual N-glycosites can be mutated (as we do at MOG N60) to demonstrate functional importance of the site, but the impact of specific glycoforms is then impossible to probe. Installing a specific N-glycan at a specific N-glycosite is therefore a key goal of glycobiology. To an extent this is achievable in an *in vitro* context with purified / soluble proteins, where sequential enzymatic treatment and lectin-based purifications can enrich for specific glycoforms^64^. But such an approach is laborious and limited to the study of a select subset of proteins. Production of a recombinant ectodomain with a selected N-glycan installed, followed by its covalent attachment to a membrane-embedded construct retaining the native transmembrane and cytoplasmic regions, could in principle be achieved using sortase-mediated ligation or split-intein-mediated protein trans-splicing^65-68^. Such semisynthetic reconstruction would be technically demanding and less readily generalizable to multipass membrane proteins, for which topology, folding, and assembly must also be preserved. Alternatively, chemoenzymatic approaches to remove and reinstall specific glycoforms on the cell surface are extremely appealing and have been pursued, but additional protein targeting is required to make this site-specific^69,70^. Improving these technologies is therefore an exciting next step for elucidating the functional significance of specific glycans at specific sites, and our work has nominated dozens of candidates across tissues.

Many of the age-related glycosite changes merit further study. In peripheral tissues, these included increased sialylation of CD68 N251 in the liver, BGN N312 in the spleen, and HRG N322 in plasma. CD68 glycosylation is also altered by inflammatory stimulation and phagocytosis^71^. Its age-associated shift toward sialylated glycans may therefore reflect broader changes in macrophage biology or could itself be contributing to a more pro-inflammatory state via altered glycan-mediated protection from the lysosomal environment, altered trafficking to the cell surface, or engagement with other proteins on the cell surface. Sialylation of BGN could similarly influence its interactions with innate immune receptors or its organization of the extracellular matrix, whereas sialylation of HRG could modify its numerous interactions with coagulation, fibrinolytic, complement, and immune proteins^72-74^. Other sites, including CD48 N186 in the spleen and TFRC N730 in the plasma, showed increased high mannose glycans and reduced glycan heterogeneity. For CD48, such changes could affect communication / signaling between immune cells^75^. In the case of shed TFRC ectodomain, these detected changes in glycosylation could be used alongside other released lysosomal proteins (MAN2B1 N497), secreted ECM proteins (POSTN N601, LAMC1 N1105), and circulating factors (CP N138, C4B N1324) to assess biological age on a novel axis, although conservation in humans would need to be confirmed^76^. Multiple IGHM glycosites also changed across plasma and peripheral tissues. IgM normally exhibits a strongly site-dependent glycosylation profile, and its C-terminal N441 site is typically high mannose and contributes to J-chain incorporation and polymer assembly^77,78^. Age-dependent loss of high mannose glycans and increased heterogeneity at these sites could therefore affect IgM structure or effector activity. At other IGHM sites, sialylated glycans decreased with age. This parallels a broader age-associated decline in immunoglobulin sialylation and could alter IgM persistence or inflammatory functions and interactions with complement proteins^79,80^.

In the brain, increased paucimannosylation extended beyond myelin proteins to several physically connected components of perineuronal nets, including multiple sites on ACAN as well as HAPLN1 N58 and TNR N470. Aging alters hyaluronan integrity and the solubility of ACAN and HAPLN1^81,82^. Experimental studies have also shown that glycosylation can influence ACAN epitope accessibility and hyaluronan binding and that distinct HAPLN1 glycoforms differ in their ability to stabilize hyaluronan-proteoglycan complexes^83-85^. Increased paucimannosylation could therefore contribute directly to altered matrix interactions, protease susceptibility, or lectin recognition. Or it could serve as a molecular record of enhanced ECM turnover and endolysosomal processing.

PROM1/CD133 provides a separate but intriguing candidate with an age-related increase of paucimannosylation. PROM1 is associated with neural progenitor cells (NPCs), and paucimannose glycans have independently been implicated in regulating early postnatal NPC proliferation^86,87^. Aging was associated with increased N2H3F1 and decreased N4H3F1 at PROM1 N291 and N332, consistent with removal of two terminal GlcNAc residues by HEXB. This is notable because PROM1 glycosylation can influence protein fate. Sialylation protects PROM1 from lysosome-dependent degradation, while complex N-glycosylation affects its endosomal trafficking and stability in glioma stem cells^88,89^. The age-associated shift toward paucimannose could therefore alter PROM1 stability, cell-surface residence, or function. Alternatively, it may be a consequence of increased endolysosomal trafficking as PROM1-expressing cells change state.

Collectively, these findings raise the possibility that site-specific paucimannosylation is both a marker and a mediator of age-related changes. While our data and the work of others clearly support a role for HEXB in paucimannosylation, it is unclear where inside (or outside) a cell this enzymatic activity takes place and when in a protein’s lifetime this occurs^51,90^. The high incidence of paucimannosylation on lysosomal proteins is consistent with the localization of HEXB, but this does not explain how membrane proteins and ECM proteins become paucimannosylated. Non-lysosomal proteins such as MOG could conceivably become paucimannosylated (1) during ER/Golgi synthesis, (2) after endolysosomal trafficking, (3) during recycling from the plasma membrane, or (4) through extracellular HEXB. Tied to the question of where a protein becomes paucimannosylated is the issue of when it becomes paucimannosylated. N-glycans are installed in the ER and trimmed in the Golgi, but the localization and mechanism of HEXB activity need not require generation right after synthesis. Post-synthesis processing via extracellular HEXB is therefore an intriguing possible mechanism, as HEXB was recently shown to be secreted in the brain and this could explain the increased incidence on membrane proteins (MOG, ABCA4, TSPAN2) as well as ECM proteins (HAPLN1, ACAN, TNR)^91^. Indeed, our *in vitro* experiments showed that HEXB could actively induce paucimannosylation of MOG at a neutral pH outside of a lysosomal context. Still, the lysosome may remain the principal site of glycan trimming, with paucimannosylated proteins subsequently reaching the cell surface or extracellular space through endolysosomal trafficking or lysosomal exocytosis.

Autoantibodies are detected in individuals without a clinically recognized autoimmune disease and become increasingly prevalent with age, demonstrating that seropositivity and overt autoimmune disease are not equivalent^30,92^. This does not necessarily mean that autoantibodies in these individuals are innocuous. Some may contribute to subtle symptoms or cumulative tissue dysfunction without producing a discrete autoimmune disorder, as is the case with persistent circulation of anti-NMDAR1 autoantibodies in mice^93^. Consistent with this continuum, patients with CNS autoimmunity can remain seropositive during prolonged periods of remission, and antibody titer is an incomplete predictor of disease activity^94^. Aging may alter several of the conditions that govern whether autoreactivity remains clinically silent or becomes damaging, yet the identities and consequences of brain-reactive autoantibodies during normal aging remain poorly understood. High-throughput peptide microarray and peptide phage display technologies are exciting technologies for this space, but the antigen-autoantibody interaction cannot always be recapitulated in such systems^95,96^. Target identity is critical for choosing the right assay for autoantibody detection. Peptide display is a poor strategy for detecting MOG autoantibodies – only with the advent of cell-based assays displaying full length MOG was reliable autoantibody testing possible^57^. Dependence of autoantibody recognition on post-translational modifications, such as N-glycan-dependent recognition of CNTN1, and on quaternary epitopes spanning distinct protein subunits, such as the GABA_A_ receptor α1–γ2 interface, further complicates autoantibody screening^60,97^.

The identity of the target further helps define what kind of damage is possible, but does not by itself determine whether damage will occur. AQP4-IgG, for example, has clear pathogenic potential, but its effects still depend on tissue access and downstream effector mechanisms^98,99^. Autoantibody pathogenicity therefore emerges from the interaction between the antibody, its antigen, and the surrounding tissue. Antibody abundance, affinity, class, and glycosylation shape its binding and effector potential. Tissue barriers govern whether it reaches its target. Antigen abundance, organization, and glycosylation influence whether productive binding occurs. And the local availability of complement and effector cells determines the resulting damage. Among these variables, glycosylation of the autoantibody has received considerable attention^100,101^. Our findings join a small number of studies demonstrating that target-protein glycosylation can also shape the autoimmune response by modulating antibody binding^60,62,102^. This principle extends beyond autoantibody recognition, as site-specific glycans can promote, obstruct, or tune interactions with receptors, ligands, lectins, and other binding partners. Together, our analyses establish N-glycan composition as a biologically meaningful layer of tissue state and provide a foundation for determining how specific glycans shape diverse molecular interactions in aging and disease.

### Limitations of the study

Glycopeptide sampling depth varied across glycosites, and many sites lacked sufficient coverage to rigorously test age-related changes. The remodeling we described therefore likely represents only a subset of the changes that occur with aging. Our measurements were also generated largely from bulk tissues, where cell-type-specific changes may be diluted or obscured. Future studies combining improved glycoproteomic sampling depth with cell type resolved profiling will be needed to detail these additional age-associated changes. Finally, our enrichment strategy measures glycan composition among the glycopeptides detected at a site but does not determine the frequency with which that glycosite is actually glycosylated since any unmodified peptides are lost during enrichment. Defining how occupancy varies across organs, cell types, and age therefore remains a largely unexplored and potentially important dimension of aging biology.

## Materials and Methods

### Animal husbandry and organ collection

6 total animal cohorts were utilized. The primary multi-organ aging cohort (cohort 1) comprised 54 male C57BL/6JN mice collected at 1, 3, 6, 9, 12, 16, 19, 22, and 28 months of age (n = 6 per age), from which brain, spleen, liver, kidney, and plasma were obtained. A separate MALDI–MSI cohort (cohort 2) comprised 32 male C57BL/6JN mice aged 3, 9, 14, 18, 22, and 25 months (n = 6, 5, 5, 5, 7, and 4, respectively). An orthogonal brain glycomics cohort (cohort 3) included male and female C57BL/6JN mice at 3 and 27 months of age (n = 3 for 3-month-old female animals and n = 4 for all others; n = 15 total), with forebrain and hindbrain analyzed separately. Myelin studies used two additional C57BL/6JN cohorts: 18 males aged 3, 16, and 29 months with 6 animals per age (cohort 4) and 12 females aged 3–4, 13, and 27 months (n = 5, 3, and 4, respectively) (cohort 5). The 3–4-month-old animals were comprised of three 3-month-old and two 4-month-old mice and are analyzed together and referred to as the 3-month (‘Young’) group for simplicity. For the myelin cohorts all takedown protocols and proteomic analysis of these cohorts were previously published^47^. Finally, the experimental autoimmune encephalomyelitis (EAE) experiment used female C57BL/6J mice, including 12 EAE and 8 control animals (cohort 6); mice were 11–13 weeks old at EAE induction, and spinal cords were collected 18 days later.

C57BL/6JN mice were shipped from the National Institute on Aging colony at Charles River or bred in house. C57BL/6J mice were purchased from The Jackson Laboratory (stock no. 000664). All mice were housed in cages of up to 5 mice at the Stanford ChEM-H animal facility (cohorts 1–5) or the Stanford School of Medicine animal facility (cohort 6) under specific pathogen-free (SPF) conditions on a 12 h light/dark cycle with ad libitum access to food and water. Animals from cohorts 1–5 were acclimatized to the facility for at least one month before takedown and cohort 6 animals were acclimatized to the facility for at least 7 days before immunization.

For cohorts 1–3, mice were anesthetized with isoflurane and transcardially perfused with at least 20 mL of 1× PBS. If plasma was to be collected, blood was extracted before perfusion, using a syringe pre-loaded with approximately 50 µL 0.5 M EDTA. The blood samples were then spun at 1,500 × g for 15 min at 4°C to pellet red blood cells and the supernatant (plasma) was removed to a fresh tube. After perfusion, organs were immediately extracted and snap-frozen in tubes on dry ice (cohort 1) or fixed in 10% neutral buffered formalin (cohorts 2 and 3). For the fixed samples, 24 h later samples were transferred into fresh tubes containing 70% ethanol before being processed into formalin-fixed paraffin-embedded (FFPE) tissue blocks. For cohort 6, animals were anesthetized as above with the additional step of an injection of Avertin (2,2,2-tribromoethanol, 200 mg/kg) before perfusion. Immediately after perfusion, the spinal column was dissected free from surrounding skin, musculature and viscera. The caudal vertebrae and pelvic bone were trimmed away to expose the distal end of the spinal canal. The intact spinal cord was then isolated by hydraulic extrusion. A trimmed 1,000 µL pipette tip mounted on a 5 mL syringe pre-loaded with sterile 1× PBS was fitted into the caudal opening of the spinal canal, and sharp pulses of PBS were applied to extrude the cord through the rostral end into a Petri dish. Isolated spinal cords were briefly washed in a second Petri dish containing PBS, then snap-frozen on dry ice and stored at −80 °C until further processing. All animal care and procedures complied with the Animal Welfare Act and were in accordance with institutional guidelines and approved by the institutional administrative panel of laboratory animal care at Stanford University.

### EAE induction and scoring

EAE was induced in 12 female C57BL/6J mice using the MOG_35–55_/CFA Emulsion PTX kit (EK-2110, Hooke Laboratories) according to the manufacturer’s instructions. On day 0, mice received the MOG_35–55_/CFA emulsion at two subcutaneous sites (100 µl per site; 200 µl total). Two hours later, each animal received an intraperitoneal injection of 110 ng pertussis toxin (PTX) in 100 µl cold PBS, diluted immediately before use from a 200 ng µl^-1^ stock; a second PTX injection was repeated 24 h after the first injection^103^. Eight age-matched female C57BL/6J mice served as controls. All injections were performed under isoflurane anesthesia.

Mice were examined daily post-immunization and assigned a clinical score on the standard 0–5 EAE scale (in 0.5 increments) as described by Hooke Laboratories cat. no. EK-2110: 0, no clinical signs; 0.5, tip of tail limp; 1.0, complete tail limpness; 1.5, limp tail with hind-leg inhibition; 2.0, limp tail with hind-leg weakness (wobbly gait); 2.5, partial hind-limb paralysis (dragging of hind legs); 3.0, complete bilateral hind-limb paralysis; 3.5, complete hind-limb paralysis with inability to right; 4.0, complete hind-limb and partial fore-limb paralysis; 4.5, moribund; and 5.0, death or euthanasia due to paralysis (a score of 5.0 was recorded for that animal thereafter)^104^. Supplemental gel food was provided to all cages during the early phase of disease and thereafter to cages housing animals with a clinical score ≥ 3.0. Animals with a clinical score ≥ 3.0 received additional supportive care in the form of subcutaneous saline injections. Mice reaching a score of 4.0 received subcutaneous fluids and softened food on the cage floor, and animals meeting institutional criteria were euthanized. Tissues were collected on day 18 as detailed above.

### Human participants and biospecimens

Plasma samples were obtained through Project BIG, the Stanford Blood Center, the Stanford University Alzheimer’s Disease Research Center (ADRC), and the Stanford Athlete Study. The dataset contained 462 plasma samples from 350 unique participants. For the primary MOG autoantibody screen, one representative plasma sample was displayed for each participant, yielding 77 Young (HC) samples, 141 Old (HC) samples, 102 Old (ND) samples, and 30 MOGAD samples. The Old (ND) group included participants with Parkinson’s disease, Alzheimer’s disease, mild cognitive impairment, or Lewy body disease. When more than one plasma sample was available, the earliest dated blood draw was selected to represent that patient in the initial screen. Cerebrospinal fluid (CSF) was analyzed from 20 participants without MOG antibody-associated disease, including 9 whose plasma screen was negative and 11 whose plasma screen was positive for antibodies binding to MOG.

### Plasma collection and processing

The collection of plasma was approved by the Institutional Review Board of Stanford University and written consent was obtained from all subjects. Blood collection and processing were done according to a rigorous standardized protocol to minimize variation associated with blood draw and processing. For ADRC samples, about 10 mL whole blood was collected in a vacutainer EDTA tube (BD Vacutainer EDTA tube) and spun at 3,000 rpm for 10 min to separate out plasma, leaving 1 cm of plasma above the buffy coat and taking care not to disturb the buffy coat to avoid cell contamination. Plasma processing takes about an hour from the time of the blood draw to the time of freezing and storage. All healthy control participants were deemed cognitively unimpaired during a clinical consensus conference that included neurologists and neuropsychologists. Cognitively impaired subjects underwent Clinical Dementia Rating and standardized neurological and neuropsychological assessments to determine cognitive and diagnostic status, including procedures of the National Alzheimer’s Coordinating Center (https://naccdata.org/). All impaired participants included in this study were deemed cognitively impaired during a clinical consensus conference that included neurologists and neuropsychologists. Project BIG (projectbig.com) contributed 40 plasma samples from 30 participants with MOGAD. Participants meeting the diagnostic criteria for MOGAD were enrolled in the study inclusive of demographic and clinical information^52^. The plasma aliquots used in this study were generated from blood collected using either EDTA or heparin as the anticoagulant; 4 samples were EDTA plasma, and 36 were heparin plasma. When multiple samples were available from the same participant, the earliest collection was used in the MOG autoantibody binding analyses, resulting in 4 EDTA and 26 heparin plasma samples. Fifteen citrate-anticoagulated plasma samples from young and older healthy donors had been acquired directly from the Stanford Blood Center. The Stanford Athlete Study contributed 67 plasma samples collected from collegiate football and volleyball athletes.

### Cerebrospinal fluid (CSF) collection and processing

Cerebrospinal fluid (CSF) was collected by lumbar puncture (LP) performed by trained clinical personnel according to standardized study procedures. Following collection, CSF samples were transferred into sterile, labeled collection tubes and transported to the laboratory for processing. Samples were centrifuged at 1,500 rpm to remove cellular components and debris. The resulting supernatant was aliquoted into labeled polypropylene tubes and stored at −80°C until further analysis. Sample collection, processing, storage, and handling were performed using standardized procedures to minimize preanalytical variability.

### General cell culture practices

HOG cells and 293METR packaging cells were cultured in DMEM with 5% FBS, 5% GlutaMAX, and 0.1% penicillin-streptomycin at 37°C and 5% CO_2_. Stocks of all stable cell lines are retained, frozen in Bambanker freezing medium and stored in liquid nitrogen.

### Cloning and construct generation

Constructs for purification of the MOG ectodomain (pMS192 (WT), pMS193 (N60A), pMS194 (N60D)) were synthesized by Twist Bioscience in the pTwist CMV backbone. MOG overexpression constructs were synthesized by Twist Bioscience with pMS156 in the pTwist Lenti CMV backbone and pMS184, pMS185, pMS188, pMS190, and pMS191 in the pTwist Lenti SFFV backbone. pMS156 uses the β1 isoform while the remaining plasmids use the α1 isoform. pMS156 encoded mScarlet-i–T2A–MOG–3×FLAG, allowing mScarlet-i and MOG to be expressed as separate proteins. The remaining constructs encoded direct C-terminal EGFP fusions comprising full-length human MOG (MOG-EGFP; pMS184), a version of MOG with the cytoplasmic tail and the second transmembrane region deleted (MOG-Δcyt-EGFP; pMS185), an extracellular-domain-deletion construct used as the Membrane-EGFP negative control (pMS188), and full-length MOG containing the N60A (MOG-N60A-EGFP; pMS190) or N60D (MOG-N60D-EGFP; pMS191) substitution. N-glycan processing enzyme overexpression constructs (pMS085 (empty vector), pMS158 (*HEXA*), pMS160 (*MGAT1*), pMS161 (*MGAT2*), pMS163 (*MAN2A2*), pMS164 (*ST6GAL1*)) were assembled using In-Fusion HD cloning. Gene fragments containing the various enzyme coding sequences (and the empty vector) were inserted into a lentiviral overexpression vector containing a CAG promoter region and an IRES2-linked puromycin-resistance–BFP selection cassette. N-glycan processing enzyme knockdown constructs (pMS167 (*HEXA*), pMS168 (*HEXB*), pMS170 (*MGAT1*), pMS172 (*MGAT2*), pMS174 (*MAN2A1*), pMS176 (*MAN2A2*), pMS178 (*ST6GAL1*), pMS180 (*FUT8*), and pMS182 (non-targeting control)) were generated by ligating annealed oligonucleotides encoding the indicated sgRNA protospacers into the BstXI and BlpI sites of the pLG1 vector. The dCas9-KRAB construct (pCL0029) was a generous gift from Luke Gilbert’s lab^105^. The pMD.G and pCMVΔR8.91 packaging plasmids were used for lentivirus production.

### Cell line generation

HOG (human oligodendroglioma) cells were used as the parental background for all cell culture experiments. For experiments examining N-glycan processing enzymes, we first established two stable working populations. HOG cells were transduced with pMS156, and mScarlet-i-positive cells were isolated by fluorescence-activated cell sorting (FACS) to generate the background used for enzyme overexpression cell lines. Separately, HOG cells were transduced with pCL0029 to generate cells expressing dCas9-KRAB and BFP. These were then transduced with pMS156 and sorted for both BFP and mScarlet-i expression. This double-positive population was used for knockdown CRISPRi experiments.

Stable knockdown lines were generated by transducing the MOG- and dCas9-KRAB-expressing HOG population with sgRNA constructs targeting *HEXA* (pMS167), *HEXB* (pMS168), *MGAT1* (pMS170), *MGAT2* (pMS172), *MAN2A1* (pMS174), *MAN2A2* (pMS176), *ST6GAL1* (pMS178), or *FUT8* (pMS180), or with a non-targeting control sgRNA (pMS182). Following transduction, cells were selected with 1.25 or 2.5 µg/mL puromycin and sorted to retain the mScarlet-i-positive, MOG-expressing population.

Stable overexpression lines were generated by transducing the MOG-expressing HOG population with constructs encoding *HEXA* (pMS158), *MGAT1* (pMS160), *MGAT2* (pMS161), *MAN2A2* (pMS163), or *ST6GAL1* (pMS164), or with the corresponding empty-vector control (pMS085). Cells positive for the BFP selection marker and retaining mScarlet-i expression were isolated by FACS and expanded. Despite repeated attempts, viable populations constitutively overexpressing *HEXB, MAN2A1*, or *FUT8* could not be established.

The stable lines used for MOG cell-based assays were generated separately in unmodified parental HOG cells. Cells were transduced with constructs encoding full-length MOG-EGFP (pMS184), MOG-Δcyt-EGFP (pMS185), Membrane-EGFP (pMS188), MOG-N60A-EGFP (pMS190), or MOG-N60D-EGFP (pMS191). Cells were sorted using a narrow fluorescence window across all five populations to generate pooled cell lines with comparable reporter expression.

For all lines, VSV-G-pseudotyped lentivirus was produced in 293METR packaging cells using pMD.G and pCMVΔR8.91. Viral supernatants were collected 48 h after transfection, passed through a 0.45-µm low-protein-binding filter, and concentrated twentyfold using an Amicon Ultra-15 concentrator (EMD Millipore) with a 100,000 Da molecular mass cutoff. HOG cells were spinfected in 1.5 mL DMEM containing 15 mM HEPES and 8 µg/mL polybrene, incubated overnight at 37°C, and returned to fresh growth medium the following day.

### MOG cell-based assays for antibody binding

HOG cells stably expressing MOG-EGFP (pMS184), MOG-Δcyt-EGFP (pMS185), Membrane-EGFP (pMS188), MOG-N60A-EGFP (pMS190), or MOG-N60D-EGFP (pMS191) were used for cell-based binding assays (CBAs) to assess antibody binding to MOG on the cell surface. Different variations of the assay were performed, with key shared steps.

#### Shared steps

Cells were seeded 3 days before use in the CBA and grown to 100% confluency or more, with media replaced the day before harvesting. On the day of the assay, cells were checked by eye for viability, then were washed with PBS (2 mL), lifted with TrypLE (1 mL per 10 cm dish or 2 mL per 15 cm dish), and resuspended in four volumes of complete medium to quench. Cells were counted (10 µL suspended cells + 10 µL Trypan Blue Solution 0.4%, TC20 Automated Cell Counter by Bio-Rad Laboratories; two counts per suspension). Equal cell numbers were used for every condition on a plate. Cells were pelleted (400 × g, 3 min) and resuspended in PBE (PBS with 0.5% BSA and 2 mM EDTA) at a density that gave the required cell number in 100 µL (for example 2.5 × 10^6^ cells/mL for 2.5 × 10^5^ cells per well). Cells were placed in 96-well V-bottom plates on ice and washed three times with 200 µL PBE (400 × g, 1 min, 4°C). Cells were then incubated with 100 µL of diluted plasma or CSF for 1 h at 4°C, washed three times, and incubated with 100 µL of Alexa Fluor 647-conjugated anti-human IgG (1:1,000 in PBE) for 30 min at 4°C. After three final washes, cells were resuspended in PBE and measured on a MACSQuant Analyzer 10 (Miltenyi Biotec) in fast mode with the same instrument settings for all runs and on a cooling block. Each plate included an unstained control (PBE only) and a positive control, the mouse anti-MOG antibody 8-18C5 (1:1,250)^106^. Alexa Fluor 647-AffiniPure Goat Anti-Human IgG, Fcγ Fragment Specific (1:1,000; Jackson ImmunoResearch 109-605-008) was used for all plasma and CSF wells and Alexa Fluor 647-conjugated AffiniPure Donkey Anti-Mouse IgG (1:1,000; Jackson ImmunoResearch 715-605-150) only for 8-18C5 wells. Plasma, CSF, and antibodies were thawed on ice, and primary dilutions were made fresh for each run with 4°C PBE (plasma dilutions at 1:25, CSF dilutions at 1:2).

#### Initial screen for MOG binders

Plasma was diluted 1:25 in PBE and tested on cells expressing MOG-EGFP (pMS184) or Membrane-EGFP (pMS188), at 2.5 × 10^5^ cells per well. Membrane-EGFP cells served as the negative control, using only the MOG transmembrane region to anchor EGFP to the cytosolic side of the membrane. Binding to MOG-EGFP was expressed as fold change over Membrane-EGFP. A sample was called positive at a fold change > 2.5, in line with the threshold used in clinical MOG-IgG CBAs^53^. In this initial screen, two technical replicates were performed for each of the 462 plasma samples.

#### Comparison across MOG mutants

Plasma (1:25 dilution) was tested in parallel against the five cell lines. 3–4 biological replicates were performed for each ADRC Draw 1 sample. For ADRC Draw 2 measurements shown in Figure 5G, a single replicate was performed.

#### Pooled format for CSF

Because CSF contains much less IgG than plasma and volumes were limited, a mixed-well format was used. Untransduced (parental) HOG cells were mixed 1:1 with MOG-EGFP (pMS184), giving 50,000 total cells per well. CSF was diluted 1:1 in PBE. In each well, MOG-EGFP cells were identified as EGFP-positive and parental cells as EGFP-negative, so the negative control saw the same CSF, secondary antibody, and staining conditions as the test cells. Binding was expressed as the AF647 median of MOG-EGFP cells divided by that of parental cells in the same well. Three biological replicates were performed.

#### Blocking N-glycosylation with NGI-1

To alter N-glycosylation in a second, independent way, MOG-EGFP (pMS184) and Membrane-EGFP (pMS188) cells were treated with NGI-1, a potent inhibitor of the oligosaccharyltransferase complex. NGI-1 (MedChem Express) was kept as a 40 mM stock in DMSO and diluted to 10 mM in DMSO (5 µL stock + 15 µL DMSO) on day of treatment. For each 10 cm dish, 20 µL of this dilution was added to 20 mL of complete medium (10 µM NGI-1, 0.1% DMSO). Control dishes received 20 µL DMSO in 20 mL medium (0.1% DMSO). Cells were resuspended in the drug-containing medium, plated, and cultured for three days before the assay. Because NGI-1 slows growth, NGI-1-treated cells were seeded at twice the density of DMSO-treated cells, so that both reached equivalent confluence on the assay day. Untreated stock plates were kept in parallel. On the assay day, cells were processed as described above, with 2.5 × 10^5^ cells per well and plasma diluted 1:25. Three biological replicates were performed.

#### Flow cytometry and analysis

All CBAs were analyzed via flow cytometry using a MACSQuant Analyzer 10. All cell sorting was performed on a Sony SH800S cell sorter. All data were exported as .fcs files and analyzed in FlowJo version 10.10.1.

#### MOG ectodomain expression and purification

His-tagged wild type, N60A, and N60D MOG ectodomains were expressed from pMS192, pMS193, and pMS194, respectively, in Expi293 cells. For each construct, 250 mL of cells at approximately 2 × 10^6^ cells/mL were transfected with 250 µg plasmid DNA using ExpiFectamine. DNA and ExpiFectamine were separately diluted in 12.5 mL Opti-MEM, combined after 5 min, incubated for 25 min at room temperature, and added dropwise to the cells. 1.25 mL of Enhancer 1 and 12.5 mL of Enhancer 2 were added 22 h later. Clarified medium was collected approximately 48 and 72 h after transfection, and the two collections were combined. Cells and residual debris were removed by centrifugation at 1,000 × g for 5 min.

MOG ectodomains were purified from the clarified media by batch binding to HisPur Ni-NTA resin. For each preparation, 4 mL resin was equilibrated by three washes with nickel wash buffer containing 20 mM HEPES, pH 7.5, 150 mM NaCl, and 40 mM imidazole. The equilibrated resin was added to the clarified media and incubated for 1 h at room temperature with gentle rotation. Resin was recovered by centrifugation at 400 × g for 5 min and washed three times with 10 mL nickel wash buffer. Bound protein was eluted through four sequential 5 min incubations with 2.5 mL nickel elution buffer containing 20 mM HEPES, pH 7.5, 150 mM NaCl, and 500 mM imidazole. The four eluates were pooled, assessed by SDS-PAGE, and concentrated to approximately 10-fold using a centrifugal concentrator with a 3.5 kDa molecular weight cutoff (MWCO). Samples were dialyzed overnight at 4°C against 2 L of 20 mM HEPES, pH 7.5, and 150 mM NaCl using 3.5 kDa MWCO dialysis cassettes. Purified proteins were aliquoted, flash-frozen in liquid nitrogen, and stored at −80°C.

### MOG ectodomain characterization

Purified WT, N60A, and N60D MOG ectodomains were assessed by SDS-PAGE on 4–12% Criterion XT Bis-Tris gels using MOPS running buffer and a Chameleon Duo prestained protein ladder. A 100 µL aliquot of purified WT MOG ectodomain was further analyzed by size-exclusion chromatography on a Superdex 200 Increase 10/300 GL column using an ÄKTA FPLC system. Protein elution with SEC buffer (20 mM HEPES, pH 7.5, and 150 mM NaCl) was monitored by absorbance at 280 nm.

### HEXB paucimannosylation assays

Purified wild type human MOG ectodomain (2.2 µL, corresponding to approximately 4 µg protein) was combined with recombinant mouse HEXB (MedChemExpress, HY-P75808; 4 µL, corresponding to approximately 4 µg protein) and brought to a final volume of 10 µL with buffer (20 mM HEPES, pH 7.4, and 150 mM NaCl). Reactions were incubated at room temperature for 0 min, 20 min, 60 min, 5 h, or 24 h. Five total replicate reactions were performed for each timepoint. At completion, an equal volume (10 µL) of lysis buffer (4% SDS, 20 mM HEPES, pH 7.4, 150 mM NaCl, and 2× protease inhibitor) was added to each sample and samples were flash frozen in liquid nitrogen and stored at −80°C until joint preparation of all samples for mass spectrometry.

### Tissue sectioning for MALDI–MSI

FFPE brain blocks were sectioned sagittally at a thickness of 5 µm and mounted on glass slides, with one section from each of up to three different blocks placed on the same slide. The 32 sections were distributed across 11 groups: Group 1 (3A, 14A, 25A), Group 2 (3B, 14B, 25B), Group 3 (3C, 14C, 25C), Group 4 (3D, 14D, 25D), Group 5 (3E, 14E, 22F), Group 6 (9A, 18A, 22A), Group 7 (9B, 18B, 22B), Group 8 (9C, 18C, 22C), Group 9 (9D, 18D, 22D), Group 10 (9E, 18E, 22E), and Group 11 (3F, 22G).

### MALDI–MSI of N-glycans

Formalin-fixed paraffin-embedded sagittal brain sections were obtained from 32 male C57BL/6JN mice at 3 months (n = 6), 9 months (n = 5), 14 months (n = 5), 18 months (n = 5), 22 months (n = 7), and 25 months (n = 4). Brain sections were dewaxed in fresh xylene twice for 3 min before being washed and rehydrated as follows: ethanol for 1 min, fresh ethanol for 1 min, Carnoy’s fluid (60% ethanol, 30% chloroform, and 10% acetic acid) for 2 min, ethanol for 1 min, 95% ethanol for 1 min, 70% ethanol for 1 min, water for 3 min, and fresh water for 3 min. Sections underwent antigen retrieval in 100 mM Tris, pH 9.0, for 20 min at 95°C in a Biocare Medical NxGen Decloaking Chamber. After cooling for 15 min, the sections were buffer exchanged into water over five changes and thoroughly dried.

Sections were coated over 15 passes with 0.1 mg/mL PNGase F (Bulldog Bio) in 20 mM ammonium bicarbonate using an HTX M5 Robotic Reagent Sprayer with a flow rate of 0.025 mL/min, track speed of 1,200 mm/min, track spacing of 3 mm, nozzle temperature of 45°C, nozzle height of 40 mm, N_2_ pressure of 10 psi, and a crisscross track pattern. Slides were incubated in a humidity chamber for 135 min at 37°C. After PNGase F digestion, the sections were dried and coated over three passes with 10 mg/mL α-cyano-4-hydroxycinnamic acid in 70% acetonitrile, 0.1% trifluoroacetic acid, and 10 mM ammonium phosphate using a flow rate of 0.120 mL/min, track speed of 1,200 mm/min, track spacing of 3 mm, nozzle temperature of 75°C, nozzle height of 40 mm, N_2_ pressure of 10 psi, and a horizontal-horizontal track pattern.

Mass spectrometry images of released N-glycans were collected in positive-ion mode with TIMS off at 20 µm resolution using a Bruker timsTOF fleX QTOF mass spectrometer. Acquisition parameters were an *m/z* range of 700–3,500, Funnel 1 RF of 450.0 Vpp, Funnel 2 RF of 500.0 Vpp, Multipole RF of 500.0 Vpp, Collision Energy of 10.0 eV, Collision RF of 2,700.0 Vpp, Quadrupole Ion Energy of 5.0 eV, Transfer Time of 140.0 µs, and Pre Pulse Storage of 4.0 µs. A total of 150 laser shots were summed per pixel.

Images were root mean square (RMS) normalized and visualized using SCiLS Lab Pro 2026b. *m/z* features were manually selected using an integration window of ±15 ppm. Putative glycan composition assignments were generated using the MetaboScape 2025 plugin for SCiLS. H&E images of all 32 brain sections were manually annotated in Hamamatsu NDP.view2 software for eight regions: cerebellum, cerebellar granule layer, hindbrain, hippocampus, hippocampal granule layer, corpus callosum, cortex, and olfactory bulb. Annotations were imported into SCiLS for region-specific quantification. SCiLS exported the average peak-area intensity for the whole section and each annotated region from the RMS-normalized data.

Average peak-area intensities were analyzed in R. An *m/z* feature was retained for statistical analysis when its mean RMS-normalized intensity across all animals was at least 5 in at least one anatomical region or the whole brain. Once selected, the feature was retained in every region. This criterion retained 165 *m/z* features. For each region and *m/z* feature, the 9-, 14-, 18-, 22-, and 25-month groups were compared separately with the 3-month group using two-sided Welch t tests on log_2_(intensity + 1).

### Hematoxylin and eosin (H&E) staining and image registration

The same sections as used for MSI were H&E stained using a modified Carazzi method. Briefly, MALDI matrix was removed by submerging the slides in 100% ethanol for 1 min. Sections were rehydrated with graded ethanol (95% and 70%) followed by water for 30 sec each. Sections were then stained in hematoxylin for 2 min before excess stain was removed by submerging in water, 70% and 95% ethanol for 30 sec each. Sections were counterstained in Eosin Y with phloxine B for 1 min before being dehydrated in 95% and 100% ethanol for 30 sec each and xylene for 2.5 min and finally coverslipped. Histologic images were obtained for the 32 brain sections used for MALDI-MSI of N-glycans using a Hamamatsu NanoZoomerSQ Digital Slide Scanner at 20X magnification. Eight regions were manually annotated in Hamamatsu NDP.view2: cerebellum, cerebellar granule layer, hindbrain, hippocampus, hippocampal granule layer, corpus callosum, cortex, and olfactory bulb. The histologic images and region annotations were registered to the corresponding MALDI images in SCiLS Lab Pro 2026b and used to extract region-specific signal intensities.

### Myelin extraction

Previously purified myelin from cohorts 4 and 5 used for proteomics was also used for glycopeptide enrichment. Myelin enrichment was therefore performed previously as described^47^. Glycopeptide enrichment followed our standard protocol.

### PGC–LC–MS for native N-glycan structural analysis

Samples were cut from FFPE blocks into forebrain and hindbrain sections then further cut into approximately 2 mm sections. Samples were warmed at 55°C for 20 min. Xylene was then added to the Eppendorf tube, samples were vortexed and incubated for 10 min. Samples were centrifuged, xylene was removed, and this was repeated a total of three times. 100% ethanol was then added to the samples and incubated for 10 min. Samples were centrifuged, ethanol was removed and the process was repeated with 95% ethanol and 70% ethanol. Final ethanol wash was removed and sample was resuspended in lysis buffer (5% SDS, 50 mM TEAB, pH 7.1).

Mouse brain samples were homogenized using a dounce homogenizer and suspended in lysis buffer, then probe sonicated for additional homogenization. Lysate was denatured using 10 mM dithiothreitol (DTT) at 56°C for 45 min. Following reduction, samples were cooled to room temperature and acidified with 1.2% phosphoric acid. Acidified protein was diluted with S-trap binding buffer (90% methanol, 100 mM TEAB, pH 7.1) and applied to a 96-well S-trap plate (Protifi, part number C02-96well-1) following manufacturer’s protocol with some modifications, as outlined here. Once loaded onto the S-trap, samples were incubated with 50 mM ammonium bicarbonate and 1 µL PNGase F (New England Biolabs) overnight at 37°C. Released N-glycans were eluted with 0.1% trifluoroacetic acid (TFA). Released N-glycans were then desalted via Hypersep carbon SPE. Glycan samples were then dried via SpeedVac. Dried samples were resuspended in 30 µL of water for mass spectrometry (MS) analysis in negative ion mode.

MS data were collected on a Thermo Fisher TSQ Altis QQQ MS coupled to a Vanquish liquid chromatography (LC) system. Samples were separated using a PGC LC column (100 mm length, 1 mm ID, 3 µm particle size) with mobile phase A containing 5 mM HFIP and 5 mM butylamine and mobile phase B containing 5 mM HFIP, 5 mM butylamine, and 60% acetone. An 80 min gradient for N-glycans utilized 0–37% mobile phase B over 70 min, held at 99% B for 5 min, then held at 0% B for 5 min.

A targeted MS/MS method was developed prior to analysis using a library of over 200 N-glycan standards. An SRM method for collision energy optimization at 9 CE steps from 10 to 90 V was performed in Skyline, and isomer mixtures of the N-glycan standards were acquired and manually analyzed to determine three quantitative and/or qualitative product ions per structure^107,108^. 429 unscheduled SRM transitions were utilized. Data were analyzed using Thermo Freestyle software and Skyline software. Specifications for analysis were based on presence of all three product ions, as well as dotp values greater than 0.8 within Skyline. Glycan analysis was performed at Beth Israel Deaconess Medical Center—Glycomics Core (RRID:SCR_024818).

### Protein lysate preparation

Solid tissues from cohorts 1, 4, 5, and 6 were thawed on ice and minced in lysis buffer (2% SDS containing 1× cOmplete protease inhibitor cocktail, 1 mM MgCl_2_, and 0.5 U/µL benzonase in PBS). Samples were then sonicated for 15–25 cycles (3 s on and 3 s off at 50% amplitude). Lysates were clarified at 21,000 × g for 10 min at 18°C, supernatant removed to a fresh tube, frozen, thawed, and clarified again under the same conditions. Plasma samples were adjusted to 2% SDS, 1× cOmplete protease inhibitor cocktail, 1 mM MgCl_2_, consistent with other tissue samples. For cell culture experiments, cell pellets were thawed on ice and resuspended in cell lysis buffer (1% Igepal containing 1× cOmplete protease inhibitor cocktail, 2 mM MgCl_2_, and 0.5 U/µL benzonase in PBS) then incubated with shaking at 4°C for 30 min. Lysates were then clarified at 21,000 × g for 15 min at 4°C and supernatant removed to a fresh tube. 10% SDS was then spiked in to bring samples to a final 2% SDS concentration. Protein concentrations for all lysates were measured by BCA assay.

### Glycoproteomics sample preparation

For each sample, 2 mg protein was processed for the five-organ atlas (cohort 1), 800 µg for the myelin datasets (cohorts 4–5), and 1 mg for the EAE (cohort 6) and HOG knockdown and overexpression datasets. Samples were brought to an equivalent volume of 500 µL in 2% SDS then mixed with 1 mL of SP3 binding solution (Sera-Mag beads in 55% absolute ethanol, 3.75% formic acid). Sera-Mag beads were a 1:1 mix of carboxylate-modified hydrophilic particles (Cytiva, 45152105050250) and carboxylate-modified hydrophobic particles (Cytiva, 65152105050250). To prepare the Sera-Mag bead mix, bead slurry was removed from stock bottles and warmed to room temperature, combined, then washed in water and added to the SP3 binding solution. The final amount of beads used per sample was proportional to the amount of sample (approximately 16.7 µL of each original bead stock per mg of protein). Samples were then incubated with shaking for 15–30 min to allow protein binding to the beads then captured on a magnetic rack. The beads were washed four times with 70% ethanol and once with 100 mM HEPES. Bead-bound proteins were digested overnight at 37°C with trypsin at a 1:25 trypsin-to-protein mass ratio in 100 mM HEPES buffer containing 5 mM TCEP and 20 mM chloroacetamide. Digested peptides (no longer bound to the beads) were recovered from the tube using the magnetic rack to separate away the magnetic beads, followed by a second elution with 2% DMSO and a repeated use of the magnetic rack. An aliquot was retained for total proteome analysis.

The peptides were then dried using a SpeedVac, resuspended in 0.05% formic acid, adjusted to approximately pH 2, and desalted using a high-capacity Oasis PRiME HLB 96-well µElution Plate (30 mg sorbent per well; Waters; 186008054). Plates were conditioned with 80% acetonitrile and 0.05% formic acid, washed with 0.05% formic acid, and then samples were loaded. Bound peptides were washed with 0.05% formic acid then peptides were eluted with 80% acetonitrile and 0.05% formic acid twice. Desalted peptides were dried and resuspended at approximately 5 mg/mL in glycopeptide enrichment buffer (50 mM carbonate-bicarbonate buffer, pH 10.5, 50% ACN). Glycopeptides were then incubated with shaking for 1 h at room temperature with Bondesil-PBA beads at a 1:2.5 peptide-to-bead mass ratio in a 96-well deep block plate^36^. The mixture (with glycopeptides bound to beads and peptides free in solution) was then transferred to a 0.45 µm filter plate (EMD Millipore MSHVN4550), centrifuged to remove unbound material, and washed seven times with glycopeptide enrichment buffer. Glycopeptides were then eluted twice for 15 min with 1% trifluoroacetic acid in 50% acetonitrile. Enriched glycopeptides were dried, desalted again using low-capacity Oasis plates (3 mg sorbent per well; Waters; 186008052), dried, and resuspended in 0.2% formic acid before LC–MS analysis.

For analysis of MOG glycosylation after HEXB treatment, a scaled-down protocol was used, assuming approximately 10 µg of protein content available for the initial bead binding steps. After trypsin digest, the large-scale Oasis desalting and glycopeptide enrichment steps were skipped, moving directly to the low-capacity Oasis desalting step. As this experiment used a known purified glycoprotein and we wanted to examine all peptides, regardless of glycosylation status, no enrichment step was necessary or performed.

### TMT proteomics sample preparation

Whole proteome peptide aliquots reserved during glycoproteomics sample preparation were used as input for TMT-based proteomics. 10 µg of peptide per sample was dried and resuspended in 10 µL water. TMTpro 18-plex reagents were dissolved in fresh acetonitrile at 0.5 mg per 68.75 µL, and 10 µL reagent was added to each sample. Labeling proceeded for 1 h at room temperature with shaking at 750 rpm and was quenched with 4 µL of 5% hydroxylamine for 15 min. The labeled channels from each experimental plex were then pooled and dried via SpeedVac to remove acetonitrile. Samples were then resuspended in 0.05% formic acid, acidified to approximately pH 2, and desalted using low-capacity Oasis plates using the above described protocol.

Each pooled sample was then fractionated using Pierce High-pH Reversed-Phase spin columns (Thermo Fisher Scientific, 84868). Columns were first conditioned with methanol, acetonitrile, and 20 mM ammonium formate, pH 10 and then the resuspended peptide mix was bound to the column. Serial elutions were then performed. Peptides were separated across a 2–100% acetonitrile step gradient in 20 mM ammonium formate, pH 10, producing 48 fractions. Fractions separated by 24 positions were combined to generate 24 final groupings, which were dried and resuspended in 0.2% formic acid before LC–MS/MS analysis.

In cohort 1, 5 organs were collected, each with 54 total samples. Each organ’s samples were therefore split across 3 total plexes, with 2 replicates from each age contained within each plex. All plexes for an organ were prepared simultaneously. For the EAE dataset, the full set of 20 samples were used for the glycoproteomics however 2 samples were dropped to allow the analysis to be performed on a single plex. The two EAE samples with the lowest endpoint severity scores (both with a final score of 1) were therefore excluded. The myelin proteomics datasets were similarly prepared as described previously^47^.

### Mass spectrometry instrument operation

The Orbitrap Fusion Tribrid mass spectrometer was operated using Xcalibur version 4.7.69.37 and Tune version 4.2.4321, and the Q Exactive HF-X mass spectrometer was operated using Xcalibur version 4.6.67.17 and Tune version 2.13 Build 3162 (Thermo Fisher Scientific).

### Glycoproteomics mass spectrometry data acquisition

Glycopeptide-enriched samples from the five-organ atlas, the myelin cohorts, and the HOG cell knockdown and overexpression experiments were analyzed using an UltiMate 3000 RSLCnano system coupled to an Orbitrap Fusion Tribrid mass spectrometer (Thermo Fisher Scientific) with an EASY-Spray ion source. The chromatography system used an Acclaim PepMap 100 C18 trapping cartridge (5 µm, 100 Å, 300 µm × 20 mm; Thermo Fisher Scientific) and an Aurora Series AUR3-25075C18-TS analytical column (25 cm × 75 µm internal diameter, 1.7 µm C18; IonOpticks). Mobile phase A was 0.2% formic acid in water and mobile phase B was 0.2% formic acid in acetonitrile. Peptides were loaded at 5 µL/min and separated at 300 nL/min with the analytical column maintained at 45°C. The 120 min method held 2% B through 6 min, increased to 5% B at 6.5 min and 28% B at 96.5 min, increased to 85% B at 101 min and held through 105 min, decreased to 50% B at 105.1 min and 2% B at 110 min, and re-equilibrated through 120 min. For each sample, 5 µL was injected twice from an initial total sample volume of 12 µL.

The Orbitrap Fusion was operated in positive-ion data-dependent acquisition mode. The spray voltage was 1.85 kV, the ion transfer tube temperature was 325°C, and the RF lens was set to 60%. Full MS scans were acquired from m/z 500–1,800 at 120,000 resolution with an AGC target of 4 × 10^5^ and a maximum injection time of 50 ms. Precursors with charge states 2–8 were selected in 3 s cycles, isolated with a 2.0 m/z window, and fragmented by higher-energy collisional dissociation (HCD) at a normalized collision energy of 35. MS2 spectra were acquired in the Orbitrap from m/z 120–4,000 at 30,000 resolution with an AGC target of 1 × 10^5^ and a maximum injection time of 200 ms. Dynamic exclusion was 30 s with a ±10 ppm mass tolerance.

Glycopeptide-enriched EAE spinal cord samples were analyzed using an UltiMate 3000 RSLCnano system coupled to an Orbitrap Q Exactive HF-X mass spectrometer (Thermo Fisher Scientific) with a Nanospray Flex ion source. The same trapping cartridge and mobile phases were used with a 110 cm µPAC Neo HPLC analytical column (COL-NANO110NEOB; Thermo Fisher Scientific). Peptides were loaded at 20 µL/min and separated at 400 nL/min with the column maintained at 50°C. The 180 min method held 5% B through 6 min, increased to 7% B at 6.1 min, 20% B at 120 min, 28% B at 146 min, and 95% B at 156 min, held 95% B through 165 min, and returned to 5% B at 165.1 min for re-equilibration through 180 min. For each EAE sample, 5 µL was injected twice from an initial total sample volume of 12 µL.

Purified MOG glycoproteomics samples from the HEXB paucimannosylation assay were analyzed using the same HF-X LC–MS/MS configuration and scan method with a 95 min chromatographic method. Samples were loaded at 20 µL/min and separated at 400 nL/min with the column maintained at 50°C. The method held 5% B through 6 min; increased to 7% B at 6.1 min, 11% B at 10 min, 15% B at 40 min, 20% B at 60 min, 28% B at 66.5 min, and 95% B at 71 min; held 95% B through 75 min; and returned to 5% B at 75.1 min for re-equilibration through 95 min. For each purified MOG sample, 5 µL was injected three times from an initial total sample volume of 20 µL.

The HF-X was operated in positive-ion data-dependent acquisition mode. The spray voltage was 1.5 kV, the capillary temperature was 250°C, and the S-lens RF level was 40. Full MS scans were acquired from m/z 300–2,000 at 120,000 resolution with an AGC target of 1 × 10^5^ and a maximum injection time of 256 ms. The 20 most intense precursors were selected, excluding unassigned and singly charged ions, isolated with a 2.0 m/z window, and fragmented by HCD at a normalized collision energy of 34. MS2 spectra were acquired from m/z 200–2,000 at 30,000 resolution with an AGC target of 1 × 10^5^ and a maximum injection time of 200 ms. The minimum AGC target was 1.2 × 10^3^, the precursor intensity threshold was 6 × 10^3^, and dynamic exclusion was 20 s.

### Glycoproteomics data processing

Thermo .raw files were converted to mzML format using ProteoWizard MSConvertGUI version 3.0.25120 with the peakPicking and zeroSamples filters enabled^109^. Data were searched using FragPipe version 22.0 with MSFragger version 4.1, ProteinProphet for protein inference, Philosopher version 5.1.1, PTM-Shepherd version 3.0.0, and IonQuant version 1.10.27^110-113^. Mouse samples were searched against the reviewed *Mus musculus* UniProt reference proteome UP000000589 downloaded on May 1, 2025, supplemented with common contaminants and reversed decoy sequences. HOG cell and purified MOG samples were searched against the reviewed *Homo sapiens* UniProt reference proteome UP000005640 downloaded on July 21, 2025, with the same contaminant and decoy strategy.

MSFragger was run in N-glycan labile-search mode using the preset glyco-N-HCD workflow as a starting point^114^. Specific glycan composition libraries were used depending on the dataset. Mouse datasets were searched using an established 1,038-composition N-glycan library (the Potel library) containing both Neu5Gc and Neu5Ac^36^. HOG knockdown and overexpression samples were searched using a 580-composition library generated from the Potel library by converting each Neu5Gc count to Neu5Ac, combining it with any Neu5Ac already present in the composition, and removing duplicate compositions. Purified MOG samples were searched using a separate 708-composition library (The MSFragger preset Human_N-glycans_large_708 library). Precursor and fragment mass tolerances were ±20 ppm, isotope errors of 0, +1, and +2 were considered, and mass calibration with automatic parameter optimization was enabled. Glycan mass offsets were restricted to asparagine, and b, y, and glycan-retaining Y ions were included. The summed diagnostic-ion intensity was required to be at least 10% of the spectrum base peak. Searches required fully tryptic peptides, allowed two missed cleavages, and considered peptides 7–50 amino acids long and 400–5,000 Da. Carbamidomethylation of cysteine (+57.02146 Da) was fixed; methionine oxidation (+15.9949 Da, up to three per peptide) and protein N-terminal acetylation (+42.0106 Da) were variable, with no more than three variable modifications per peptide.

Protein inference was performed with ProteinProphet, and Philosopher sequential target-decoy filtering controlled PSM, peptide, ion, and protein false discovery rates at 1%. PTM-Shepherd assigned glycan compositions with an 8 ppm precursor tolerance, a 10 ppm spectral tolerance (20 ppm for the purified MOG dataset), isotope errors from −1 to +3, a 50 ppm glycan-mass tolerance, and a glycan false discovery rate of 1%. Glycan delta masses were removed after assignment. The primary outputs were the PTM-Shepherd-annotated PSM tables and the corresponding FragPipe peptide and protein reports; IonQuant label-free intensity tables were also generated. Technical injections belonging to the same biological sample were assigned to the same sample group during processing.

### TMT proteomics mass spectrometry data acquisition

High pH reversed-phase fractions from the five-organ atlas and EAE TMTpro 18-plex experiments were analyzed using an UltiMate 3000 RSLCnano system coupled to an Orbitrap Q Exactive HF-X mass spectrometer with a Nanospray Flex ion source. The chromatography system used an Acclaim PepMap 100 C18 trapping cartridge (5 µm, 100 Å, 300 µm × 20 mm) and a 110 cm µPAC Neo HPLC analytical column (COL-NANO110NEOB; Thermo Fisher Scientific). Mobile phase A was 0.2% formic acid in water and mobile phase B was 0.2% formic acid in acetonitrile. Peptides were loaded at 20 µL/min and separated at 400 nL/min with the column maintained at 50°C. The 185 min method held 5% B through 6.1 min, increased to 7% B at 10 min and 28% B at 156.5 min, increased to 95% B at 161 min and held through 170 min, and returned to 5% B at 170.1 min for re-equilibration through 185 min.

The HF-X was operated in positive-ion data-dependent acquisition mode with a spray voltage of 1.5 kV, capillary temperature of 250°C, and S-lens RF level of 40. Full MS scans were acquired from m/z 350–1,500 at 120,000 resolution with an AGC target of 3 × 10^6^ and a maximum injection time of 50 ms. The 15 most intense precursors were selected, excluding unassigned and singly charged ions, isolated with a 0.7 m/z window, and fragmented by HCD at a normalized collision energy of 31. MS2 spectra were acquired from m/z 200–2,000 at 45,000 resolution with a fixed first mass of m/z 110, an AGC target of 1 × 10^5^, and a maximum injection time of 120 ms. The minimum AGC target was 1.2 × 10^3^, the precursor intensity threshold was 1 × 10^4^, and dynamic exclusion was 30 s. For atlas and EAE proteomics, injection volumes were adjusted by experiment to target approximately 750 ng of peptide per LC–MS/MS injection. Each of the five atlas organs was analyzed using three TMTpro 18-plexes (15 plexes total); each plex was separated into 24 concatenated fractions, and each fraction was analyzed in two LC–MS/MS injections (48 injections per plex). The EAE TMTpro 18-plex was separated into 24 concatenated fractions, and each fraction was analyzed in one LC–MS/MS injection (24 injections for the plex).

### TMT proteomics data processing

Thermo .raw files were converted to mzML format using MSConvertGUI with the peakPicking and zeroSamples filters enabled. All datasets were processed using the same FragPipe version 22.0 workflow with MSFragger version 4.1, Percolator version 3.6.5, ProteinProphet for protein inference, Philosopher version 5.1.1, IonQuant version 1.10.27, and TMT-Integrator version 5.0.9^115,116^. Files were searched against the same reviewed *Mus musculus* UniProt reference proteome as above. Fractions and repeated sample injections were searched jointly within their assigned group. Each dataset (brain, liver, kidney, spleen, plasma, and EAE spinal cord) was processed separately using the TMT16 workflow as a starting point for default parameters. The previously published myelin datasets were reprocessed with the same workflow^47^.

MSFragger searches used precursor and fragment mass tolerances of ±20 ppm with mass calibration and automatic parameter optimization enabled. Isotope errors from −1 to +3 were considered. Searches required fully tryptic peptides, allowed two missed cleavages, and considered peptides 7–50 amino acids long and 200–5,000 Da. Carbamidomethylation of cysteine (+57.02146 Da) and TMTpro labeling of lysine (+304.20715 Da) were fixed. TMTpro labeling of the peptide N terminus (+304.20715 Da), methionine oxidation (+15.9949 Da), and protein N-terminal acetylation (+42.0106 Da) were variable, with no more than three variable modifications per peptide. Peaks from m/z 125.5–134.5 were excluded during sequence matching. PSMs were rescored with Percolator, proteins were inferred with ProteinProphet, and Philosopher target-decoy filtering controlled PSM, peptide, ion, and protein false discovery rates at 1%.

TMTpro 18-plex reporter-ion intensities were extracted from MS2 spectra with IonQuant using a 20 ppm reporter-ion tolerance and the experiment-specific channel annotation files. TMT-Integrator summarized reporter intensities using a minimum PSM probability of 0.90 and minimum precursor isolation purity of 0.50, retained the best PSM, removed outliers, and used the top three peptides for protein-level aggregation. PSM-level and protein-level normalization were disabled at this stage. TMT-Integrator generated the protein abundance and ratio tables used as the inputs for downstream normalization and analysis.

### Quantification and statistical analysis

#### Glycoproteomics post-processing and quality filtering

For each biological sample, we used its FragPipe psm.tsv report to quantify glycosite coverage and glycan composition. PSMs were retained only if they contained a reported glycan composition, had a glycan q value < 0.05, were explicitly marked as unique (Is Unique = TRUE), and were not assigned to a contaminant protein. Inclusion further required exactly one valid N position in Best Positions and Number Best Positions = 1. PSMs with multiple candidate positions, no reported position, or an invalid position were excluded. The glycosite coordinate was calculated from the peptide’s start coordinate in the protein and the localized position within the peptide. Because glycosites were numbered relative to the full UniProt precursor sequence, annotated signal peptides were not removed or renumbered. Accordingly, the sole glycosite of mouse MOG is reported as N59, equivalent to N31 when numbered from the mature protein. Each retained PSM was also required to map to a single protein accession. Glycosites were indexed by UniProt accession and asparagine position.

Within each biological sample, PSMs identifying the same exact glycan composition at the same glycosite were counted together. This produced one PSM count for each sample, site, and exact-glycan combination. Exact glycans were also assigned hierarchically to one of eight mutually exclusive glycan types. Glycans containing phosphate were classified as phospho. Compositions containing two HexNAcs and one to three hexoses without Neu5Ac or Neu5Gc were classified as paucimannose, and compositions containing one or two HexNAcs, no hexose, and no Neu5Ac or Neu5Gc were classified as stubs. Glycans containing both fucose and Neu5Ac or Neu5Gc were classified as fucosylated & sialylated. Glycans containing fucose but no sialic acid were classified as fucosylated, whereas glycans containing Neu5Ac or Neu5Gc but no fucose were classified as sialylated. Compositions containing two HexNAcs and at least four hexoses, without fucose, sialic acid, or phosphate, were classified as high mannose. All remaining compositions were classified as undecorated. PSM counts for exact glycans assigned to the same type were summed within each sample and glycosite.

For each biological sample, the total PSM count at a glycosite was the sum of all retained PSMs assigned to that site, regardless of exact glycan composition. The percent abundance of an exact glycan was calculated by dividing its PSM count by the total PSM count at that site and multiplying by 100. Glycan type percent abundance was calculated in the same way after exact-glycan counts were summed by type. For a glycan or glycan type observed at a given glycosite in at least one sample, samples in which the glycosite was detected but that glycan or type was not observed were assigned 0% abundance. If the glycosite itself was not detected, its abundance was recorded as missing rather than zero.

Global glycan composition was calculated separately for each biological sample. Retained PSMs were summed for each exact glycan across all glycosites and divided by the total number of retained glycoPSMs in that sample. Global glycan type composition was calculated similarly. Mean glycosite depth was calculated by dividing the total number of retained PSMs assigned to the site across the dataset by the total number of biological samples (e.g., 54 in the brain atlas). Sites were assigned to depth tiers as follows: Tier 0, < 5/54 (approximately 0.0926) mean PSMs per sample; Tier 1, ≥ 5/54 to < 0.5; Tier 2, ≥ 0.5 to < 1; Tier 3, ≥ 1 to < 2; Tier 4, ≥ 2 to < 4; Tier 5, ≥ 4 to < 10; Tier 6, ≥ 10 to < 20; and Tier 7, ≥ 20 mean PSMs per sample. Thus, a Tier 0 site in the brain could have 1–4 PSMs across all 54 samples, whereas a Tier 7 site would have at least 1,080 total PSMs across all samples.

#### Proteomics post-processing and quality filtering

Protein-level TMT reporter abundances were obtained from the abundance_protein_None.tsv reports generated by FragPipe and matched by Protein ID to the corresponding FragPipe protein.tsv reports. For each atlas organ, the 54 biological samples were arranged as six nine-channel blocks, with each block containing one animal from each of the nine ages. A protein was retained within a block only when an abundance was reported in all nine channels and the associated group-level protein report contained at least two total spectral counts for that Protein ID. Entries annotated as contaminants in the combined protein report were excluded from downstream atlas eligibility sets. A protein could be retained in some blocks and absent from others; absent values remained missing and were not replaced with zero or imputed. Proteins required support in at least three complete replicate blocks for downstream comparisons. For the EAE dataset, the protein abundance table contained 18 reporter channels representing eight control and ten EAE samples. Proteins were retained only if they had at least two total spectral counts and a positive, finite reporter abundance in all 18 samples. Proteins with any missing reporter abundance were therefore excluded, and no missing values were imputed.

Reporter abundances were log_2_ transformed before normalization. Median normalization was performed independently for each atlas organ and for EAE using a shared protein set with finite log_2_ abundances in every sample of the respective dataset. For each sample, the median log_2_ abundance was calculated from exactly the same shared proteins. A single target median was then calculated from all shared-protein log_2_ abundances across that dataset, and the difference between the target median and each sample median was added to every retained protein value in that sample. Quality-control calculations confirmed that the normalized shared-protein median equaled the target median in every sample. Missing measurements remained missing throughout normalization. For scaled intensity plots, each protein’s normalized abundance was divided by its median abundance across all samples. A scaled intensity of 1 therefore represents the protein’s median abundance, whereas values of 0.5 and 1.5 represent half and 1.5-fold the median abundance, respectively.

#### Proteomic differential-abundance analysis

Differential-abundance analyses used the median-normalized log_2_ protein abundances described in the proteomics post-processing section. Log_2_ fold change was the difference between group mean log_2_ abundances, and groups were compared with a two-sided Welch’s t test, displaying unadjusted p values. To qualify for significance testing, atlas (whole brain, spleen, kidney, plasma, and liver) proteins required complete values for all nine age channels across 3 replicate blocks. Proteins from the myelin and EAE datasets required complete coverage across all replicates.

#### Total variation distance (TVD) calculations and permutation testing

Differences in glycan composition were quantified using TVD. For two compositions, TVD was calculated as half the sum of the absolute differences between the proportions of each of the eight glycan types. All glycan types observed in either composition were included, with types absent from a detected glycosite assigned a percent abundance of zero. TVD ranges from 0 for identical compositions to 100 for compositions with no shared glycan types.

TVD was calculated in several ways depending on the analysis. For pooled comparisons, glycoPSM counts were summed across the relevant samples before glycan type proportions and TVD were calculated. These calculations weigh each PSM equally regardless of replicate source. TVD is used as a metric to compare glycan type compositions 1) between whole datasets, 2) at specific glycosites between two datasets, and 3) at two glycosites within a dataset. Pooled inter-organ glycosite comparisons were classified as conserved (TVD < 20%), partially shifted (20% ≤ TVD < 50%), or strongly remodeled (≥ 50%). For whole-organ comparisons, all retained glycoPSMs within each animal and organ were pooled to generate an animal-specific glycan type composition. TVD was then calculated between matched organs from the same animal.

For comparisons between ages or experimental conditions, glycan type proportions were first calculated separately for each animal. The group composition was the arithmetic mean of the proportions among animals in which the glycosite was detected, giving each measured animal equal weight regardless of its glycoPSM depth. Animals in which the site was not detected were considered unmeasured and were not assigned a composition. TVD was then calculated between the two group-mean compositions. Statistical significance was assessed by fixed-size animal-label permutation. For each glycosite, labels were shuffled among animals in which the site was measured while preserving the original numbers of detected animals in the two groups. Group-mean compositions and TVD were recalculated after each permutation. The permutation p value was calculated as the number of permuted TVDs greater than or equal to the observed TVD, plus one, divided by the number of permutations plus one^117^. Atlas age-bin comparisons required detection in at least nine animals and at least 18 glycoPSMs in each group; myelin comparisons required at least three animals and six glycoPSMs per group; and EAE comparisons required at least five animals and ten glycoPSMs per group. Atlas analyses used 99,999,999 permutations, EAE analyses used 999,999 permutations, and myelin analyses used 9,999 permutations.

#### Relationship between protein abundance and glycan type remodeling

For each organ, protein abundance change was calculated as the mean normalized log_2_ abundance in old animals (19, 22, and 28 months) minus that in young animals (1, 3, and 6 months). Glycosites were included if they were a hit in any age comparison (old versus young, old versus middle-aged, or middle-aged versus young). For each glycan type, the plotted glycosite change was its mean percentage abundance in old animals minus its mean percentage abundance in young animals. When a protein contained multiple included glycosites, their percentage-point changes were averaged so that each protein contributed one value per glycan type. Associations between protein abundance change and glycan type change were evaluated using two-sided Spearman correlation.

### Glycosite heterogeneity metrics

We adapted a recently described Shannon entropy framework for quantifying N-glycosylation heterogeneity from glycoPSM distributions^44^. Calculations were performed at the glycan type level using the eight defined glycan types. For each detected glycosite, the proportion of glycan type *i* (*p*_*i*_) was its glycoPSM count divided by the total glycoPSM count at that site. The observed number of glycan types was the total number of glycan types supported by at least one glycoPSM (a minimum of 1 and a maximum of 8). Shannon entropy (H) was then calculated as *H = −∑ p*_*i*_ *ln(p*_*i*_*). H* is zero when all PSMs belong to one glycan type and increases as PSMs become more evenly distributed among multiple types.

To express this diversity on a more intuitive scale, we calculated exp(H), which we refer to as weighted glycan types. This value is 1 when a site contains only one glycan type and equals the observed number of types when all are equally abundant. When the distribution is uneven, the value falls between these limits and is lower when one type predominates.

For sample-level analyses, these metrics were calculated separately for each biological sample. For pooled analyses, glycoPSM counts were first summed across the specified samples, after which the proportions, H, and weighted glycan types were recalculated. A glycan type absent from a detected site was assigned a proportion of zero, whereas no heterogeneity value was assigned when the glycosite itself was not detected. For inter-organ analyses, sites were categorized as having low (< 1.5 weighted glycan types), intermediate (1.5 to < 2.5), or high (≥ 2.5) heterogeneity.

#### Within-protein and structure-distance analyses

Within each organ, TVD was calculated for every unique pair of Tier 2+ glycosites. Pairs were assigned to nonoverlapping within-protein and between-protein groups, and their TVD distributions were compared using a two-sided Wilcoxon rank-sum test.

Within-protein pairs were then used to test whether glycosites that are closer in three-dimensional space have more similar glycan type compositions. Each within-protein pair was matched to a structure spanning both glycosites. Experimentally determined mouse structures in the Protein Data Bank (PDB) were preferred over predicted structures. Candidate PDB structures were ranked with X-ray structures first, followed by lower reported resolution and greater mapped sequence coverage. UniProt positions were mapped to PDB residues using SIFTS^118^. If no experimental structure yielded a usable measurement, the corresponding AlphaFold Database model was used^119^. Across the Tier 2+ analysis, the measurement stage used 85 distinct PDB structures and 493 distinct AlphaFold models. After source-specific quality filtering, 2,659 organ-specific pair measurements were retained: 858 derived from PDB structures and 1,801 from AlphaFold models.

For each selected structure, a surface mesh was constructed after expanding atomic van der Waals radii by a 1.40 Å probe radius; mesh vertices were spaced at approximately 1.35 Å. Each glycosite was anchored to the mesh vertex nearest the glycan-bearing side-chain nitrogen (ND2) of its asparagine. The 5.5 Å anchor cutoff was a maximum tolerance for mapping the atomic coordinate to the discretized solvent-accessible surface. The median ND2-to-anchor offset was 2.88 Å and the maximum was 3.66 Å. Two measures of distance were then calculated. Direct distance was the Euclidean distance between the two surface anchors, whereas surface distance was the shortest path between them along the mesh. Measurements were eligible only when both surface anchors were within 5.5 Å of ND2 and both distance values were finite. For PDB structures, reported resolution had to be 4 Å or better when available, and both glycosites had to be at least five residues from the boundary of the SIFTS-mapped UniProt segment.

AlphaFold measurements required pLDDT ≥ 70 at both glycosites and predicted aligned error (PAE) ≤ 10 Å between them^120^. At least 70% of the surface path had to be supported by residues with pLDDT ≥ 70. Residues with pLDDT < 50 could account for no more than 10 Å of the path in total, and no continuous low-confidence segment could exceed 8 Å. No more than 20% of the path could lie within 5 Å of a residue with pLDDT < 50, and the 90th percentile of the PAE between path-supporting residues and the two glycosite endpoints had to be ≤ 15 Å.

Using the retained direct-distance measurements, median TVD was recalculated as progressively more distant pairs were included, using successive 10 Å upper limits from < 10 to < 90 Å followed by an estimate containing all measured pairs. Exact nonparametric 95% confidence intervals for each median were calculated from order statistics. For a simplified near-versus-far comparison, pairs separated by ≤ 20 Å were compared with pairs separated by > 20 Å using a two-sided Wilcoxon rank-sum test with continuity correction; each glycosite pair was one observation. To determine whether shared peptide evidence contributed to the distance relationship, the analyses were repeated after excluding any pair for which a retained glycopeptide spanned both glycosites.

#### Protein annotations and feature enrichment

Protein annotations were derived from the reviewed *Mus musculus* UniProt proteome downloaded on July 16, 2026. Reviewed UniProt subcellular-location annotations were mapped to the curated location classes used in this study. UniProt feature records were used to annotate signal peptides, transmembrane regions, disulfide bonds, coiled coils, repeats, zinc fingers, lipidation, phosphorylation, acetylation, ubiquitination, and protein domains. Glycoproteins and glycosites were linked to these annotations by UniProt accession. Glycosites were mapped to domains by comparing their protein coordinates with UniProt DOMAIN feature coordinates after excluding isoform-specific domain records.

For organ-level enrichment analyses, a protein was considered detected in an organ when it was supported by at least one retained glycoPSM in that organ. For each curated subcellular-location class and selected UniProt feature, the fraction of detected glycoproteins carrying the annotation was compared with the fraction among all other proteins in the reviewed mouse UniProt proteome using a two-sided Fisher’s exact test. Each protein was counted once in each analysis.

For the atlas-wide protein-domain analysis, glycoproteins detected in at least one organ were compared with all other proteins in the reviewed mouse UniProt proteome for each UniProt DOMAIN annotation using two-sided Fisher’s exact tests. Domains were tested only when at least 10 proteins in the reviewed-proteome background carried the annotation.

The mouse N-glycosylation machinery list was constructed by first assembling candidate genes from the reviewed mouse UniProt proteome, the mouse glycan-related gene set of Nairn et al. (2008), GlycoEnzDB and GlycoEnzOnto commit 8c28a4ba52d1198150884ac174e4546fd096a451, MGI Gene Ontology annotations and mouse-human homology, NCBI Gene, the Gene Ontology, Reactome, and a manually reviewed N-glycosylation-stage reference^121-126^. Candidates identified only in human sources were mapped to mouse orthologs using MGI. Evidence from these sources was then integrated to assign each retained gene to an N-glycosylation stage and to one of three analysis classes. Primary genes directly participate in N-glycan precursor assembly, transfer to protein, processing, quality control, or degradation. Supporting genes supply or transport dedicated precursors or donors, and secondary genes mediate mannose-6-phosphate recognition and trafficking. These three classes defined the N-glycosylation machinery analyses. External NCBI, MGI, Gene Ontology, and Reactome records were retrieved on September 9, 2026.

#### Gene Ontology (GO) enrichment

For Gene Ontology enrichment, glycosites were mapped to proteins by UniProt accession and then to Entrez Gene identifiers, with each gene counted once^125^. In R, one-sided hypergeometric tests were performed separately for Biological Process, Cellular Component, and Molecular Function using clusterProfiler::enricher or the equivalent direct calculation^127^. The background comprised genes that satisfied the same testing and coverage rules as the foreground, and terms required at least five background genes. p values below 0.05 defined significant displayed terms. Annotations were frozen to UniProt mouse reference proteome UP000000589, org.Mm.eg.db version 3.23.0, AnnotationDbi version 1.74.0, GO.db version 3.23.1, and the January 23, 2026 Gene Ontology release. As an example, for Figure 6G, proteins carrying the 23 shared aging-EAE hit sites were tested against the 74 uniquely mapped proteins carrying any of the 97 EAE-hit glycosites.

#### Proportional-odds models of inter-organ remodeling

For each organ pair, Tier 1+ glycosite comparisons were assigned to one of three ordered states based on their TVD: conserved (TVD < 20%), partially shifted (20% ≤ TVD < 50%), or strongly remodeled (TVD ≥ 50%). Each glycosite contributed one equally weighted observation. Each protein or glycosite feature was then tested in a separate proportional-odds logistic-regression model^128^.

The model asks whether a feature shifts glycosites toward higher or lower remodeling states. It evaluates two cumulative boundaries at the same time: conserved versus partially shifted or strongly remodeled, and conserved or partially shifted versus strongly remodeled. The model allowed the two boundaries to have different baseline probabilities by estimating a separate intercept for each boundary, but it estimated only one shared coefficient for the tested feature. Exponentiating this coefficient gives the common odds ratio. This is a joint maximum-likelihood estimate from the complete three-state distribution, not the arithmetic average of odds ratios calculated separately at the two boundaries. For a binary feature, the odds ratio compares sites with and without that feature; values above 1 indicate greater remodeling and values below 1 indicate greater conservation. Binary features required at least 10 sites in both groups. Continuous features were standardized within each organ pair, so their odds ratios represent a one-standard-deviation increase; they required at least five sites and measurable variation.

Models were fit in R with the polr function in the MASS package. To determine significance, we used a two-sided Wald test that tested whether the odds ratio differed from 1. Models were reported as not estimable when the predictor or outcome lacked variation, the data were insufficient, or the log-odds standard error exceeded 2.

For a binary example, the brain-kidney comparison contained 739 glycosites: 454 conserved, 162 partially shifted, and 123 strongly remodeled. Of the 302 sites on proteins annotated as plasma membrane or cell surface, 147 were conserved, 79 were partially shifted, and 76 were strongly remodeled. Of the 437 sites without this annotation, 307 were conserved, 83 were partially shifted, and 47 were strongly remodeled. At the first boundary, 155 of 302 annotated sites (51.3%) and 130 of 437 unannotated sites (29.7%) were partially shifted or strongly remodeled. Their cumulative odds were therefore 155/147 = 1.05 and 130/307 = 0.42, giving a boundary-specific odds ratio of 2.49. At the second boundary, 76 of 302 annotated sites (25.2%) and 47 of 437 unannotated sites (10.8%) were strongly remodeled. Their cumulative odds were 76/226 = 0.34 and 47/390 = 0.12, giving a boundary-specific odds ratio of 2.79.

The proportional-odds model fitted both boundaries to the original six category counts at once rather than averaging the boundary-specific odds ratios of 2.49 and 2.79. The fitted shared log-odds coefficient was 0.94, and exponentiating this coefficient gave a common odds ratio of 2.56 (95% confidence interval, 1.90–3.43; p = 4.57 × 10^-10^). The influence of each boundary on this joint estimate depends on the number and balance of observations that it contributes. Thus, the model estimated that annotated sites had approximately 2.6-fold greater odds of falling above either remodeling boundary than unannotated sites.

For a continuous example, we tested whether high mannose abundance was associated with brain-kidney remodeling. For each of the 739 shared Tier 1+ glycosites, we averaged the site’s high mannose percent abundance in brain and kidney to obtain one site-level value between 0% and 100%. The 454 conserved sites had a mean of 82.1%, the 162 partially shifted sites had a mean of 46.6%, and the 123 strongly remodeled sites had a mean of 39.1%. Thus, higher high mannose abundance was concentrated among sites with more similar glycan compositions in brain and kidney.

The site-level values were standardized across all 739 sites. A site at the overall mean was assigned a standardized value of 0, and a site 35.9 percentage points above the mean was assigned a value of 1 because 35.9 percentage points was one standard deviation. The model retained the value for every site and fitted both remodeling boundaries jointly. A one-standard-deviation increase in mean high mannose abundance, equivalent to 35.9 percentage points on the abundance scale, had an odds ratio of 0.35 (95% confidence interval, 0.30–0.41; p = 9.84 × 10^-40^). Thus, when two sites differed in mean high mannose abundance by 35.9 percentage points, the site with the higher value had 0.35 times the odds of occupying a higher remodeling state, equivalent to a 65% relative reduction in odds. This shared odds ratio applied to both cumulative comparisons: partially shifted or strongly remodeled versus conserved, and strongly remodeled versus conserved or partially shifted.

#### Cosine similarity of glycan type remodeling

Cosine similarity was used to compare the direction of glycan type remodeling between a set of glycosites in two datasets. For each glycosite and dataset, the eight glycan type percentages were first calculated separately for each animal. Group compositions were then calculated as the arithmetic mean across animals in which the site was detected, so each detected animal contributed equally. Two eight-element change vectors were generated for each shared site. Vector components were ordered as stubs, paucimannose, high mannose, phospho, undecorated, fucosylated, sialylated, and fucosylated & sialylated. In the aging versus EAE analysis, the aging vector was the mean composition in old brain minus the mean composition in young brain, and the EAE vector was the mean composition in EAE spinal cord minus the mean composition in control spinal cord. Each element therefore represents the percentage-point change in one glycan type.

Cosine similarity was calculated by dividing the dot product of the two change vectors by the product of their lengths. The resulting value ranges from −1 to 1. A value near 1 indicates that the glycan types increase and decrease in similar relative directions in both comparisons, a value near 0 indicates little overall directional agreement, and a value near −1 indicates opposing remodeling. A cosine similarity of exactly 1 requires one vector to be a positive scalar multiple of the other: every component of one vector must equal the corresponding component of the other multiplied by the same positive constant. Cosine similarity therefore compares direction rather than the overall size of the change.

For example, using the glycan type order above, the ITGAV N74 aging vector was (0, 0, −10.29, 0.35, 1.76, 8.02, 0.17, 0), and the EAE vector was (−0.73, 0, −12.59, 0, 4.22, 9.11, 0, 0). Both vectors were dominated by a loss of high mannose glycans and an increase in fucosylated glycans, but they were not exact positive scalar multiples. For example, the undecorated increase was 2.4 times as large in EAE as with aging, whereas the fucosylated increase was only 1.14 times as large; EAE also produced a small decrease in stubs that was absent with aging. The vectors were nonetheless quite similar overall. Using the rounded vector values above, the nonzero terms in the dot product were (−10.29 × −12.59) + (1.76 × 4.22) + (8.02 × 9.11) = 210.04. The lengths of the aging and EAE vectors were 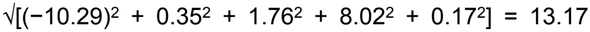 and 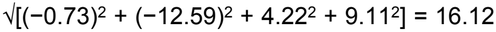, respectively. The cosine similarity was therefore 210.04 ÷ (13.17 × 16.12) = 0.989.

#### Quantification of MOG autoantibody binding

Median Alexa Fluor 647 intensity was used to quantify bound IgG, while median EGFP intensity reported expression of the corresponding MOG-EGFP construct. Different normalization strategies were used depending on the assay design.

For the initial plasma screen, the IgG Binding Index (IBI) was calculated within each replicate as the median Alexa Fluor 647 signal on MOG-EGFP cells divided by the signal on Membrane-EGFP negative-control cells. The replicate ratios were averaged to obtain one IBI for each plasma sample. Samples with a mean IBI > 2.5 were classified as positive candidates. For comparisons among MOG constructs, the Alexa Fluor 647 signal in each well was first normalized to the corresponding EGFP signal to account for variability in MOG levels, producing a normalized IgG binding signal for each MOG construct (what is plotted in Figure 5J). This normalized binding signal for MOG-Δcyt-EGFP, MOG-N60A-EGFP, or MOG-N60D-EGFP was then divided by the normalized IgG binding signal for MOG-EGFP within the matched replicate. This produced an IgG binding ratio relative to MOG-EGFP (what is plotted in Figures 5F–5I). A construct/MOG-EGFP ratio of 1 indicates equal binding, whereas values above or below 1 indicate increased or decreased binding to the alternative construct, respectively.

For the NGI-1 experiment, the antibody signal attributable to MOG was calculated separately under NGI-1 and DMSO treatment by subtracting the Alexa Fluor 647 signal measured on the matched Membrane-EGFP cells from that measured on MOG-EGFP cells. Each background-subtracted value was then divided by the EGFP signal of the MOG-EGFP cells. The effect of NGI-1 was expressed as the normalized signal under NGI-1 treatment divided by the normalized signal under DMSO treatment, and replicate ratios were averaged for each plasma sample.

For CSF measurements, parental HOG and MOG-EGFP cells were mixed in the same well and distinguished by EGFP fluorescence. Similar to the initial plasma screen, the CSF IBI was calculated as the median Alexa Fluor 647 signal of the MOG-EGFP-expressing population divided by that of the EGFP-negative parental population.

#### Clustering of autoantibody binding response patterns

The clustering analysis used the mean MOG-Δcyt-EGFP/MOG-EGFP, MOG-N60A-EGFP/MOG-EGFP, and MOG-N60D-EGFP/MOG-EGFP binding ratios described above. One representative plasma sample from each of 94 patients (10 Young (HC), 28 Old (HC), 26 Old (ND), and 30 MOGAD) was analyzed together with the 8-18C5 monoclonal-antibody control. For each construct, a mean construct/MOG-EGFP ratio below 0.85 was classified as decreased binding, a ratio from 0.85 to 1.15 as unchanged, and a ratio above 1.15 as increased binding. Each sample was assigned to one of the 27 possible three-construct response patterns. The ten patterns observed in the dataset were displayed as clusters. Patterns were ordered from an overall increase toward an overall decrease in binding to the alternative constructs, and samples within each pattern were ordered by their combined N60A and N60D responses followed by their combined response across all three constructs. To calculate significance, we compared binding to each test construct with binding to MOG-EGFP across replicates, applying a two-sided one-sample t test to the log_2_-transformed ratios and using 0 as the null value for equal binding.

#### Glycan shielding

We modeled N-glycans on the crystal structure of the rat MOG extracellular domain (PDB 1PKO, 1.45 Å resolution), the highest-resolution of the four MOG structures in the Protein Data Bank (1PKO, 1PKQ, 1PY9, and 3CSP), none of which carries a glycan^129^. We kept chain A, removed water molecules, kept the first position of side chains with two alternative positions, and converted selenomethionine 35 back to methionine. The original residue numbering was kept, so the glycosylation site is N31 (sequon N31-A32-T33).

Four N-glycan compositions (N2F1, N2H3F2, N4H5F1, N6H7F1) of increasing size, all detected at the MOG glycosite, were modeled with GlycoSHIELD^130^. Glycan conformers were taken from the GlycoSHIELD library, which contains 30,001 conformers per glycan from 3 µs molecular dynamics simulations. Three of the four compositions are in the library. The N2F1 conformers were made by deleting the extra monosaccharides from every conformer of the N2H2F1 library glycan and adding back the hydrogen on the freed hydroxyl group, so their conformations come from simulations of the larger glycan. Every fifth library conformer (6,001 per glycan) was grafted onto N31 by superimposing it on residues 30 to 32. Conformers that clashed with the protein were removed in the coarse-grained mode of GlycoSHIELD with a 3.5 Å cutoff, which rejects a conformer if any sugar ring oxygen lies within 3.5 Å of a protein Cα atom. Between 4,907 and 5,537 conformers were kept per glycan.

For the structural views, 100 evenly spaced accepted conformers of each glycan were shown on the MOG surface in PyMOL (version 3.1.5.1) as sticks with a transparent surface. Glycan atoms were colored by conformer density, defined as the number of glycan atoms within a 12 × 12 × 12 Å cube around each atom, on the same scale for all four glycans. Analyses used Python 3.13, MDAnalysis 2.10.0, NumPy 2.5.3, Matplotlib 3.11.2, and GROMACS 2023.3^131-134^.

## Acknowledgments

We gratefully acknowledge the human research participants. We also acknowledge the sacrifice of the 151 mice used in this research. We thank Project BIG and its biorepository staff, the Stanford Alzheimer’s Disease Research Center clinical and sample-acquisition teams, Stanford Blood Center donors and staff, and Stanford Athlete Study participants, David Benjamin Camarillo, and staff. We also thank Ada, Dalton, Quixote, and Melissa for their guidance and assistance throughout the project. We thank all members of the Wyss-Coray and Bertozzi laboratories for their engagement and feedback during laboratory meetings and small group discussions. We thank Dina Schuster, Sharon Pitteri, and Fernando J. García-Marqués for advice on mass spectrometry topics, and Morgane Boone for guidance on manuscript readability. We acknowledge the Beth Israel Deaconess Medical Center - Glycomics Core (RRID: SCR_024818). Mass spectrometry imaging was performed in the University of Texas at Austin Mass Spectrometry Imaging Core, supported by Cancer Prevention and Research Institute of Texas (CPRIT) Awards RP190617 and RP240559.

M.S. was supported by Schmidt Science Fellows, in partnership with the Rhodes Trust. G.T.-L. was supported by a Stanford Bio-X Undergraduate Summer Research Program grant. S.C. was supported by a Masason Foundation fellowship. D.W. was supported by the National Multiple Sclerosis Society–American Brain Foundation MS Clinician-Scientist Development Award (FAN-2207-39823). T.I. was supported by European Research Council Starting Grant 101163696, Israel Science Foundation grant 1870/24, and Alzheimer’s Association Research Grant AARG-24-1308080. I.H.G. was supported by National Institute on Aging Pathway to Independence Award R00AG088304 and the MAC3 Dementia and Ageing Fellowship. This work was supported by the Phil and Penny Knight Initiative for Brain Resilience and by a Simons Foundation International grant administered by the Simons Foundation (SFI-AN-NC-AB-Research-00018033) (T.W.-C.). Additional support was provided by the Stanford Alzheimer’s Disease Research Center through National Institute on Aging grants P50AG047366 and P30AG066515, the Garrett Immunology Research Fund, and MCHRI-TIP. Funding for Project BIG was provided by the Project BIG Research Fund and the CRUSH MS Foundation. This research was supported in part by a grant to C.R.B. from the National Institutes of Health (CA200423).

## Contributions

M.S., T.W.-C., and C.R.B. conceived the study. The study was designed and led by M.S., who performed experiments and bioinformatic analyses, prepared the figures, and wrote the manuscript. J.R. performed experiments, assisted with analyses, and contributed to manuscript review and editing. Additional experiments were performed by K.A., G.T.-L., S.C., Y.F., L.E.P., T.I., E.H.S., and I.H.G. Under the supervision of M.W., D.W. performed the EAE experiments. Important study samples were obtained with assistance from D.C., J.E.D., M.H.H., I.H.G., and T.I. The glycoproteinatlas.org website was designed by M.S. and S.C. The study was supervised by T.W.-C. and C.R.B. The manuscript was edited by M.S., T.W.-C., and C.R.B. with input from all authors. All authors read and approved the final manuscript.

## Resource availability

All datasets and code will be made publicly available at time of final publication.

---

**Figure S1.**
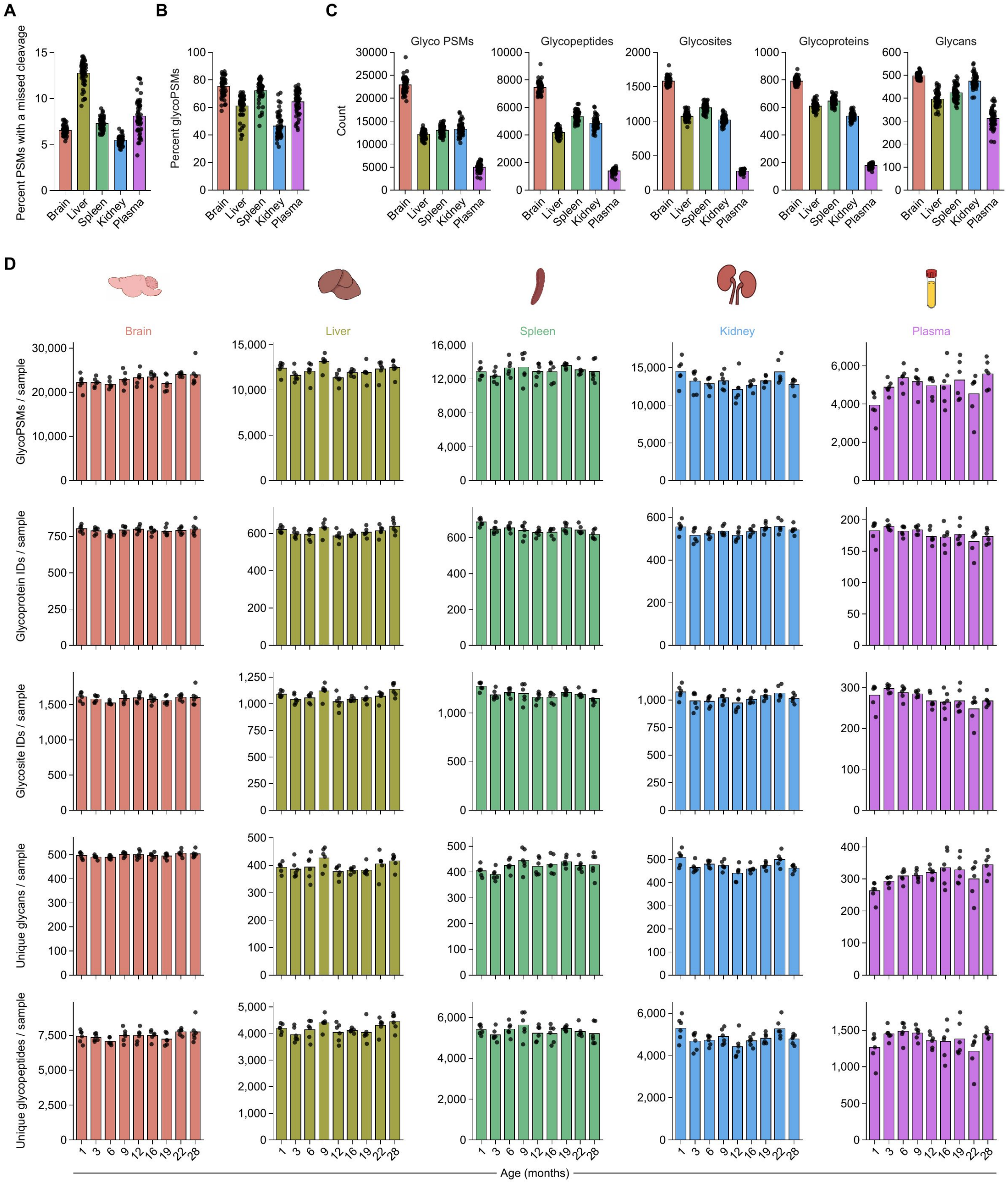
Atlas glycoproteomics sampling and quality control statistics. **(A)** Percentage of peptide-spectrum matches (PSMs) containing at least one missed tryptic cleavage in each atlas organ. **(B)** Percentage of all filtered glycoPSMs relative to all PSMs. **(C)** Numbers of glycoPSMs, unique glycopeptides, glycosites, glycoproteins, and exact glycans detected per sample. **(D)** GlycoPSMs, glycoproteins, glycosites, exact glycans, and glycopeptides detected per sample across the nine sampled ages in brain, spleen, liver, kidney, and plasma. Points represent biological samples; bars and lines denote the mean and error bars denote SEM. A glycosite is an accession-specific asparagine position, and an exact glycan is defined by monosaccharide composition.

**Figure S2.**
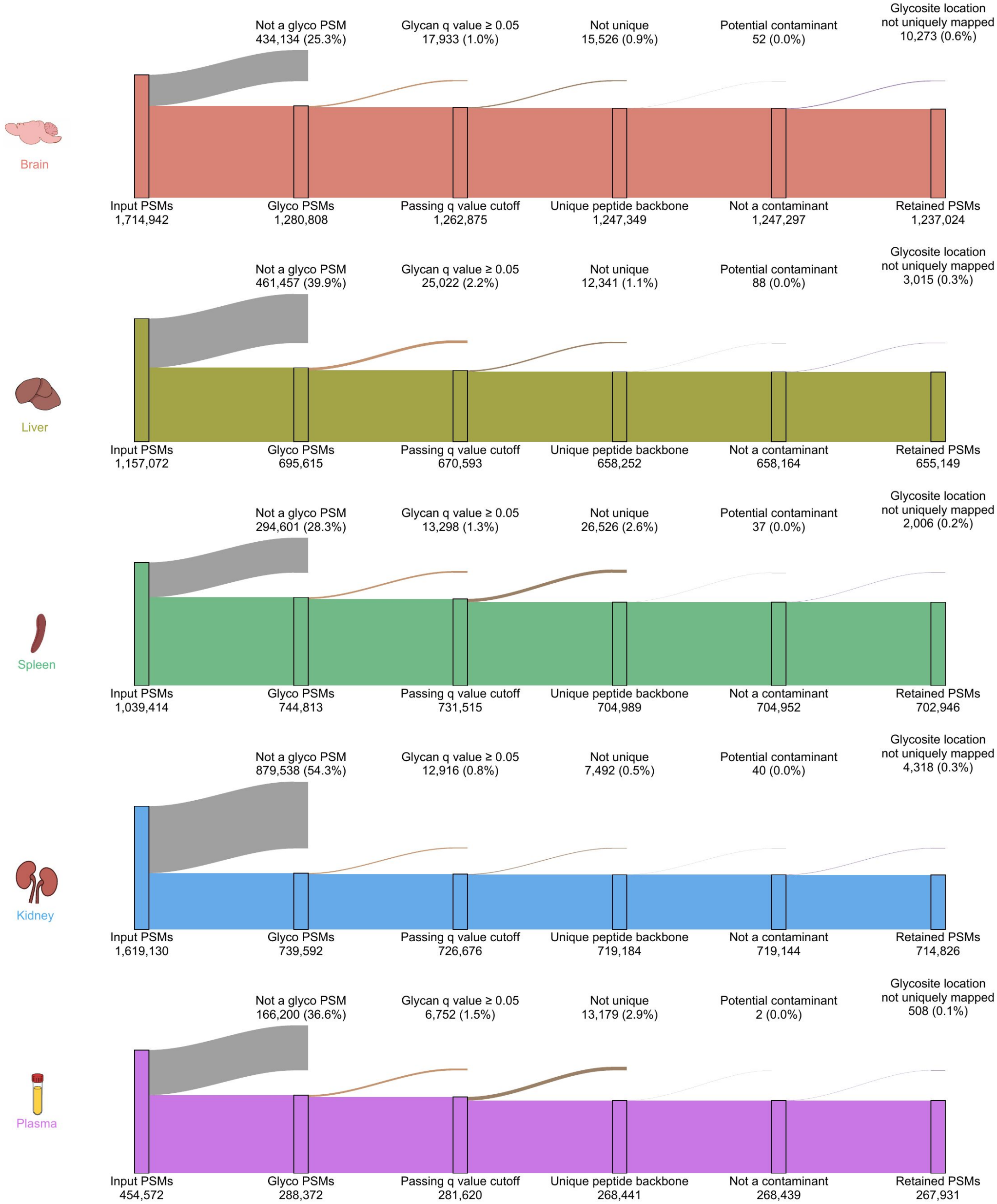
Sequential filtering of atlas PSMs. Sankey diagrams show the sequential filtering of glycoPSMs for brain, spleen, liver, kidney, and plasma. Starting PSMs were restricted to glycan-bearing spectra, glycan assignments with q < 0.05, unique peptide backbones, non-contaminant proteins, and glycopeptides that mapped unambiguously to one accession-specific glycosite. Values on the retained and excluded branches give the number and percentage of PSMs at each step.

**Figure S3.**
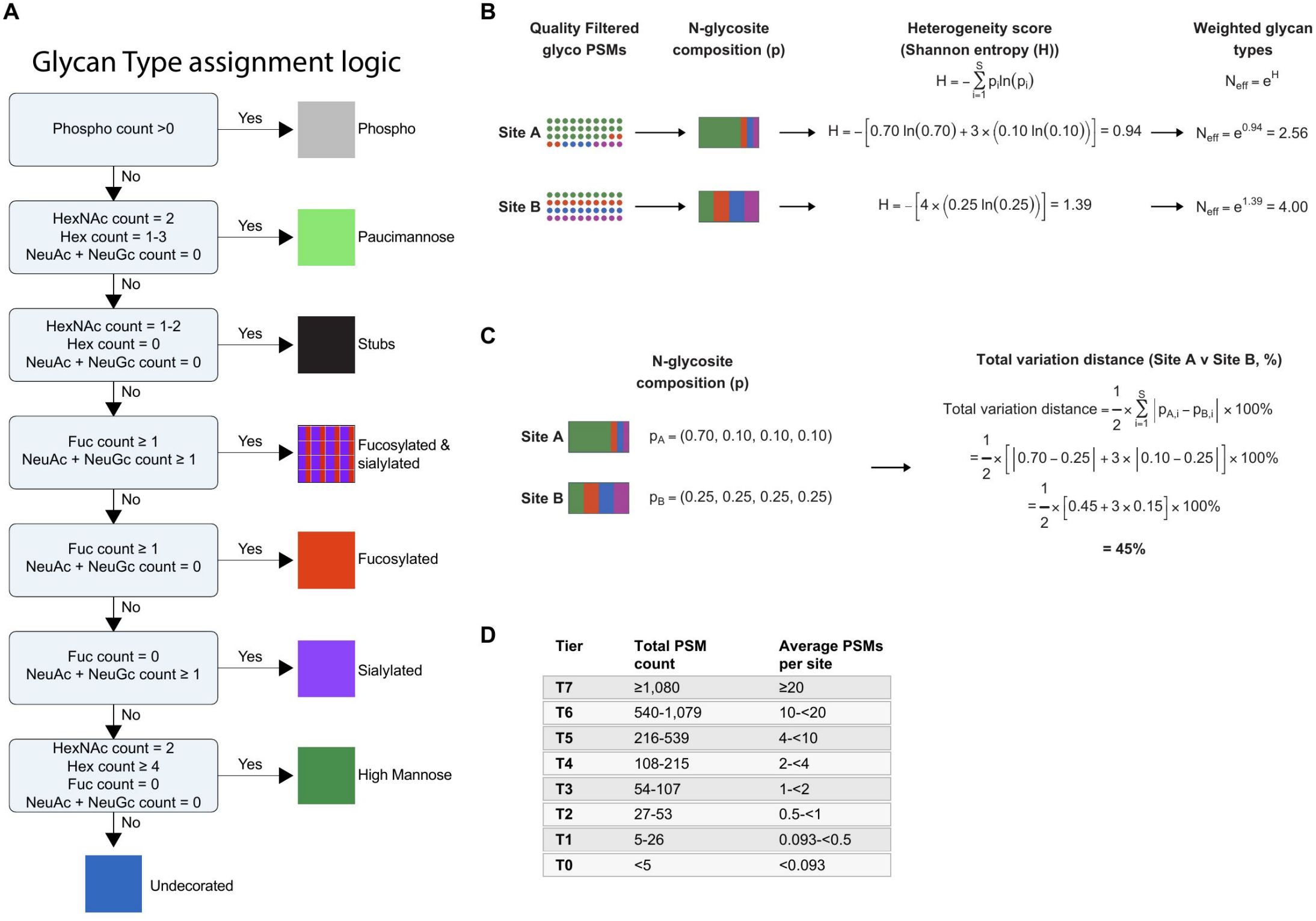
Definitions of glycan types, compositional heterogeneity, distance, and coverage tiers. **(A)** Decision tree used to assign exact N-glycan compositions to eight glycan types from phospho, HexNAc, hexose, fucose, Neu5Ac, and Neu5Gc counts. **(B)** Example calculation of heterogeneity score (Shannon entropy, H). H = −Σp_i_ln(p_i_). Weighted glycan types, exp(H), were calculated from within-site glycan type proportions. **(C)** Example calculation of total variation distance (TVD): 100 × 1/2 Σ|p_i_ − q_i_|. **(D)** Coverage tiers used for atlas glycosites. Tiers 0–7 correspond to < 5, 5–26, 27–53, 54–107, 108–215, 216–539, 540–1,079, and ≥ 1,080 total glycoPSMs, respectively; the equivalent mean depths are < 0.093, 0.093–< 0.5, 0.5–< 1, 1–< 2, 2–< 4, 4–< 10, 10–< 20, and ≥ 20 glycoPSMs per sample.

**Figure S4.**
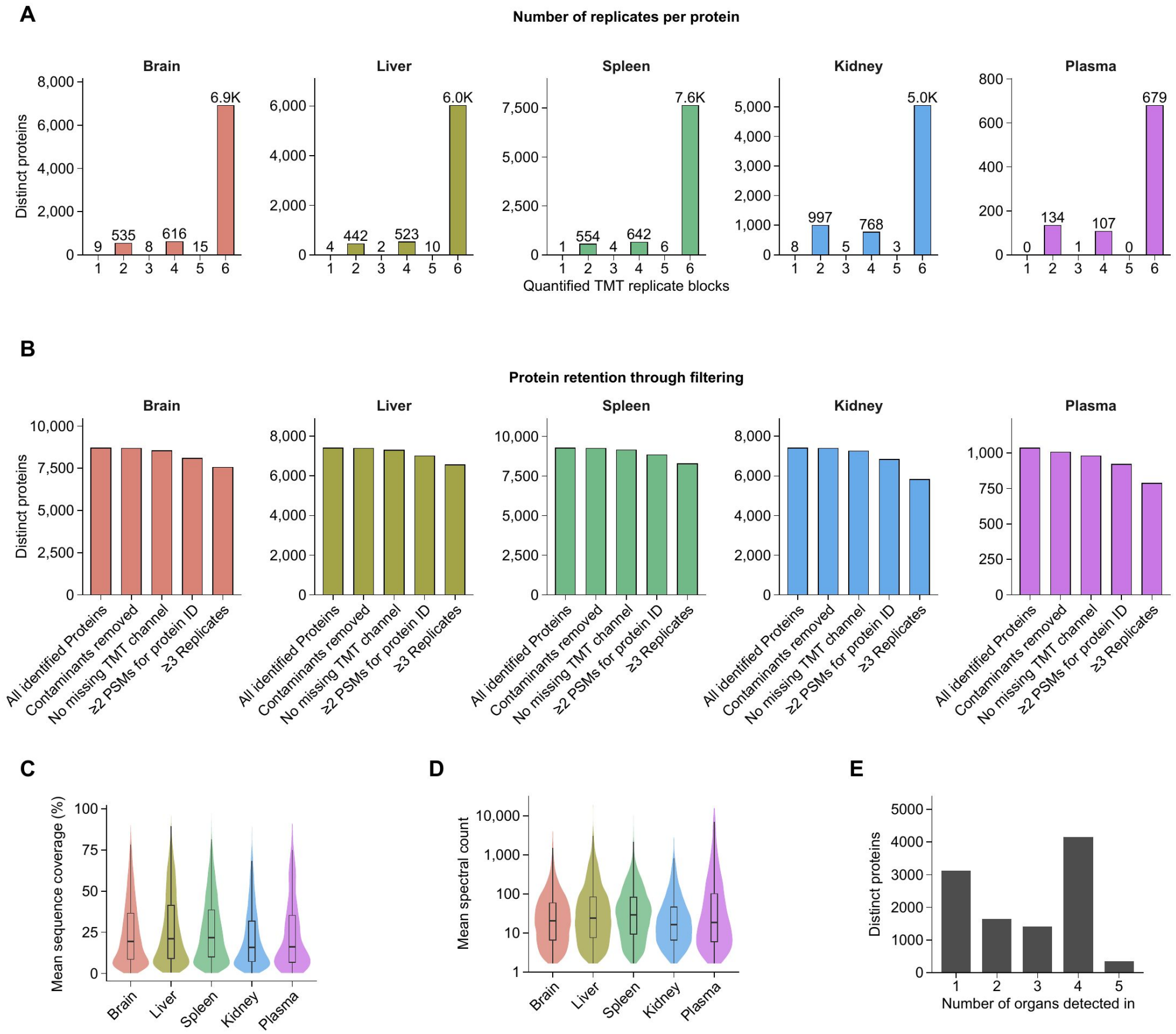
Atlas proteomics filtering and quality control. **(A)** Number of quantified tandem mass tag replicate blocks per retained protein in each organ. **(B)** Sequential protein filtering from all identified proteins through contaminant removal, complete TMT channel quantification, a minimum of two identifying PSMs, and detection in at least three biological replicates. **(C and D)** Violin plots show distributions of mean protein sequence coverage **(C)** and mean spectral count **(D)** among retained proteins; embedded box plots show the median, interquartile range, and whiskers extending to 1.5 × the interquartile range. **(E)** Number of atlas organs in which each retained protein was quantified.

**Figure S5.**
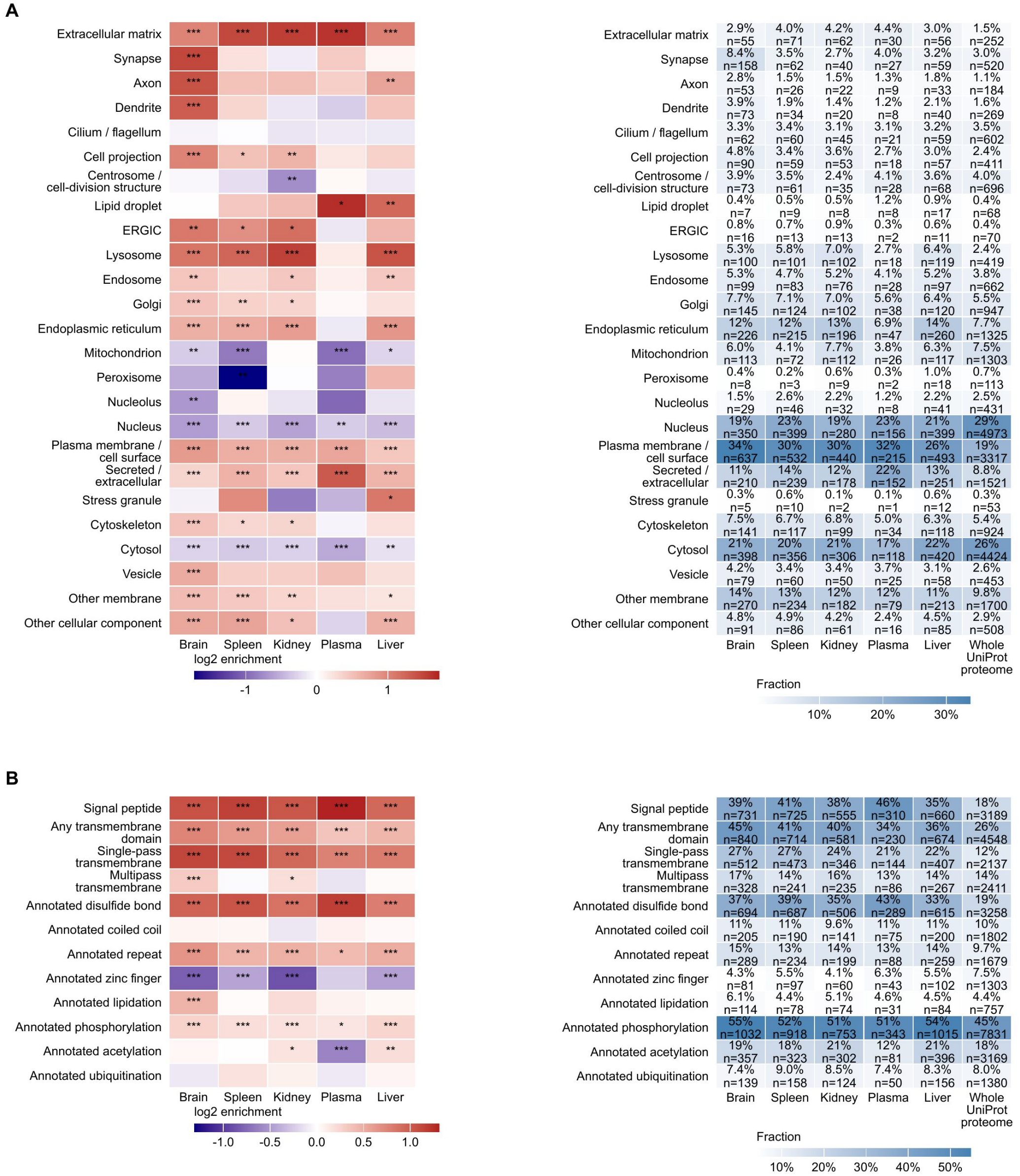
Cellular localization and protein features of the glycoproteome. **(A)** Enrichment or depletion of annotated cellular component and subcellular localization terms among glycoproteins detected in each organ and in the combined atlas, relative to the reviewed mouse proteome. The right matrix gives the fraction and number of proteins carrying each annotation. **(B)** Corresponding analysis of selected UniProt protein features, including signal peptides, transmembrane regions, disulfide bonds, repeats, zinc fingers, and annotated modifications. Color denotes log_2_ enrichment relative to the reference proteome. Two-sided Fisher’s exact tests were performed for each feature; displayed significance symbols use BH adjusted p values (*p < 0.05, **p < 0.01, ***p < 0.001, and ****p < 0.0001).

**Figure S6.**
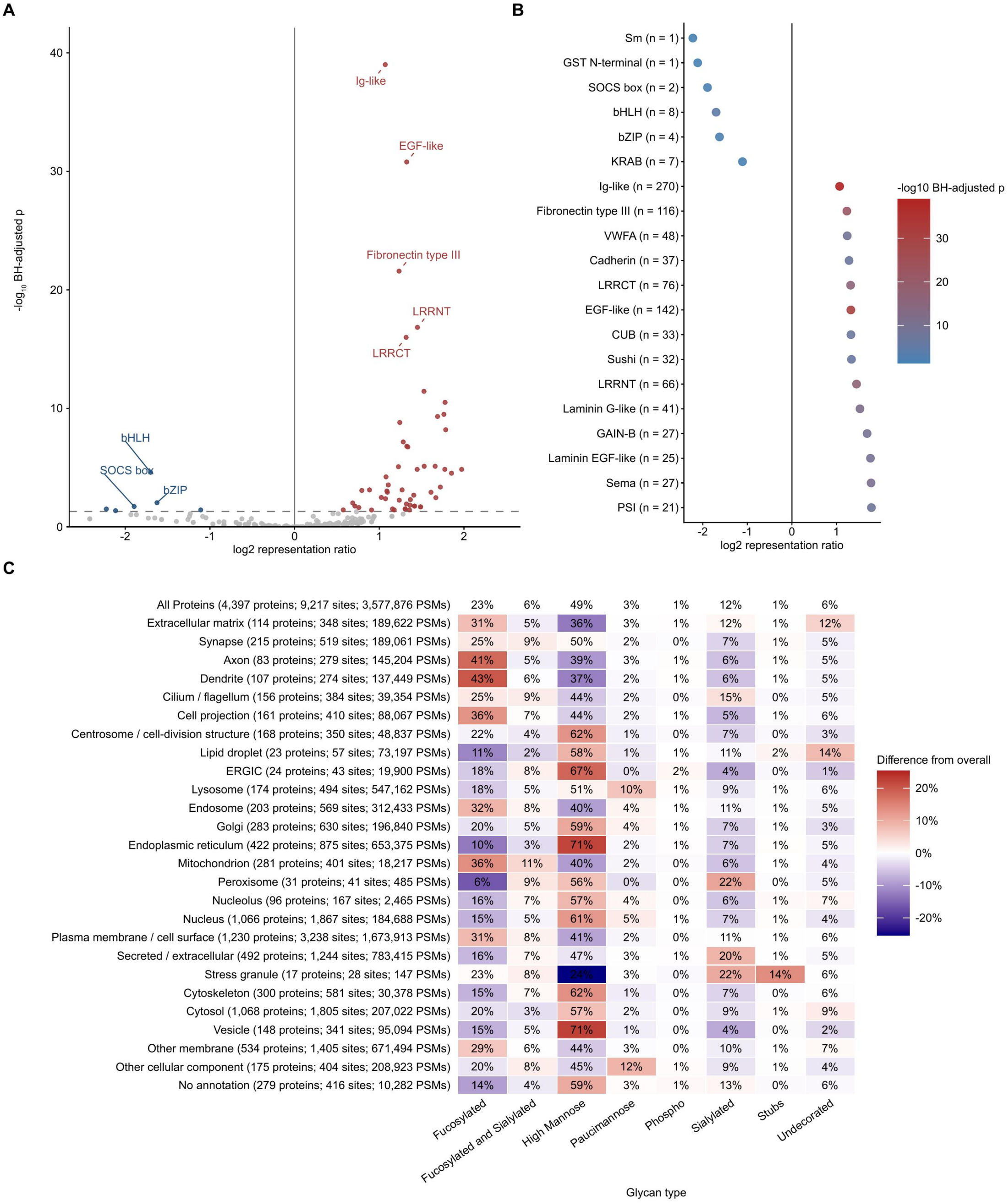
Protein domains and compartment-specific glycan composition in the atlas. **(A)** Representation of protein domains among atlas glycoproteins relative to the reference proteome. The x axis shows the glycoprotein-to-background representation ratio and the y axis shows −log_10_(BH adjusted p value); selected enriched and depleted domains are labeled. **(B)** Representation ratios for selected domains. Numbers give detected glycoproteins carrying the domain; point color denotes −log_10_(BH adjusted p value). **(C)** PSM-weighted glycan type composition of glycoproteins assigned to the indicated subcellular localizations. Cell values are percentages, and color shows the percentage-point difference from the overall atlas composition. Row labels report the numbers of proteins, glycosites, and glycoPSMs contributing to each localization. Domain and localization enrichment used two-sided Fisher’s exact tests with BH adjustment.

**Figure S7.**
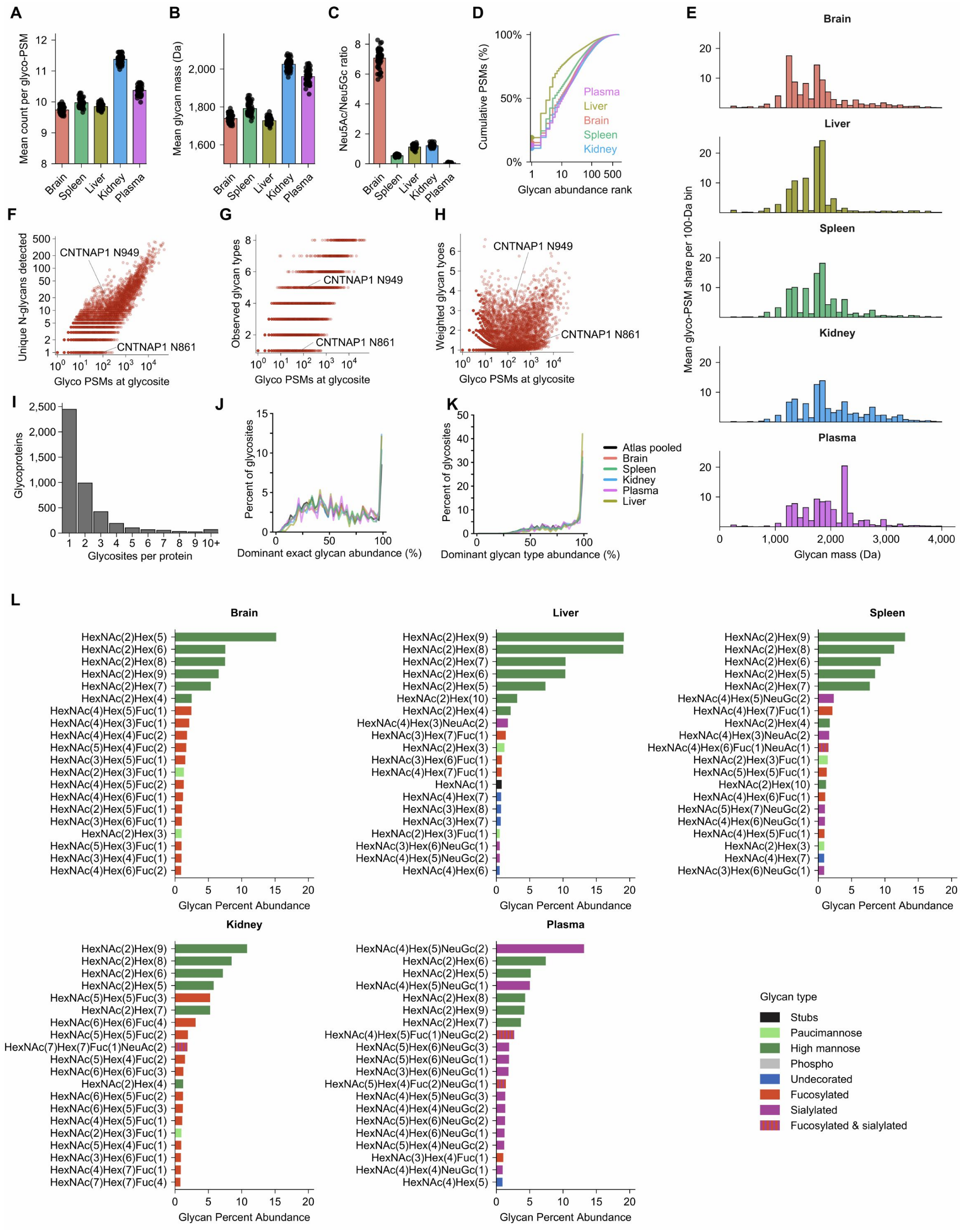
Glycan abundance and heterogeneity across organs. **(A)** Mean number of monosaccharides per glycoPSM in each organ. **(B)** PSM-weighted mean glycan mass by organ. **(C)** Neu5Ac/Neu5Gc monosaccharide ratio by organ. **(D)** Cumulative glycoPSM abundance of exact N-glycans ranked within each organ. **(E)** Mean glycoPSM share in 100-Da glycan mass bins. **(F–H)** Relationships between glycoPSM coverage and unique exact N-glycans **(F)**, observed glycan types **(G)**, or weighted glycan types **(H)**, at each atlas-pooled glycosite. CNTNAP1 N861 and N949 illustrate distinct heterogeneity at comparable coverage. **(I)** Number of glycosites detected per glycoprotein. **(J and K)** Distributions of dominant glycan abundance **(J)** and dominant glycan type abundance **(K)** among Tier 1+ glycosites for the pooled atlas and individual organs. At each glycosite, the percent abundance of the most abundant exact glycan or glycan type is used. **(L)** The 20 exact N-glycan compositions contributing the largest fraction of pooled glycoPSMs are shown separately for brain, spleen, liver, kidney, and plasma. Percent abundance was calculated from all glycoPSMs pooled within an organ. Monosaccharide counts exclude phosphate. Points in **(A–C)** are biological samples; bars denote means and error bars denote SEM.

**Figure S8.**
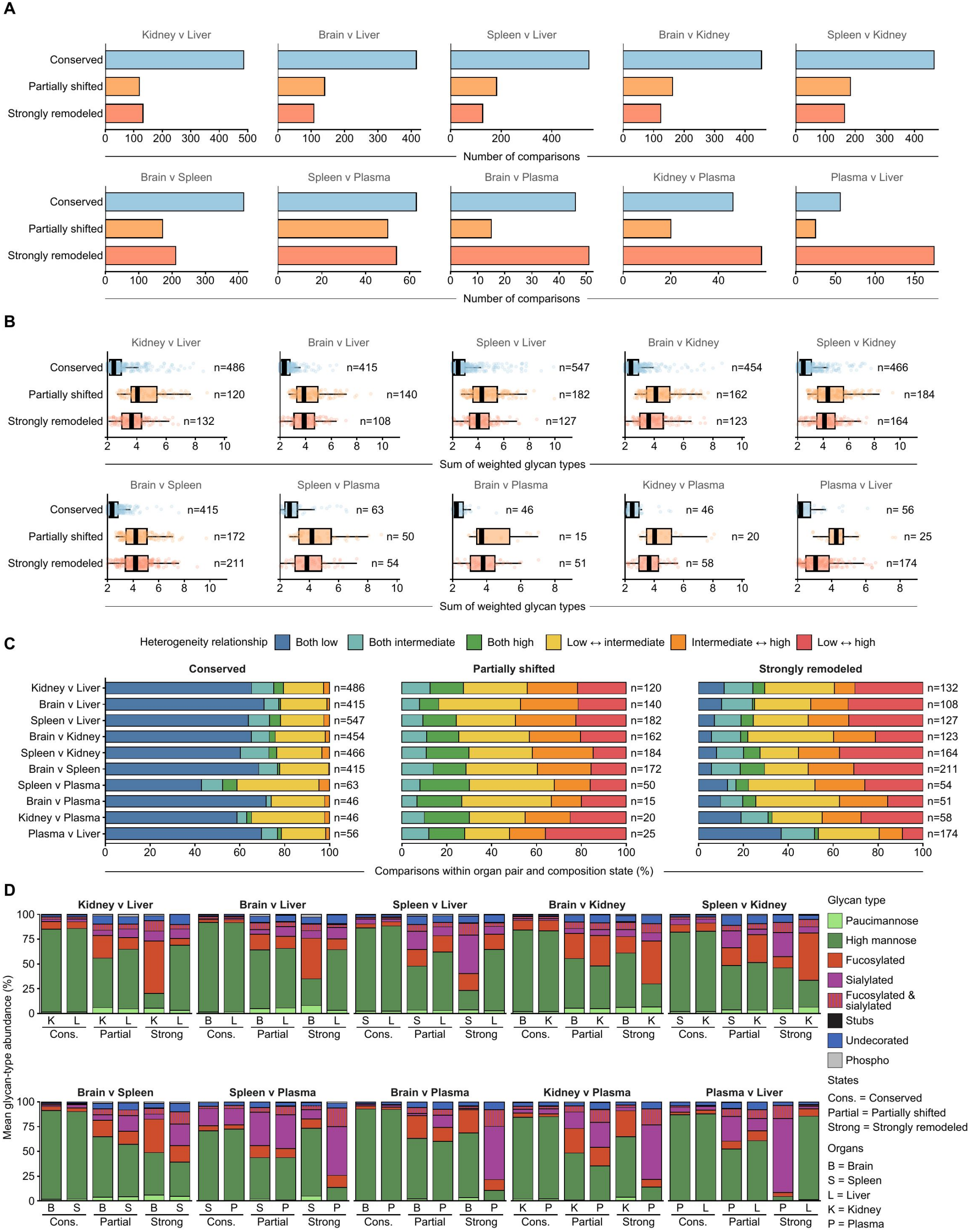
Glycan type heterogeneity and composition across inter-organ compositional states. **(A)** Numbers of Tier 1+ glycosites classified as conserved (TVD < 20%), partially shifted (20% ≤ TVD < 50%), or strongly remodeled (TVD ≥ 50%) for each organ pair. **(B)** Sum of weighted glycan types across the two organs for every site comparison in each compositional state. The weighted glycan types metric was calculated as exp(Shannon H) from pooled site-level PSM proportions in each organ. Numbers denote site comparisons; box plots show the median, interquartile range, and whiskers extending to 1.5 × the interquartile range. **(C)** Percentage of comparisons in each state with low, intermediate, or high heterogeneity in the two organs. Low heterogeneity was defined as < 1.5 weighted glycan types, high heterogeneity as ≥ 2.5, and intermediate heterogeneity as values between these cutoffs; mixed categories indicate different heterogeneity classes in the two organs. **(D)** Mean glycan type composition is shown for the two organ members of each Tier 1+ glycosite comparison, stratified as conserved (TVD < 20%), partially shifted (20% ≤ TVD < 50%), or strongly remodeled (TVD ≥ 50%). Each glycosite contributes one composition calculated from pooled glycoPSMs per organ; stacked bars are the site-weighted means across eligible glycosites and sum to 100%. Numbers above the pairs denote shared glycosites. B, brain; S, spleen; L, liver; K, kidney; P, plasma.

**Figure S9.**
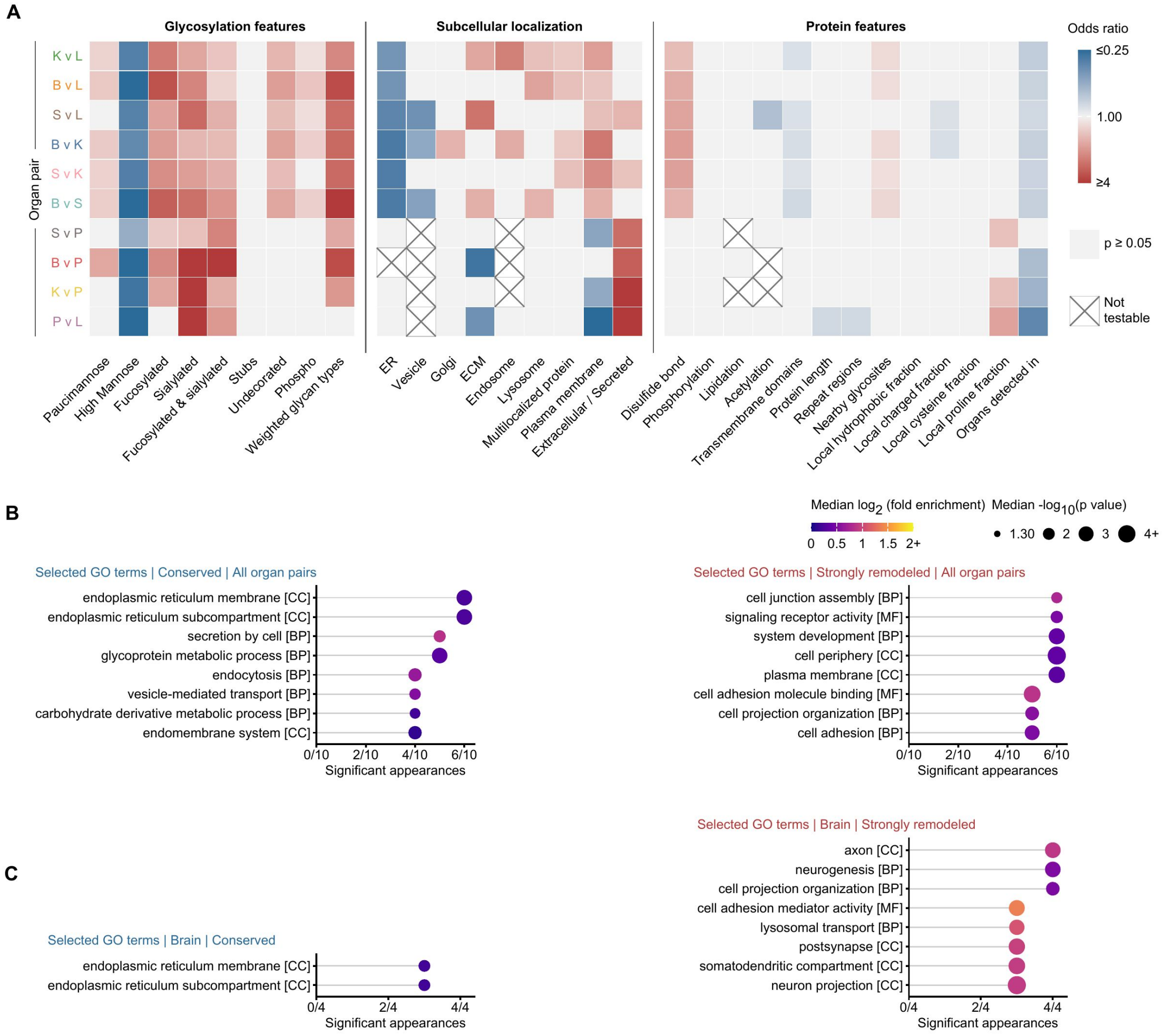
Features and GO programs associated with conserved and strongly remodeled glycosites. **(A)** Odds ratios from separate unadjusted proportional-odds models relating glycosylation, subcellular localization, and protein features to the ordered remodeling state for each organ pair. Red and blue denote increased and decreased odds of greater remodeling, respectively; colored cells have p < 0.05, gray cells have p ≥ 0.05, and crossed cells were not testable. **(B and C)** Selected Gene Ontology (GO) terms recurring among proteins carrying conserved or strongly remodeled Tier 1+ glycosites are shown across all ten organ pairs **(B)** and comparisons containing brain **(C)**. For each organ pair and compositional state, the foreground was the unique proteins carrying qualifying sites and the background was the unique proteins carrying any tested Tier 1+ site in that organ pair. The x axis gives the number of pairwise enrichment profiles in which a term had p < 0.05; point color denotes the median log_2_ fold enrichment and point size denotes the median −log_10_(p value) across those appearances. Enrichment used one-sided hypergeometric tests. BP, biological process; CC, cellular component; MF, molecular function.

**Figure S10.**
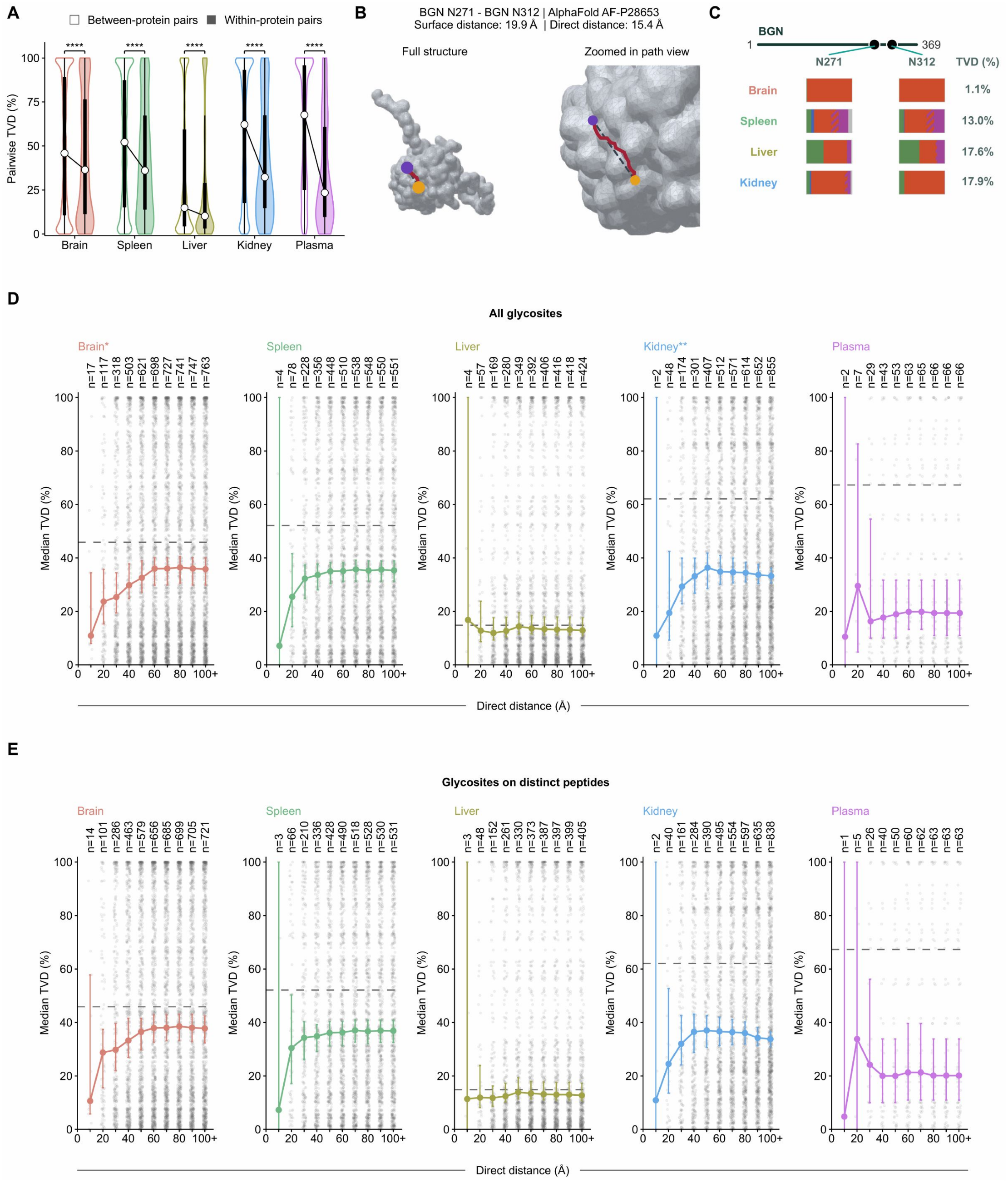
Within-protein and distance-based similarity of glycosite compositions. **(A)** Pairwise TVD for Tier 2+ glycosite pairs drawn from different proteins or from the same protein in each organ. White circles mark medians. Two-sided Wilcoxon rank-sum tests compare the pair-level distributions; ****p < 0.0001. **(B)** AlphaFold model of BGN showing N271 and N312 and the shortest surface path between them; surface and direct distances are reported above the model. **(C)** Pooled glycan type compositions of BGN N271 and N312 and their within-organ TVDs. **(D and E)** Median TVD among Tier 2+ within-protein site pairs as progressively larger direct-distance thresholds are included, using all site pairs **(D)** or only sites observed on distinct peptides **(E)**. Gray points are pair-level TVDs; colored points and lines are medians, error bars are exact nonparametric 95% confidence intervals for the median, dashed lines are the median of all eligible pairs, and n above each threshold is the number of included pairs. The 100+ point includes all measured pairs. Asterisks appended to organ labels denote p values from two-sided Wilcoxon rank-sum comparisons of pairs at ≤ 20 Å versus > 20 Å (*p < 0.05; **p < 0.01).

**Figure S11.**
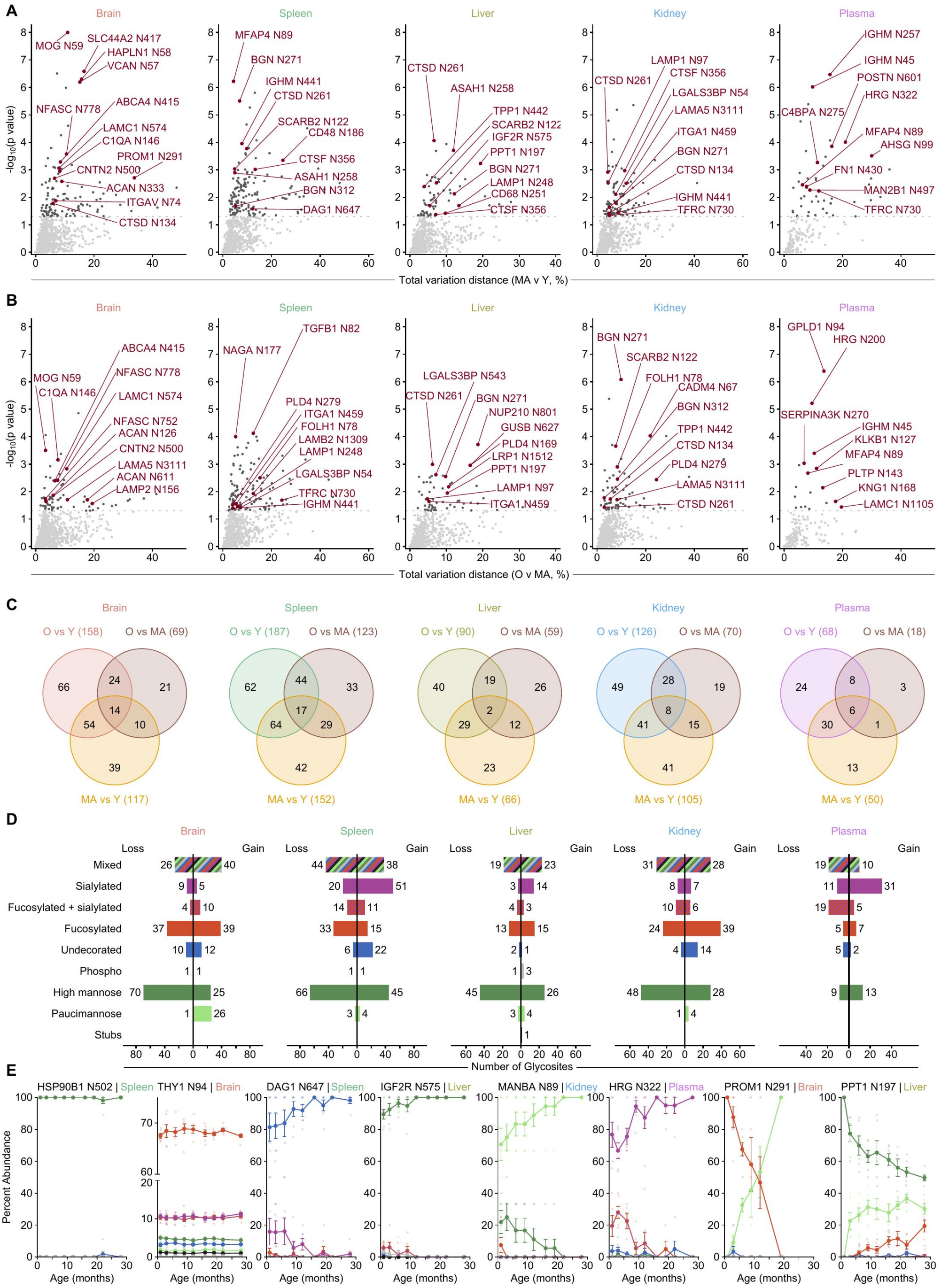
Additional age-bin comparisons, glycan type transitions, and representative trajectories. **(A and B)** Glycosite compositional change for middle-aged versus young animals **(A)** and old versus middle-aged animals **(B)** in each organ. The horizontal dashed line marks raw p = 0.05, and selected sites are labeled. **(C)** Overlap of glycosites with p < 0.05 in old versus young, old versus middle-aged, and middle-aged versus young comparisons. Venn areas are schematic and numbers are exact. **(D)** Dominant glycan types lost and gained among old-versus-young hits. A glycan type is dominant if it accounts for at least 60% of summed gains or absolute losses; otherwise, the direction is classified as mixed. Young, middle-aged, and old atlas bins were 1–6, 9–16, and 19–28 months, respectively. Each tested site required detection in at least nine animals and at least 18 glycoPSMs in each age bin. P values were obtained by fixed-size animal-label permutation; displayed hits use p < 0.05. **(E)** Age trajectories of glycan type percent abundance at representative glycosites. Examples include stable compositions (HSP90B1 N502, THY1 N94), broadly shared losses of high mannose, and organ-specific gains of paucimannose or sialylated glycans. Faint points represent individual animals; symbols and error bars show mean ± SEM.

**Figure S12.**
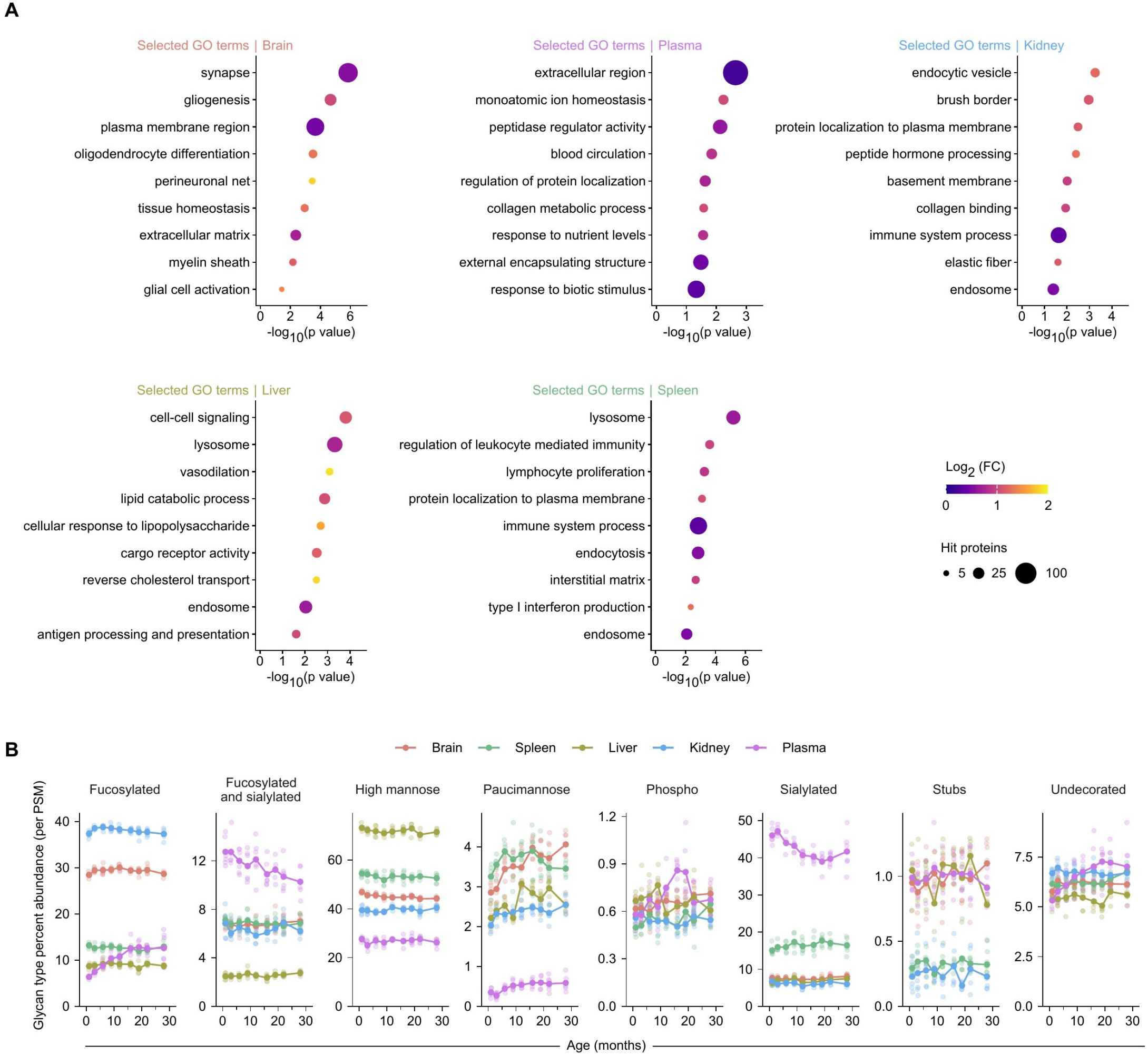
Organ-specific GO enrichment and global glycan type composition across aging. **(A)** Selected GO terms enriched among proteins carrying old-versus-young glycosite-composition hits are shown separately for each atlas organ. Within an organ, the foreground comprised unique proteins carrying hit sites and the background comprised unique proteins carrying sites eligible for the old-versus-young test. Enrichment used one-sided hypergeometric tests. BP, biological process; CC, cellular component; MF, molecular function. **(B)** Glycan type abundance across age. Each point represents a biological sample; lines denote means and error bars denote SEM.

**Figure S13.**
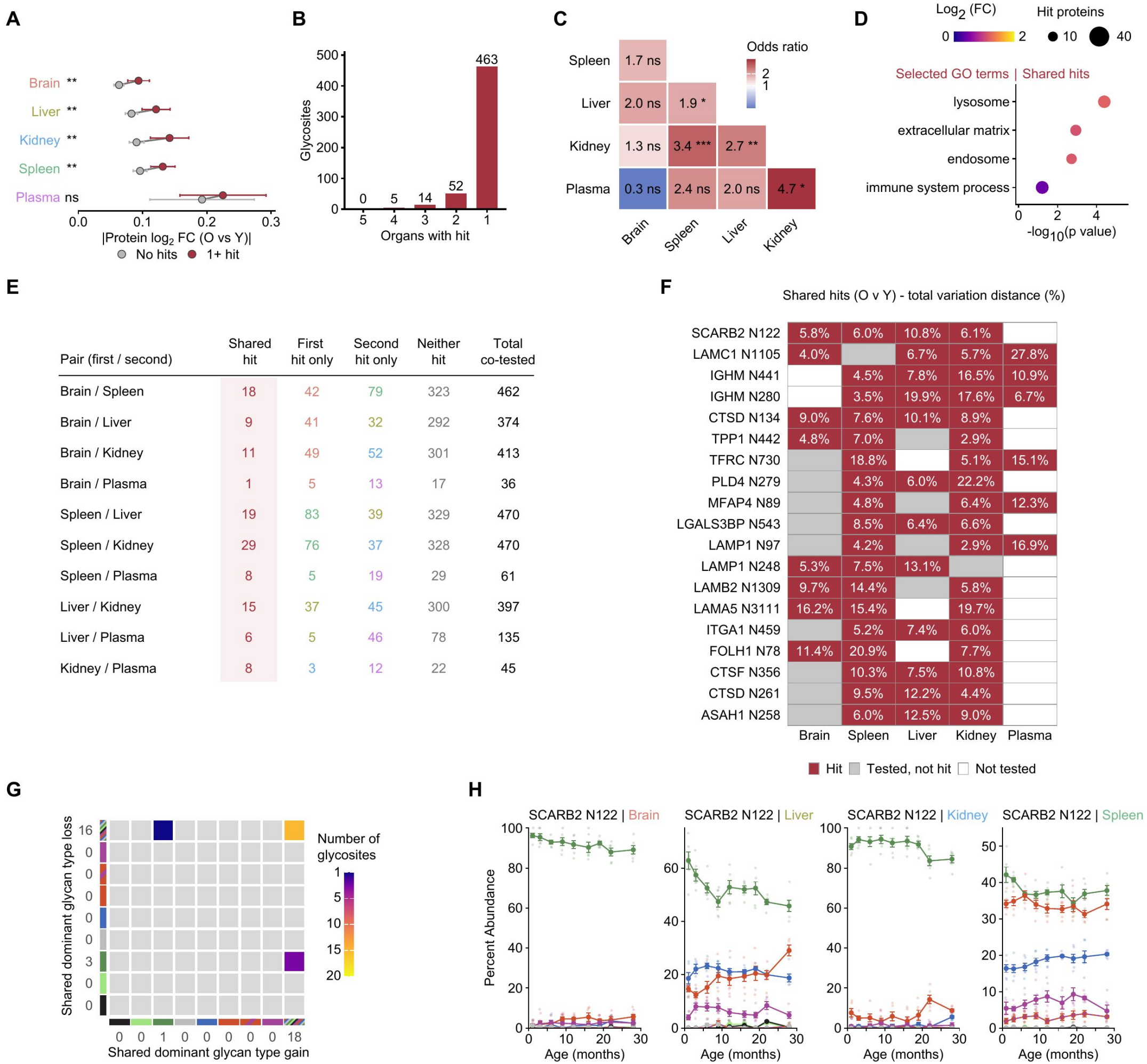
Protein abundance and cross-organ recurrence of age-remodeled glycosites. **(A)** Absolute old-versus-young protein log_2_ fold change for proteins with no glycosite composition hit or at least one hit, shown separately for each organ. Points denote proteins and horizontal intervals show mean ± SEM. Two-sided Welch t tests compare groups; symbols report p values. **(B)** Number of old-versus-young hit glycosites detected as hits in one to five organs. **(C)** Pairwise association of hit status among sites testable in both organs. Cells report odds ratios from Fisher’s exact tests and p value symbols. For **(A)** and **(C)**, significance symbols denote *p < 0.05, **p < 0.01, ***p < 0.001, and ****p < 0.0001. **(D)** Selected GO terms enriched among proteins carrying the same hit site in at least two organs. The background was proteins carrying a site testable in at least two organs; one-sided hypergeometric p values are plotted. **(E)** Contingency counts underlying panel C. **(F)** Old-versus-young TV for selected sites significant in multiple organs. Red cells are hits, gray cells were tested but not significant, and white cells were not tested. **(G)** Concordant dominant glycan type losses and gains among shared hits; numbers are glycosites. A glycan type is dominant if it accounts for at least 60% of summed gains or absolute losses. **(H)** Glycan type trajectories for SCARB2 N122. High mannose content declines across tissues but to different degrees and at different ages. Points are animals; lines show mean ± SEM.

**Figure S14.**
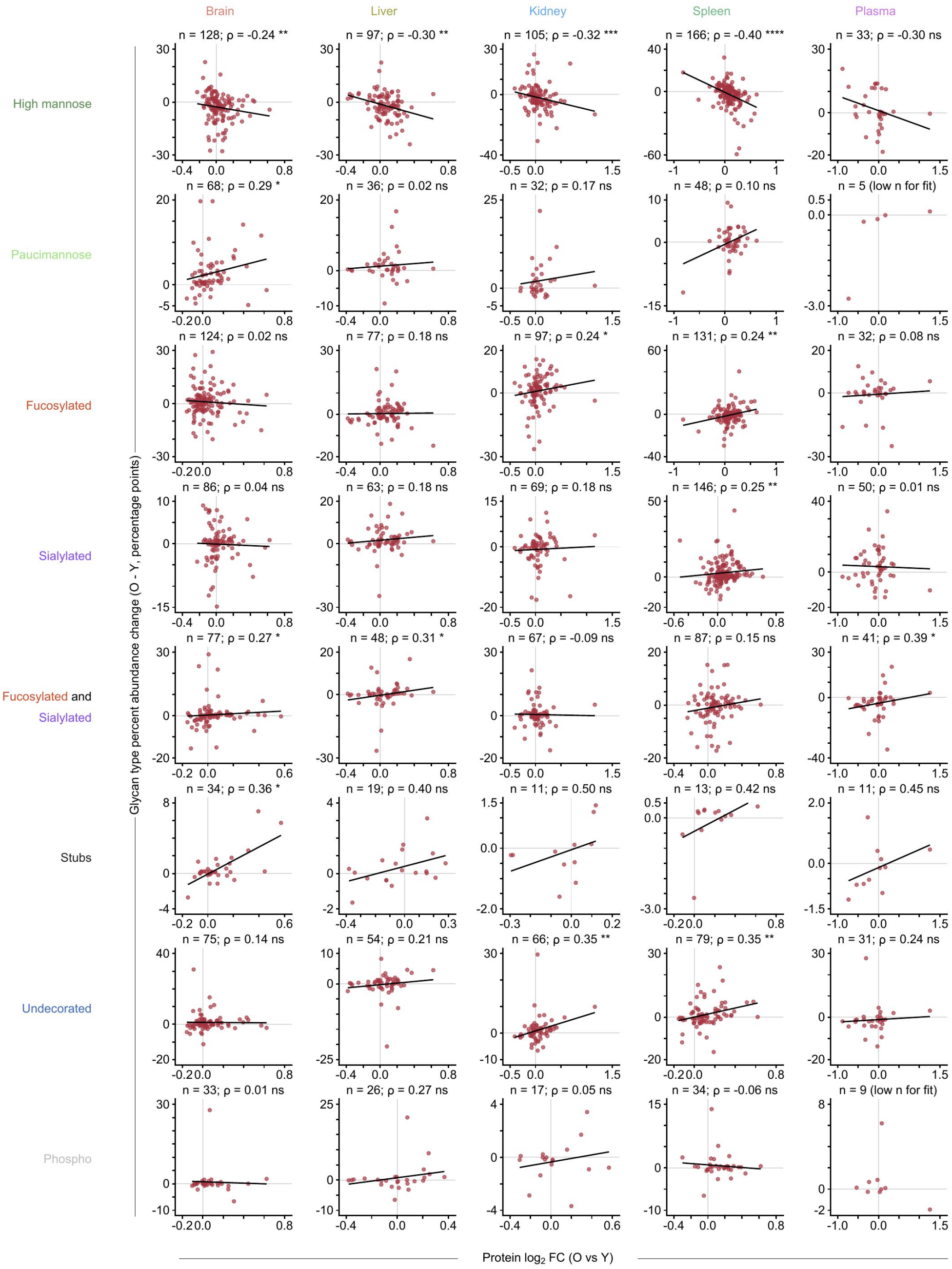
Relationship between protein abundance and glycan type remodeling with age. For each organ and glycan type, old-versus-young protein log_2_ fold change is plotted against the mean old-versus-young percentage-point change in that glycan type across included glycosites on the protein. Glycosites were included if they were a hit in any age comparison (O v Y, O v MA, or MA v Y). One point represents one protein; when a protein carried multiple included glycosites, site-level changes were averaged. Panel annotations give the number of proteins, Spearman’s ρ, and its two-sided p value. Significance symbols denote *p < 0.05, **p < 0.01, ***p < 0.001, and ****p < 0.0001; ns, not significant. Black lines show linear regression fits for visualization.

**Figure S15.**
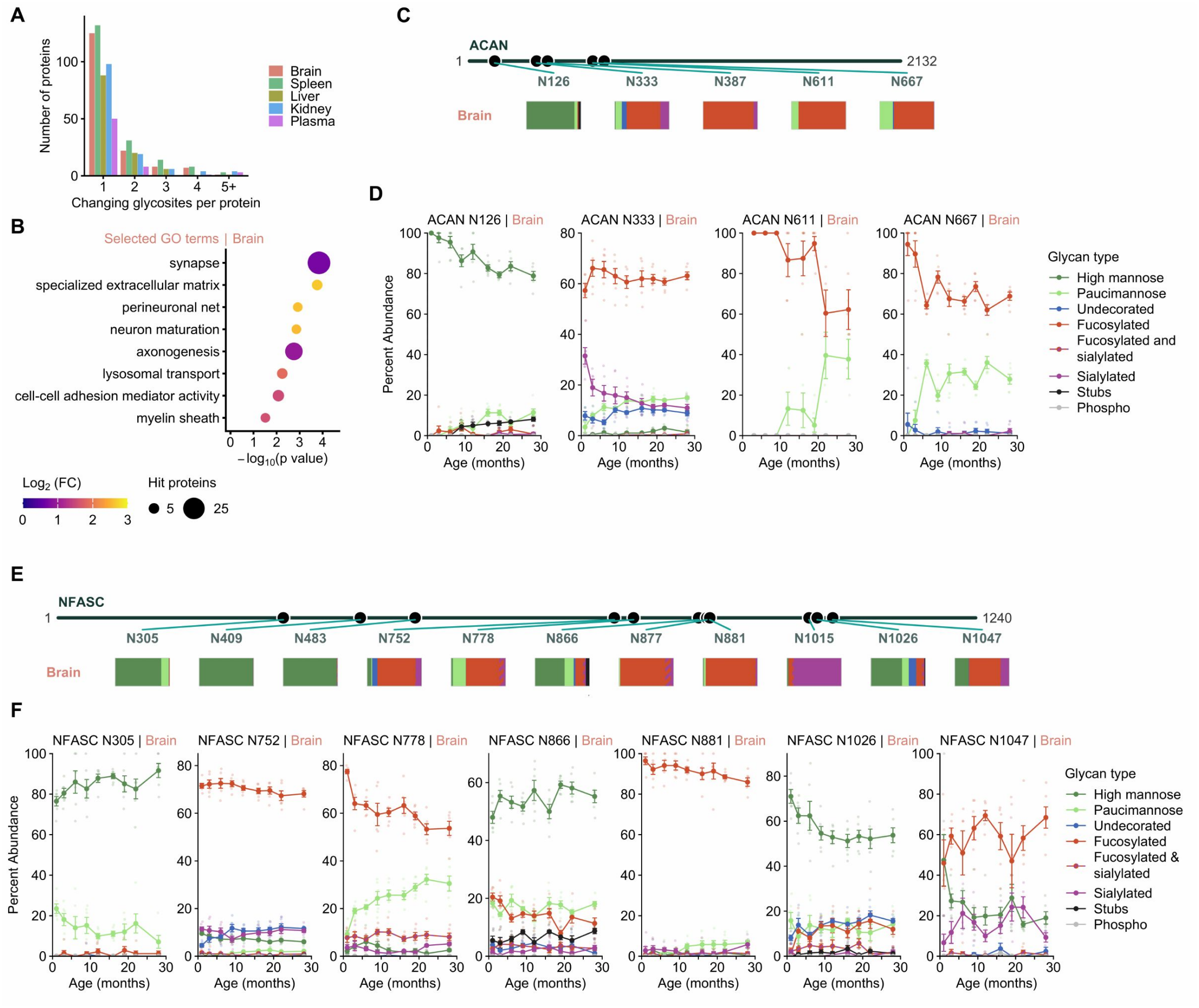
Proteins containing multiple age-remodeled glycosites. **(A)** Distribution of the number of age-remodeled glycosites per protein in each organ. **(B)** Selected GO terms enriched among brain proteins carrying at least two distinct hit glycosites. The foreground was proteins with at least two hits and the background was proteins with at least two tested sites. **(C)** Pooled glycan type composition of selected ACAN glycosites in the indicated organs. **(D)** Age trajectories for selected ACAN sites. **(E)** Pooled glycan type composition of selected NFASC glycosites. **(F)** Age trajectories for selected NFASC sites. In trajectory panels, points are animals and lines and error bars show mean ± SEM. Age-remodeled sites were defined by p < 0.05 in at least one atlas age-bin comparison.

**Figure S16.**
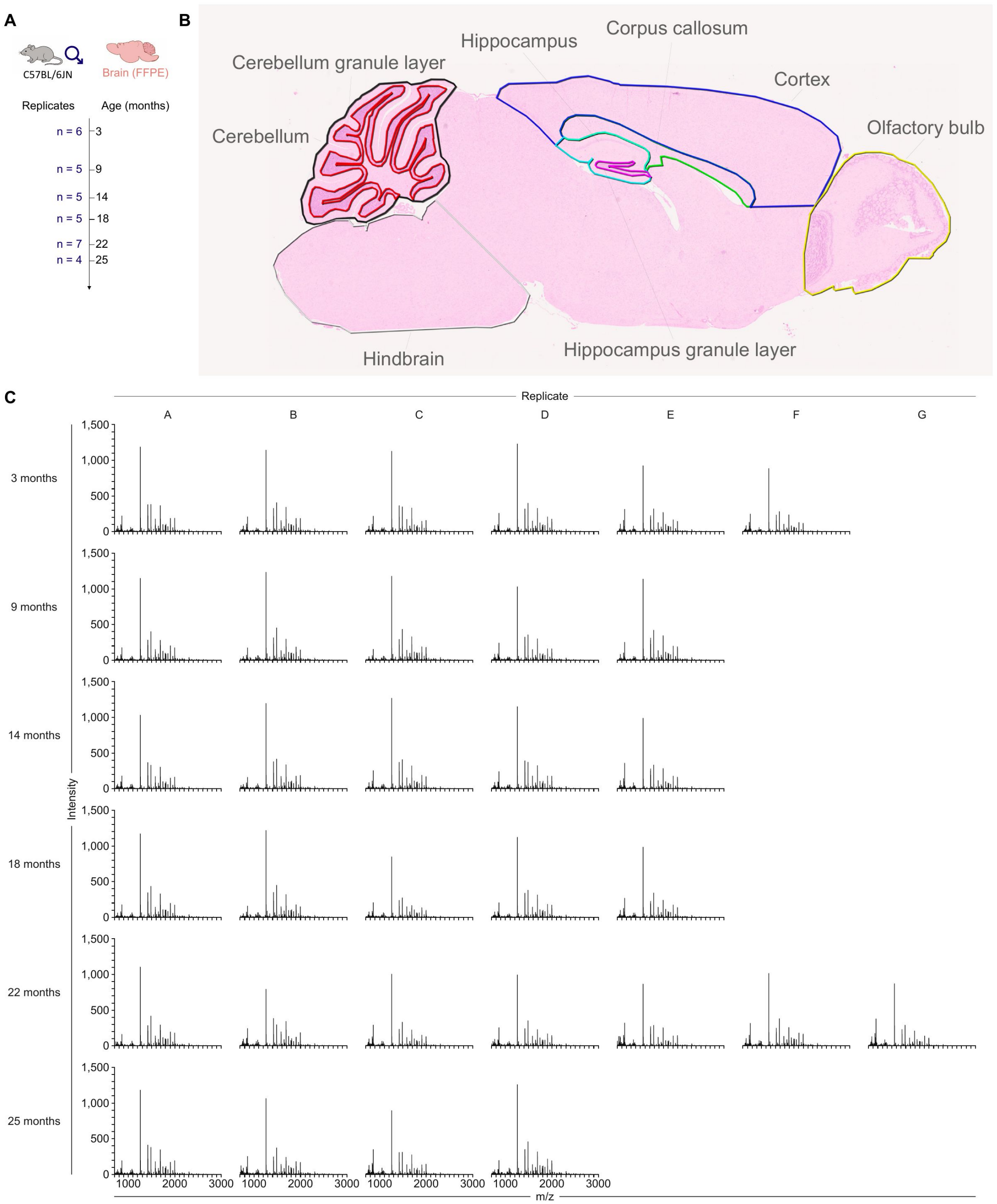
MALDI–MSI cohort, anatomical regions, and raw whole-brain spectra. **(A)** Cohort of male C57BL/6JN mice used for MALDI–MSI of formalin-fixed paraffin-embedded sagittal brain sections. Numbers of biological replicates are shown for each age. **(B)** Representative sagittal section stained with hematoxylin and eosin showing the eight regions of interest used for spatial analysis: cerebellum, cerebellar granule layer, hindbrain, hippocampus, hippocampal granule layer, corpus callosum, cortex, and olfactory bulb. **(C)** Unprocessed MALDI spectra from individual sections, arranged by age and biological replicate. Intensity is plotted against mass-to-charge ratio (m/z).

**Figure S17.**
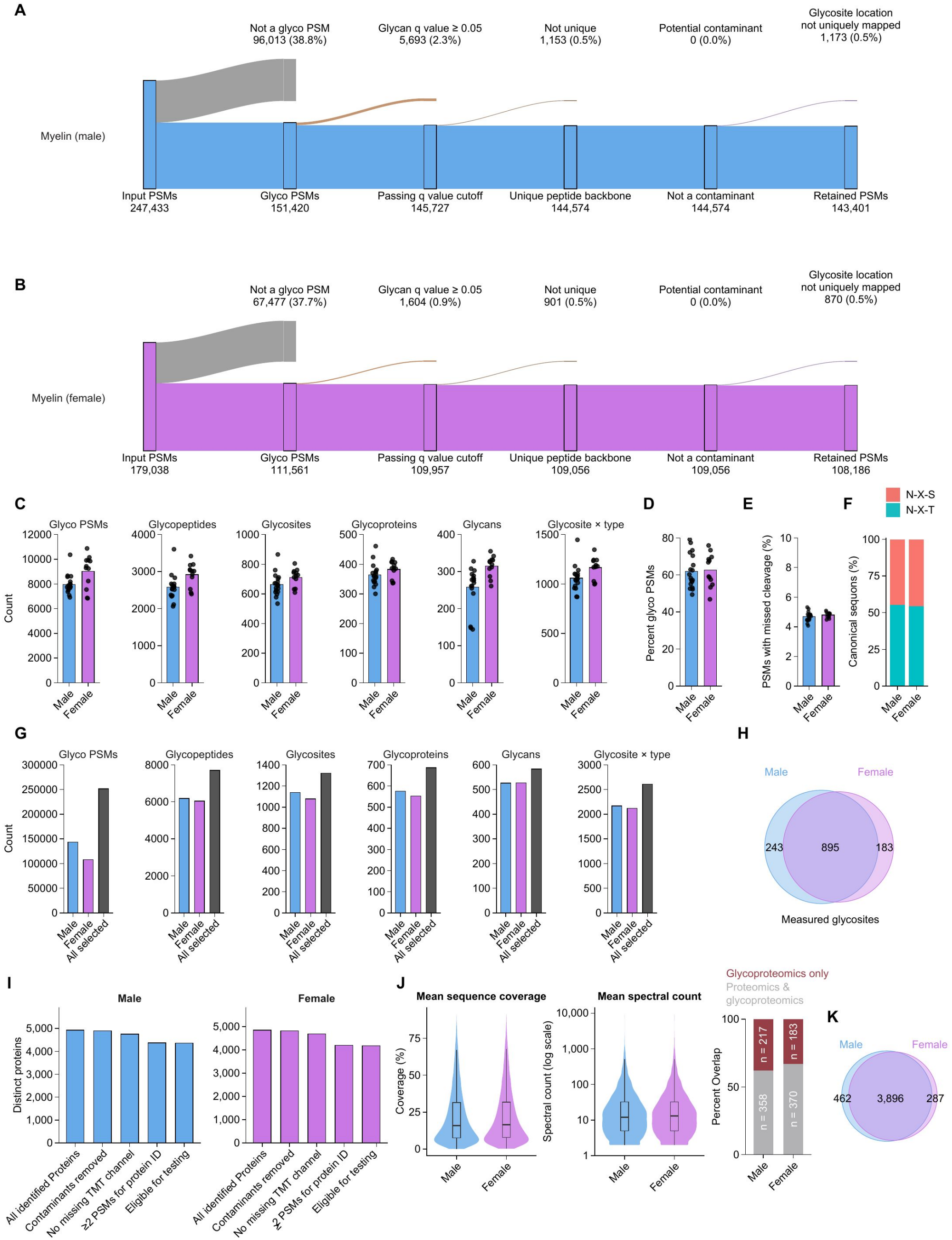
Myelin glycoproteomics and proteomics filtering, depth, and overlap. **(A and B)** Sequential glycoPSM filtering for male myelin **(A)** and female myelin **(B)**, from input spectra through glycan q value < 0.05, unique peptide backbones, contaminant removal, and unambiguous glycosite mapping. Branch labels give retained or excluded PSM counts and percentages. **(C)** Per-sample numbers of glycoPSMs, glycopeptides, glycosites, glycoproteins, exact glycans, and glycosite–glycan type combinations. **(D)** Percentage of all PSMs assigned to quality-filtered glycopeptides. **(E)** Percentage of PSMs with at least one missed cleavage. **(F)** Fraction of localized sites in N-X-S or N-X-T sequons. **(G)** Aggregate numbers of identified entities in male myelin, female myelin, and the combined dataset. **(H)** Overlap of glycosites detected in male and female myelin. Points in C–F are biological samples; bars denote means and error bars denote SEM. **(I)** Sequential filtering of male and female myelin proteomics data from identified proteins through contaminant removal, complete reporter channel quantification, minimum PSM support, and replicate coverage, following reprocessing of previously published data. **(J)** Violin plots show distributions of protein sequence coverage and mean spectral count in male and female myelin; embedded box plots show the median, interquartile range, and whiskers extending to 1.5 × the interquartile range. Stacked bars show the percentage of glycoproteins also quantified by total proteomics and the complementary percentage detected only by glycoproteomics. **(K)** Overlap of proteins quantified in male and female myelin.

**Figure S18.**
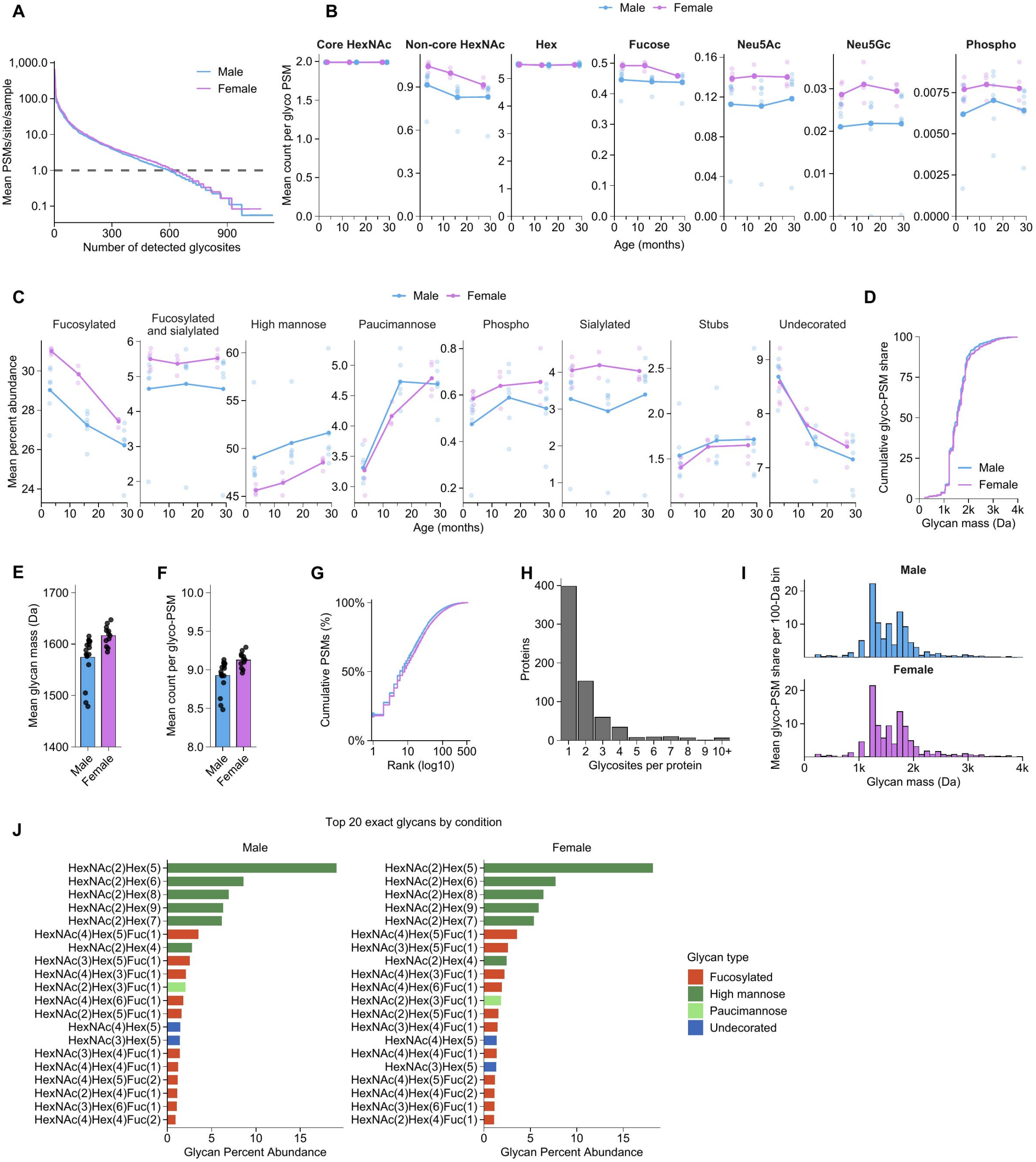
Global properties of the male and female myelin glycoproteomes. **(A)** Glycosites ranked by mean PSM depth per sample; the dashed line marks one mean PSM per sample. **(B)** Mean monosaccharide counts per glycoPSM across age. **(C)** Mean glycan type percent abundance across age. **(D)** Cumulative glycoPSM share as a function of glycan mass. **(E)** PSM-weighted mean glycan mass. **(F)** Mean total monosaccharide count per glycoPSM. **(G)** Cumulative glycoPSM abundance of exact N-glycans ranked within each myelin dataset. **(H)** Numbers of glycosites detected per glycoprotein. **(I)** Mean glycoPSM share in 100-Da glycan mass bins. **(J)** Twenty most abundant exact glycans in male and female myelin, calculated from pooled glycoPSMs. Points are biological samples; bars or lines show means and error bars show SEM where present.

**Figure S19.**
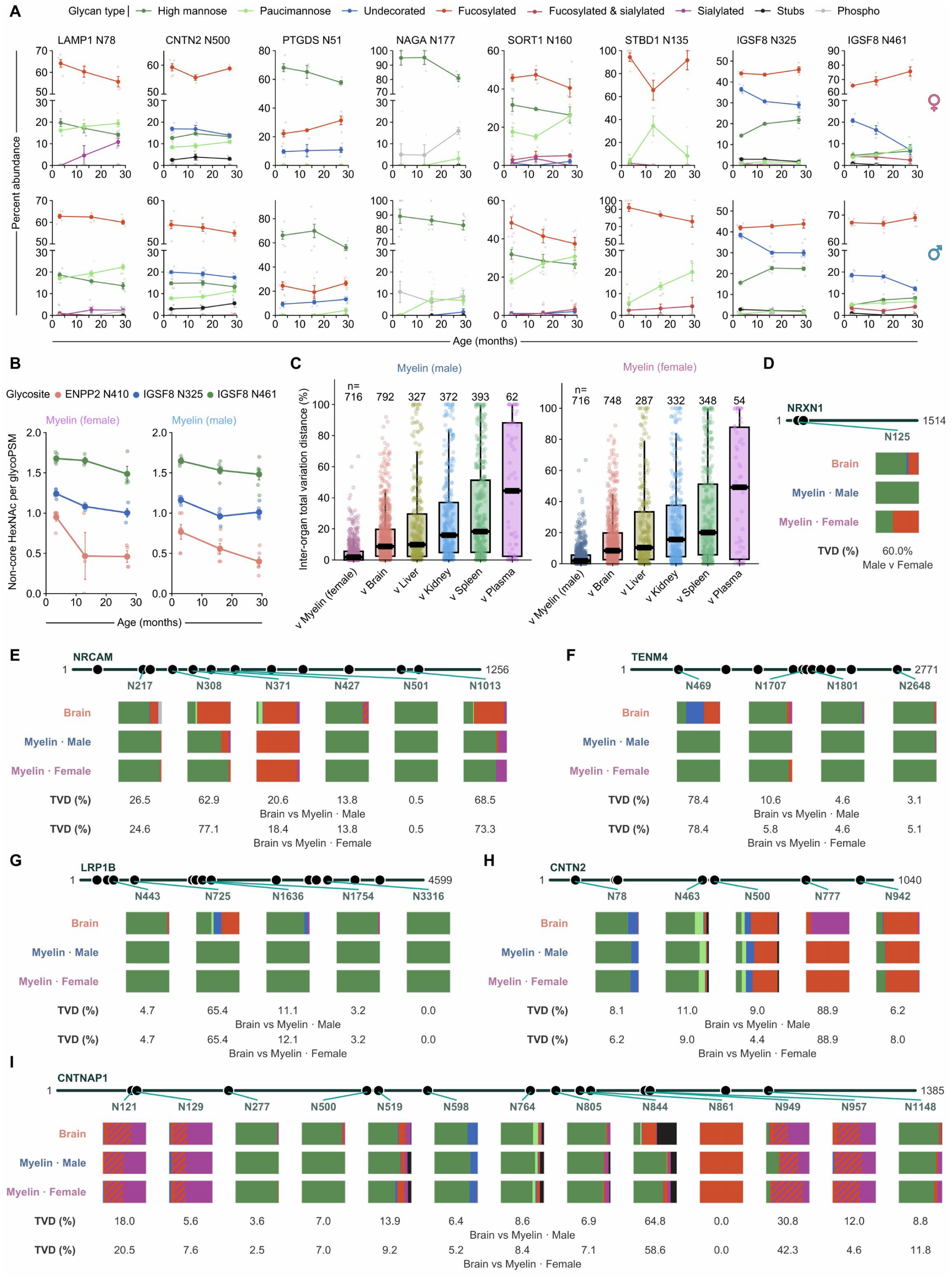
Site-resolved comparison of brain and myelin glycosylation. **(A)** Glycan type trajectories for selected glycosites in female and male myelin. Points are biological samples and lines and error bars show mean ± SEM. **(B)** Non-core HexNAc counts per glycoPSM across age at ENPP2 N410, IGSF8 N325, and IGSF8 N461. **(C)** Glycosite-level TVD comparing male and female myelin with one another and with atlas brain, liver, kidney, spleen, and plasma. Numbers denote shared eligible sites; box plots show the median, interquartile range, and whiskers extending to 1.5 × the interquartile range. **(D–I)** Pooled glycan type compositions for selected sites on NRXN1 **(D)**, NRCAM **(E)**, TENM4 **(F)**, LRP1B **(G)**, CNTN2 **(H)**, and CNTNAP1 **(I)** in atlas brain and male and female myelin. Values below each site are TVDs between male and female myelin **(D)** or brain and myelin **(E–I)**.

**Figure S20.**
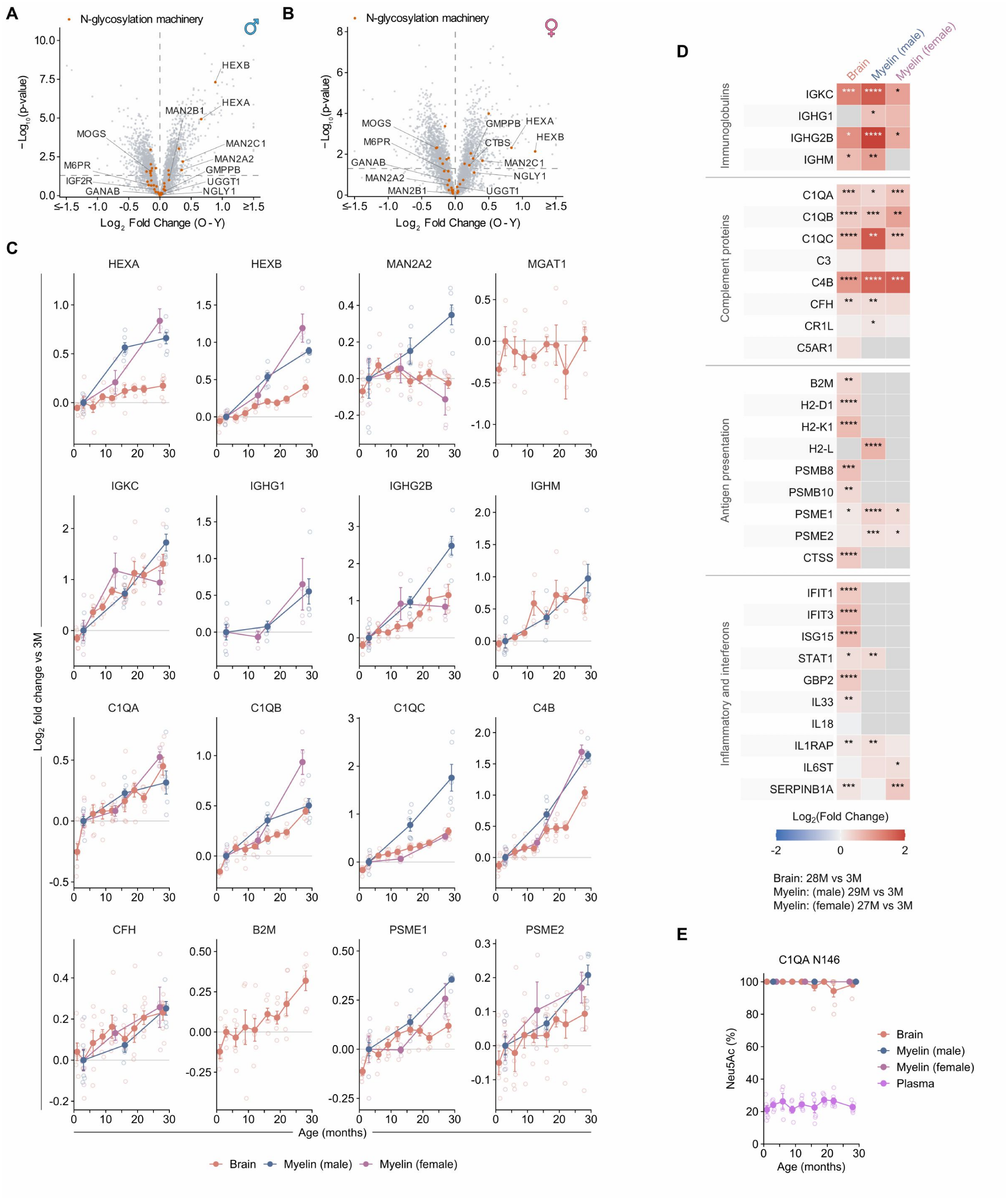
Age-associated changes in glycan-processing and immune proteins in the brain, purified myelin, and plasma. **(A and B)** Old-versus-young proteomic volcano plots for male myelin **(A)** and female myelin **(B)**. Orange points identify proteins annotated to N-glycosylation machinery; selected proteins are labeled. Horizontal dashed line indicates p = 0.05. **(C)** Age trajectories of selected glycan-processing enzymes, immunoglobulins, complement proteins, antigen-presentation proteins, and inflammatory proteins in whole brain and male and female myelin. Values are log_2_ fold change relative to 3 months; points are biological samples and lines and error bars show mean ± SEM. **(D)** Old-versus-young log_2_ fold changes for selected immune proteins in brain (28 versus 3 months), male myelin (29 versus 3 months), and female myelin (27 versus 3 months). Significance symbols denote p values from two-sided Welch t tests (*p < 0.05, **p < 0.01, ***p < 0.001, and ****p < 0.0001). **(E)** Percentage of sialic acid monosaccharides that are Neu5Ac at C1QA N146 across age in brain, male and female myelin, and plasma.

**Figure S21.**
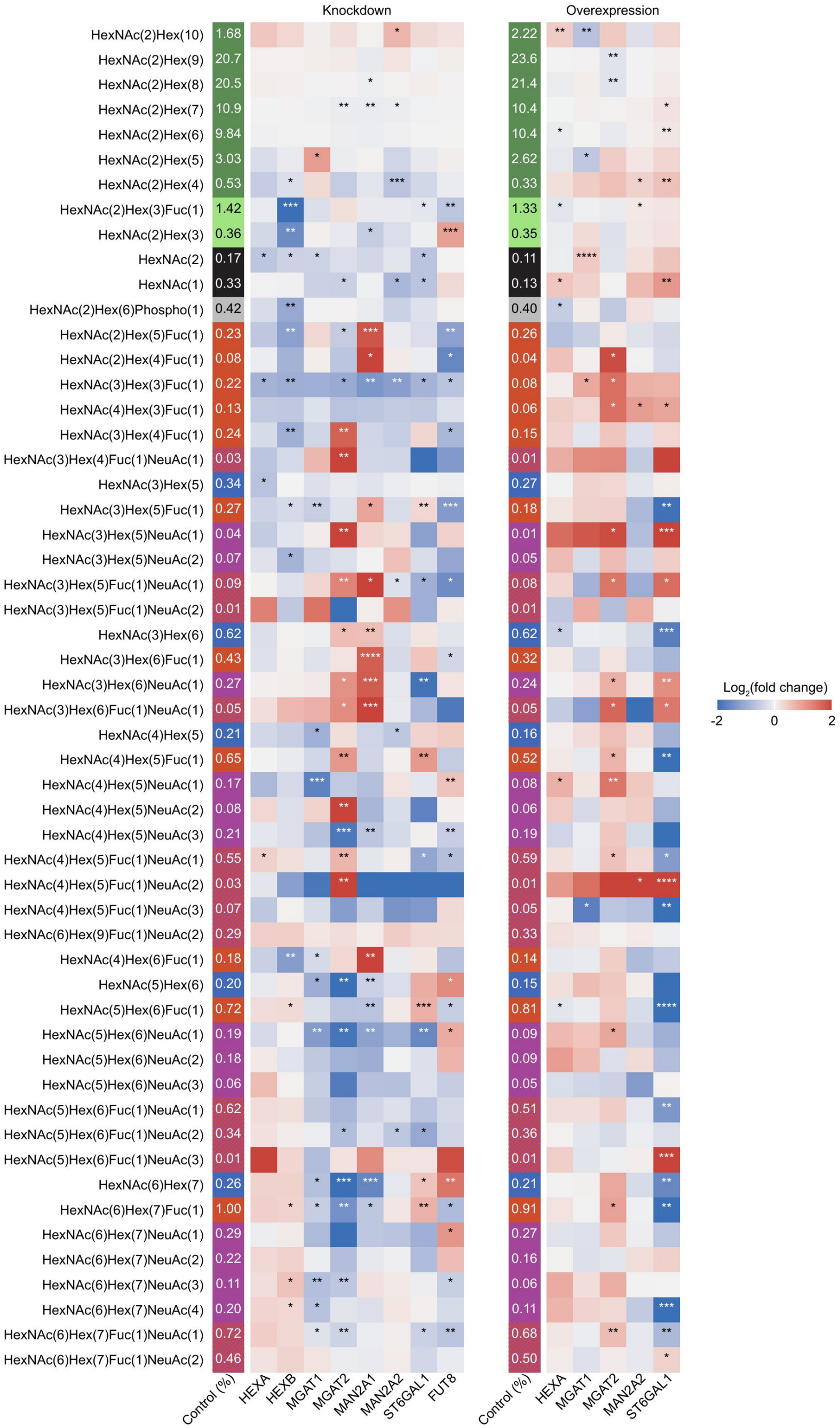
Exact-glycan responses to perturbation of N-glycan processing enzyme levels. Heatmaps show changes in the global percent abundance of individual exact glycans after knockdown or overexpression of the indicated glycan-processing enzymes in HOG cells. The first column of each block gives mean control cell abundance as a percentage of glycoPSMs; treatment cells show log_2_ fold change relative to the matched control. Rows are exact monosaccharide compositions, and control cells and each perturbation were analyzed in three biological replicates. Significance was assessed by two-sided Welch t tests on replicate-level abundances; symbols show p values (*p < 0.05, **p < 0.01, ***p < 0.001, ****p < 0.0001).

**Figure S22.**
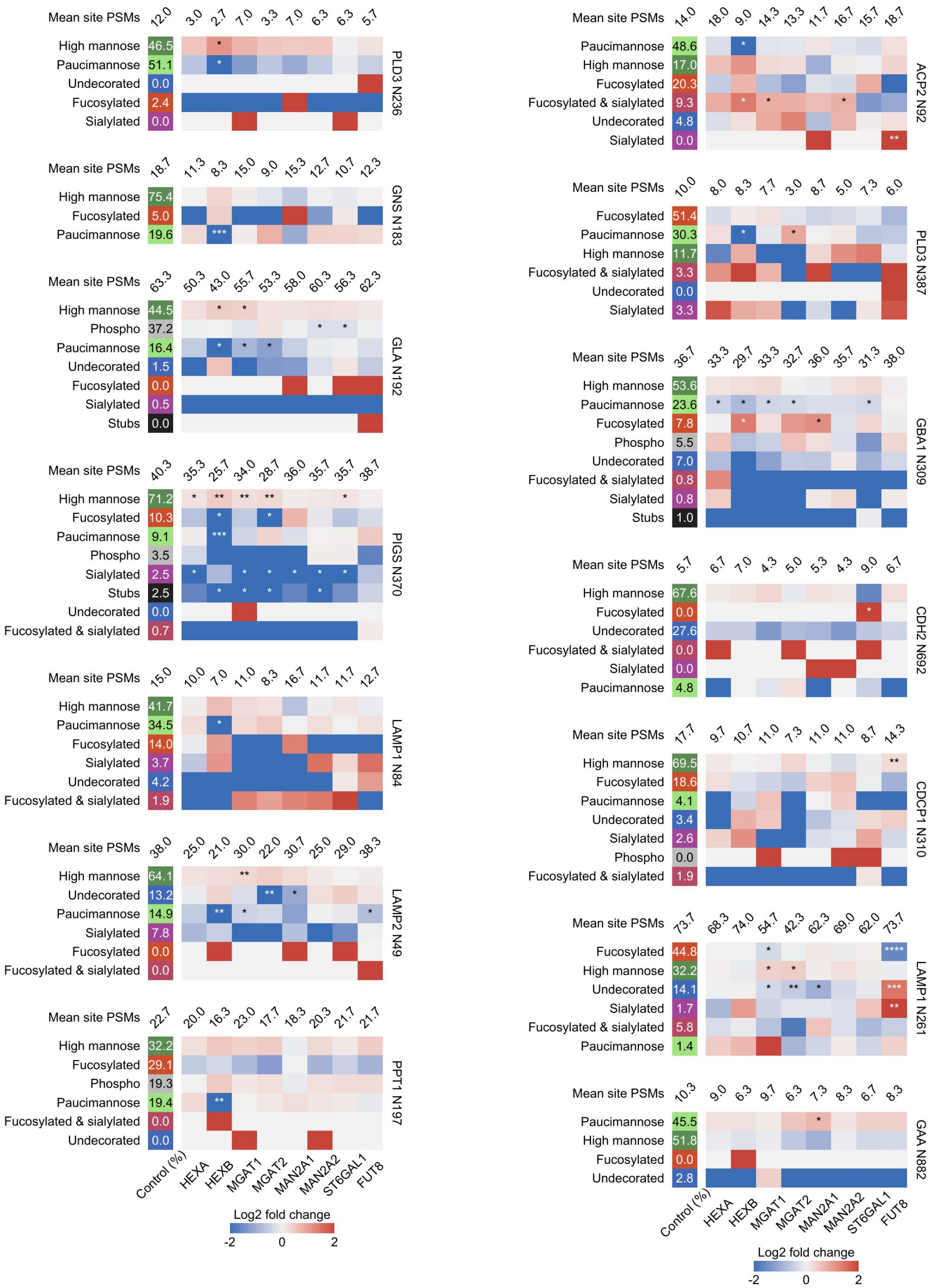
Site-specific glycan type responses to enzyme knockdown. Heatmaps show glycan type remodeling at selected HOG cell glycosites after knockdown of HEXA, HEXB, MGAT1, MGAT2, MAN2A1, MAN2A2, ST6GAL1, or FUT8. For each site, the left column gives the mean control cell within-site percent abundance of each detected glycan type. Colored treatment cells show the log_2_ fold change in within-site percentage relative to the matched control cell group, and values above columns give mean total glycoPSMs at that site. Each condition comprised three biological replicates. Significance was assessed with two-sided Welch t tests on replicate-level within-site percentages; symbols show p values (*p < 0.05, **p < 0.01, ***p < 0.001, ****p < 0.0001).

**Figure S23.**
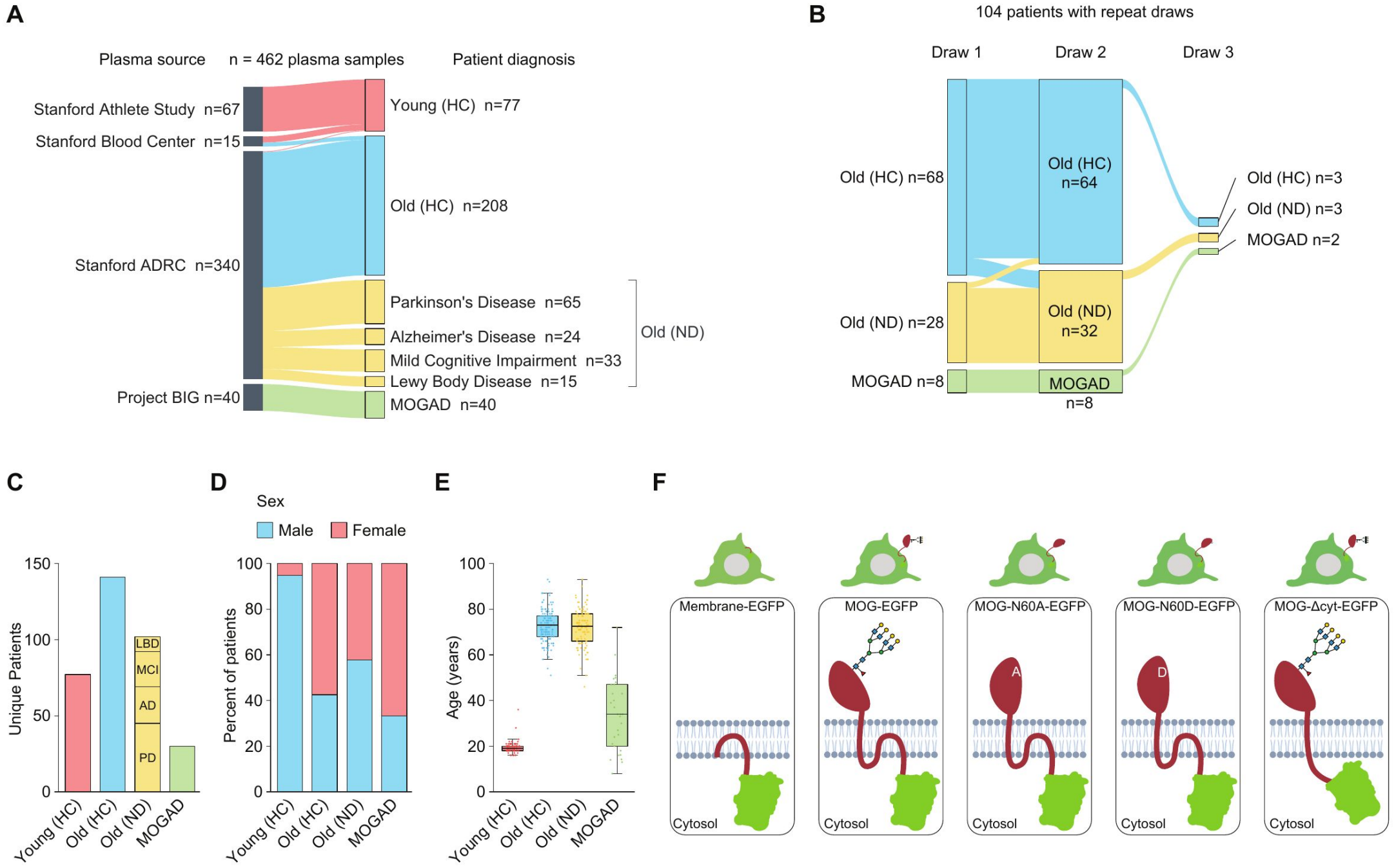
Participant plasma cohorts and MOG expression constructs. **(A)** Sources and diagnostic composition of the 462 plasma samples analyzed. Old neurological disease samples comprise Parkinson’s disease, Alzheimer’s disease, mild cognitive impairment, and Lewy body disease. **(B)** Diagnostic groups represented across repeated draws from 104 participants. Flow widths are proportional to sample counts, which are reported at each draw. **(C)** Unique participants in the Young (HC), Old (HC), Old (ND), and MOGAD groups; the Old (ND) bar is partitioned by diagnosis. **(D)** Sex distribution within each group. **(E)** Participant age distributions. Points denote participants; box plots show the median, interquartile range, and whiskers extending to 1.5 × the interquartile range. **(F)** Cell-surface expression constructs used for antibody-binding assays: Membrane-EGFP control, full-length MOG-EGFP, MOG-N60A-EGFP, MOG-N60D-EGFP, and MOG-Δcyt-EGFP. N60A and N60D prevent N-glycosylation at the MOG N60 sequon, and Δcyt removes the cytoplasmic tail and second transmembrane region.

**Figure S24.**
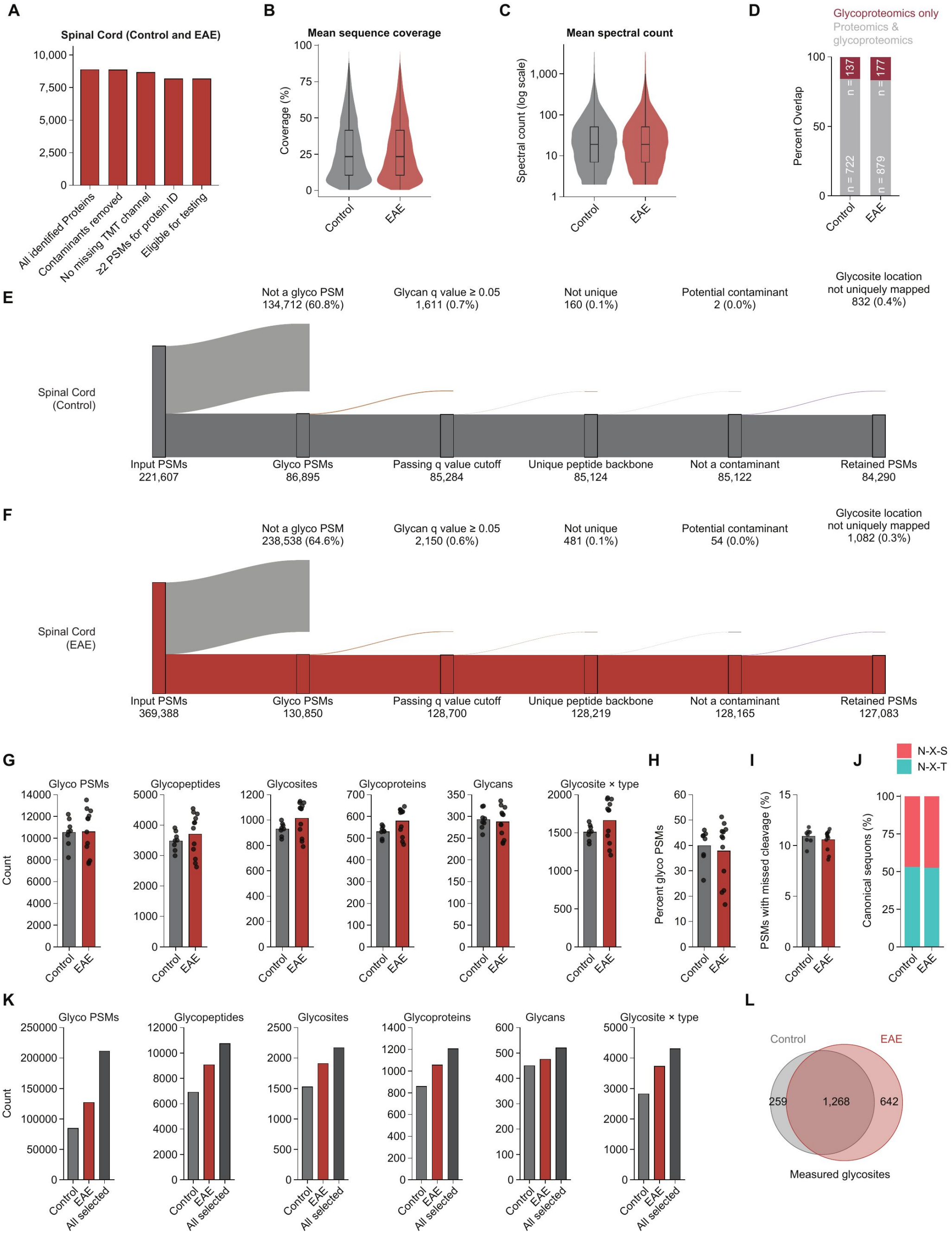
EAE spinal cord proteomics and glycoproteomics filtering and quality control. **(A)** Sequential filtering of spinal cord proteomics identifications through contaminant removal, complete reporter channel quantification, minimum PSM support, and replicate coverage. **(B and C)** Violin plots show distributions of protein sequence coverage **(B)** and spectral count **(C)** in control and EAE samples; embedded box plots show the median, interquartile range, and whiskers extending to 1.5 × the interquartile range. **(D)** Percentage of glycoproteins also quantified by total proteomics and the complementary percentage detected only by glycoproteomics. **(E and F)** Sequential filtering of control **(E)** and experimental autoimmune encephalomyelitis (EAE; **F**) spinal cord glycoPSMs through glycan q value < 0.05, unique peptide backbones, contaminant removal, and unambiguous glycosite mapping. Branch labels give retained or excluded PSM counts and percentages. **(G)** Numbers of glycoPSMs, glycopeptides, glycosites, glycoproteins, exact glycans, and glycosite–glycan type combinations per sample. **(H)** Percentage of filtered PSMs assigned to glycopeptides. **(I)** Percentage of PSMs with at least one missed cleavage. **(J)** Fraction of localized sites in N-X-S or N-X-T sequons. **(K)** Aggregate numbers of identified entities in control, EAE, and the combined dataset. **(L)** Overlap of glycosites detected in control and EAE. Points in **(G–J)** represent biological samples (control, n = 8; EAE, n = 12); bars denote means and error bars denote SEM.

**Figure S25.**
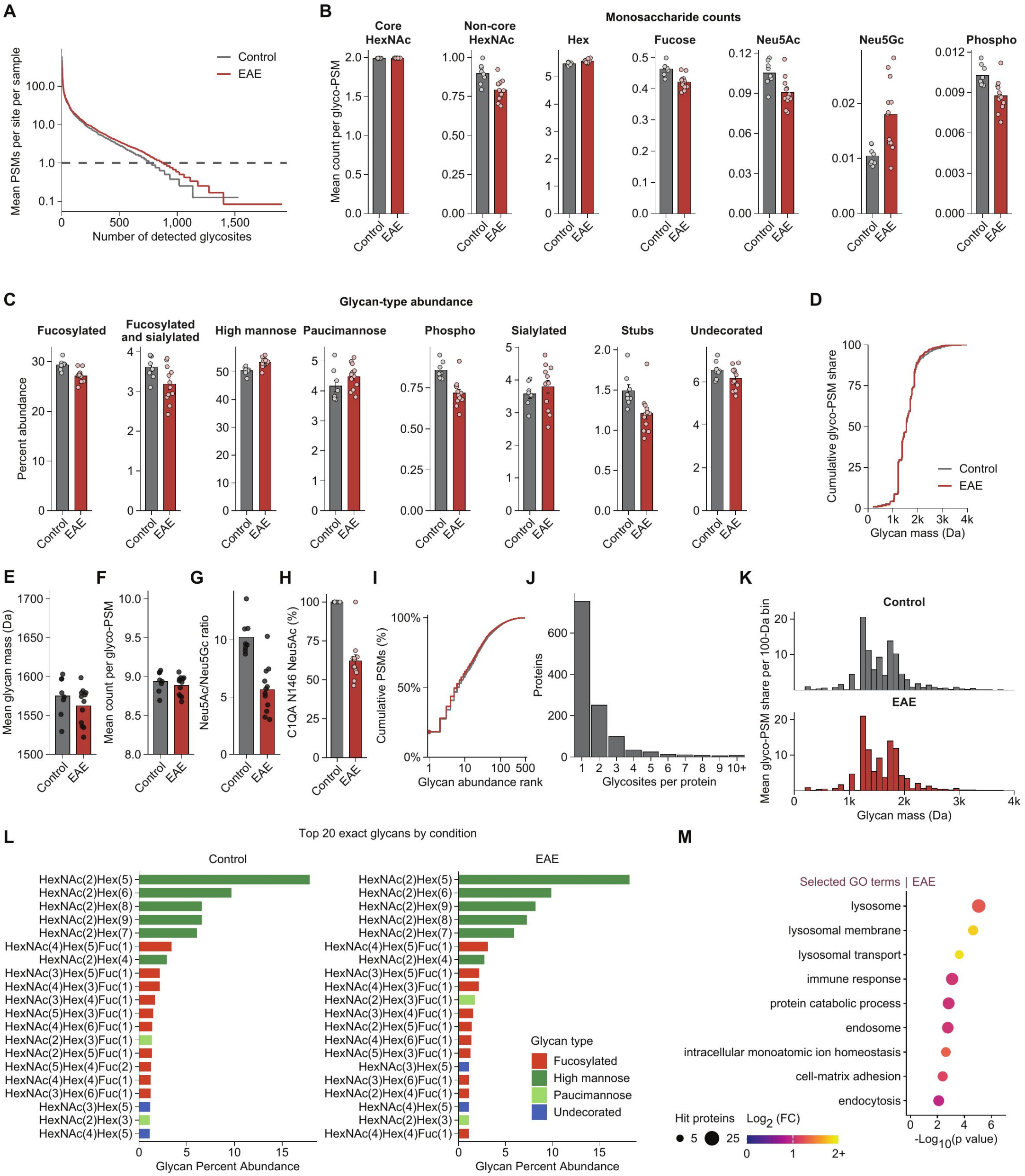
Global glycan properties and functional enrichment in EAE spinal cord. **(A)** Glycosites ranked by mean glycoPSM depth per sample; the dashed line marks one mean glycoPSM per sample. **(B)** PSM-weighted monosaccharide counts per glycoPSM. **(C)** Mean within-sample glycan type percent abundance. **(D)** Cumulative glycoPSM share as a function of glycan mass. **(E)** PSM-weighted mean glycan mass. **(F)** Mean total monosaccharide count per glycoPSM. **(G)** Neu5Ac/Neu5Gc monosaccharide ratio. **(H)** Percentage of sialic acid monosaccharides that are Neu5Ac at C1QA N146 in control and EAE spinal cord. **(I)** Cumulative glycoPSM abundance of exact N-glycans ranked in control and EAE spinal cord. **(J)** Numbers of glycosites per glycoprotein. **(K)** Mean glycoPSM share in 100-Da glycan mass bins. **(L)** Twenty most abundant exact glycans in control and EAE, calculated from pooled glycoPSMs. In **(B), (C)**, and **(E–H)**, points are biological samples and bars denote means ± SEM. **(M)** Selected GO terms enriched among proteins carrying glycosites with significant composition changes in EAE-versus-control.

